# Uterocalin – a protein uniquely specialised to support mobile encapsulated embryos of horses, rhinos and tapirs

**DOI:** 10.64898/2026.09.17.752455

**Authors:** Malcolm W. Kennedy

## Abstract

Equids are considered unique in having encapsulated embryos that are actively moved between the uteri of mares during the first two and three weeks of pregnancy and do not form a placenta until about 40 days, the longest pre-placentation period known amongst Eutheria. Their embryos are likely supplied through the glycoprotein capsule with selected nutrients by uterocalin, a carrier protein secreted by maternal endometrial glands. Uterocalin binds polyunsaturated fatty acids and retinol (Vitamin A), concentrates in the capsule, and is internalised by the embryo. Here it is shown that the structures of equid uterocalin comprise an excess of metabolism-related essential amino acids, and structural specialisations potentially involved in stabilising the capsule matrix until emergence and engagement with a receptor on the trophoblast surface. The genomes of other perissodactyls - rhinos and tapirs – encode uterocalins with amino acid sequences, excesses of essential amino acids, and predicted structural properties, all closely similar to horse uterocalin. These characteristics are unique to the perissodactyl uterocalins and are not found in similar proteins of other mammal groups. Hence, rhinos and tapirs probably have equid-like reproductive specialisations retained since before the Cretaceous-Paleogene mass extinction and that uterocalin is key to both embryo nutrition and capsule formation.

## 1. Introduction

How mothers support their embryos varies dramatically across the viviparous mammals. Within the Eutherian (“placental”) mammals placentation processes and placental types are remarkably diverse, even within the four major clades. An extreme example across the Eutheria would be the difference between humans and horses. In the former a blastocyst attaches to uterine epithelium within a few days of fertilisation and invades the endometrium to be surrounded by, and in direct cell:cell contact with, maternal tissue and nourished by secretions prior to forming a haemochorial placenta in which the trophoblast is ultimately bathed in maternal blood (1–3). In contrast, horses have the longest pre-placentation period (without embryonic diapause) known amongst eutherians in which the conceptus exists within an acellular glycoprotein/proteoglycan capsule that is moved along and between the twin uterine horns by peristaltic action during the second and third weeks of pregnancy (1, 4–6). Movement of the conceptus around the uteri is essential to the maternal recognition of pregnancy (5–8). It then emerges from the capsule around twenty days, attaches to the endometrium (“fixation”), and goes on to form a diffuse, villous, epitheliochorial placenta at around 40 days, in which the maternal epithelium is not deeply penetrated (1, 2, 4). During the mobile encapsulated stage the conceptus accumulates fluid to maintain sufficient internal turgor to ensure a spherical state (9–12). The acellular glycoprotein capsule is therefore essential for the survival against uterine contractions of a spherical, mobile conceptus that would otherwise be bounded by only a fragile trophoblast cell layer.

The possession of an enclosing capsule means that there can be no direct cell:cell contact between embryo and maternal tissues for the delivery of nutrients and the conceptus will consequently be dependent on uterine glandular secretions (histotrophe) that can traverse the capsule. The uterus upregulates expression of amino acid and glucose transporters (13, 14), for example, to provide such soluble nutrients, but delivery of poorly soluble and chemically sensitive lipids that, unless packaged, could simply bind back to maternal membranes is conceptually more difficult. Delivery of lipids to the conceptus is considered to rely on a protein of the lipocalin family that binds lipids such as polyunsaturated essential fatty acids and retinol (vitamin A). This protein, uterocalin, secreted by the mare’s uterine glands, preferentially associates with the capsule and is internalised by the embryo, thereby delivering both essential lipids and its own amino acids (15–19).

Horses are members of the Perissodactyla (odd-toed ungulates; equids, rhinos, tapirs) that are currently estimated to have separated from the Ferae (Carnivora and Philodota (pangolins)), and prior to that the Ferungulata (Ferae and Artiodactyls (sometimes now referred to as Cetartiodactyla; even-toed ungulates, cetaceans, hippopotamus)), around 10 million years before the Cretaceous-Paleogene mass extinction event (20–22). The Equidae subsequently split within the Perissodactyla around 7 Myrs in the Paleocene, after the extinction, with the ancestors of the Rhinoceridae and Tapiridae separating a few million years thereafter, around the Paleocene-Eocene junction (20–22). Within the large Ferungulata clade, and indeed even all of the Eutheria, only the equids are known to have such prolonged pre-placentation, mobile, encapsulated embryos, a reproductive feature regarded as unique to them. The question tackled here is whether this is true and whether it is instead a long-retained characteristic of perissodactyls as a whole?

But, how can we tell? There is no readily accessible comparable information on early conceptuses and placentation to that in equids on any species of tapir or rhino. Because of practicalities and conservation concerns, the latter two are considerably less available than equids for investigation in terms of accessibility and basic research. One way to address the question is to seek encoded within the genomes of these species a characteristic that is seemingly unique to the distinctive form of early conceptus support found in equines. The uterocalin protein of horses provides such a clue. The lipocalin family of proteins to which uterocalin belongs has a wide range of functions (23–27) ranging from immune protection, enzymic modification of lipids, pheromone distribution, and vitamin transfer, uterocalin itself being a lipid carrier (15). Lipids delivered to the embryo would contribute to energy metabolism and membrane construction, but can also be precursors of signalling lipids acting as morphogens and in tissue (re)generation and include those central to bilateral symmetry (28–33). Of particular importance to encapsulated embryos of horses would be the provision of fatty acid precursors to secreted signalling compounds such as prostaglandins that induce myometrial peristalsis and contribute to maternal recognition of pregnancy in this species (8, 34–36).

As shown here, the uterocalin of equids has compositional and predicted structural characters that distinguish it from other lipocalins, including putative ancestral proteins. And that both rhinos and tapirs have a protein encoded in their genomes that have the same, distinctive characters. The corollary of this is that rhinos and tapirs also have long pre-placentation, mobile, encapsulated conceptuses like those of equids. While there is accumulating direct evidence for this in rhinos (37–41), there is none yet for tapirs. But, such is the similarity of their uterocalin proteins to that of equids and rhinos, and their phylogenetic closeness (20–22, 42), it is likely that the same applies to them. It is therefore argued here that perissodactyls collectively exhibit a highly unusual reproductive characteristic amongst eutherians that has been retained since their separation from major mammalian clades before the Cretaceous-Paleogene mass extinction event. And that the uterocalin protein is uniquely specialised to support their unusual reproductive mode, not only for the delivery of nutrients via the capsule, but also as a crucial component in the stability of the capsule itself.

## 2. Materials and Methods

### (a) Protein amino acid sequences

All but one of the protein sequences used were obtained from the National Center for Biotechnology Information (https://www.ncbi.nlm.nih.gov/) database, and similar protein sequences sought using its BLAST procedure set for protein-protein sequence searches. All the protein sequences dealt with here are given in the supplementary material. The cleavable N-terminal secretory signal sequences (usually comprising the first 18 amino acids) were predicted using Signalp 6.0 (https://services.healthtech.dtu.dk/services/SignalP-6.0/). BLAST searches were only carried out using the predicted mature, secreted protein with the cleavable secretory signal peptide removed, and only these were used for amino acid content and other calculations. A search using the horse, *Equus caballus,* uterocalin sequence found near-identical sequences in several equid species (listed in the supplementary material), but only the *E. caballus* sequence is used here; the uterocalin-like protein sequences from *Equus asinus, Equus quagga,* and *Equus przewalskii* are either identical to that of the horse or have with only one amino acid substitution. The uterocalin-like protein in the database from *Diceros bicornis minor* (South-central black rhinoceros) is longer than the 180 amino acid length of uterocalins and has been labelled as an epididymal-specific lipocalin-9. This is likely to be a category error in that lipocalin-9s do not have the unusual characteristics as we discuss for uterocalins, the first 180 amino acids align closely with equine uterocalin, and the amino acid stretch beyond gives no similarities in BLAST searches other than to self, so, only the first 180 amino acids were used. A similar extended sequence pertains to the predicted protein from *Ceratotherium simum simum* (southern white rhinoceros), and in which the final five amino acids of the uterocalin-like sequence also appear to be in error, so only the first 175 positions of this are used in the amino acid composition analysis. NCBI or other protein databases did not provide the *Tapirus indicus* protein, which was instead found in the *T. indicus* genome database via https://www.dnazoo.org/assemblies/tapirus_indicus and link to https://dnazoo.s3.wasabisys.com/index.html?prefix=Tapirus_indicus/. Other reference protein sequences used were as given below. Supplementary material table S1 gives a list of all of the proteins dealt with, their database accession codes, current descriptors, ProtParam calculations for amino acid content and predicted isoelectric point (pI), and how they were edited.

The term “uterocalin” has been applied to two different types of lipocalin family proteins. One is more commonly termed neutrophil gelatinase-associated lipocalin (NGAL), lipocalin2, or siderocalin, and was originally defined from mouse uteri but is now found systemically in response to disease. The other, the one dealt with here, is the protein secreted in abundance into the lumen of the pregnant uterus of mares, and associates with the glycoprotein capsule around the pre-placentation conceptus, although is expressed at low levels elsewhere. Several proteins that appear from BLAST similarity searches (see supplementary material) have been named as uterocalins (uterocalin being the name originally ascribed to the horse uterine protein (15)), presumably due to some similarity amongst lipocalin proteins that are otherwise highly sequence-diverse but may not have functional relationships, as illustrated in Results.

The term conceptus refers to the embryo and its associated structure, here, the capsule.

### (b) Protein amino acid sequence and composition analysis

Amino acid sequence alignments were made using MultAlin (http://multalin.toulouse.inra.fr/multalin/) set for the Blosum62 substitution matrix. Amino acid compositions and predicted isoelectric points were calculated using ProtParam (expasy.org/protparam/).

OriginLab ORIGIN 21 was used for dot plot graphing and regression analysis with the intercept set to zero. This used the amino acid percent content of only the mature part of the sequence (with the secretory signal peptide removed) graphed against the Reviewed SwissProt statistics for average amino acid compositions across the entire protein database (https://www.uniprot.org/uniprotkb/statistics). Hierarchical clustering and Partial Least Squares Discriminant analysis (PLS-DA; which also provides the important features (VIP) analysis) was carried out with MetaboAnalyst 6.0 (https://www.metaboanalyst.ca/) using the number of each amino acid in each protein as factor concentrations, without normalisation.

The essential amino acids that cannot be synthesised by mammals and must be supplied from diet or by mothers to their embryos and are histidine (His; H), isoleucine (Ile, I), leucine (Leu, L), lysine (Lys, K), methionine (Met, M), phenylalanine (Phe, F), threonine (Thr, T), tryptophan (Try, W), and valine (Val, V). A few others, notably arginine (Arg, R), are considered a semi-essential or conditional essential amino acids that are important under some metabolic conditions or embryonic stage.

### (c) Molecular graphics

The protein molecular structure of equine uterocalin was created in Pymol (https://pymol.org/) using molecular coordinates from the AlphaFold prediction through UniProtKB entry for *Equus caballus* (Horse) uterocalin (Q28388 · UCAL_HORSE), with link to AlphaFold Protein Structure Database entry AF-Q28388-F1-v6. In that prediction the secretory peptide is not part of the structure of the mature secreted form and has not been included in the molecular structures shown. This is a computer-generated model rather than empirical and as such cannot be regarded as definitive, although given that there are a very large number of empirically-obtained structures of lipocalins to provide templates, a good degree of confidence is merited. Geometry checks carried out using the Savesv6.1 online facility at https://saves.mbi.ucla.edu/ indicate a favourable Ramachandran plot - residues in favoured regions, 81.2%; residues in allowed regions, 17.5%; residues in generously allowed regions, 1.2%; residues in disallowed regions, 0.0%). The culled cavity prediction was set for four solvent radii using Pymol.

## 3. Results

### (a) Amino acid sequence similarities between horse, rhinoceros and tapir uterocalins

Alignments of the amino acid sequences of horse uterocalin with similar proteins encoded in the genomes of two species of rhinoceros and a tapir show similarities sufficient to indicate true orthologues (figure 1). The cleavable secretory sequence leader peptides that will be lost prior to secretion (amino acids 1 to 18) are closely similar. Also, where there are substitutions between the sequences, they are frequently one essential amino acid for another – e.g. Met for Lys, Ile for Lys, Phe for Leu, Phe for Val, Met for Val, or amino acids that similar in size or charge or size, such as Lys for Arg, Gly for Ala.

**Figure 1.**
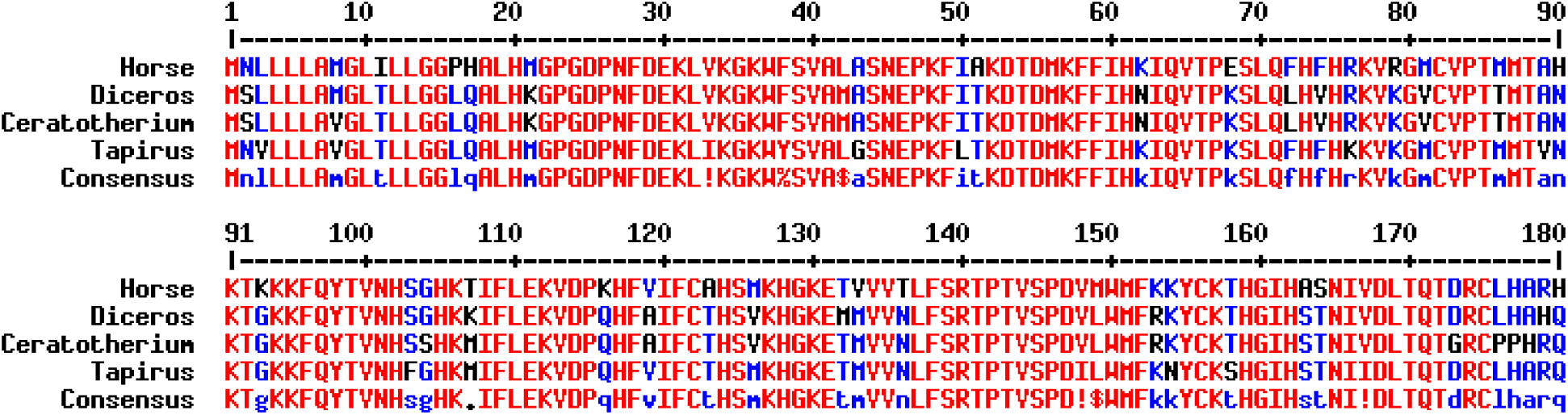
Amino acid sequence alignment of the uterocalin proteins of horses, rhinos, and a tapir. The first 18 amino acids comprise the cleavable secretory leader peptide that would be removed during secretion by the synthesising cell. Single letter codes used for the amino acids.

### (b) Unusual enrichment in essential amino acids

When the proportions of amino acids present in any of the perissodactyl uterocalins (excluding the secretory signal peptides) are plotted against the entire SwissProt database for the average amino acid content covering all protein sequence entries, the four perissodactyl uterocalins are highly enriched in five or six of the nine essential amino acids (figure 2A,B,C,D). One of the exceptions, leucine, occurs at unusually low levels for an amino acid that that it is overall the most abundant in proteins.

**Figure 2.**
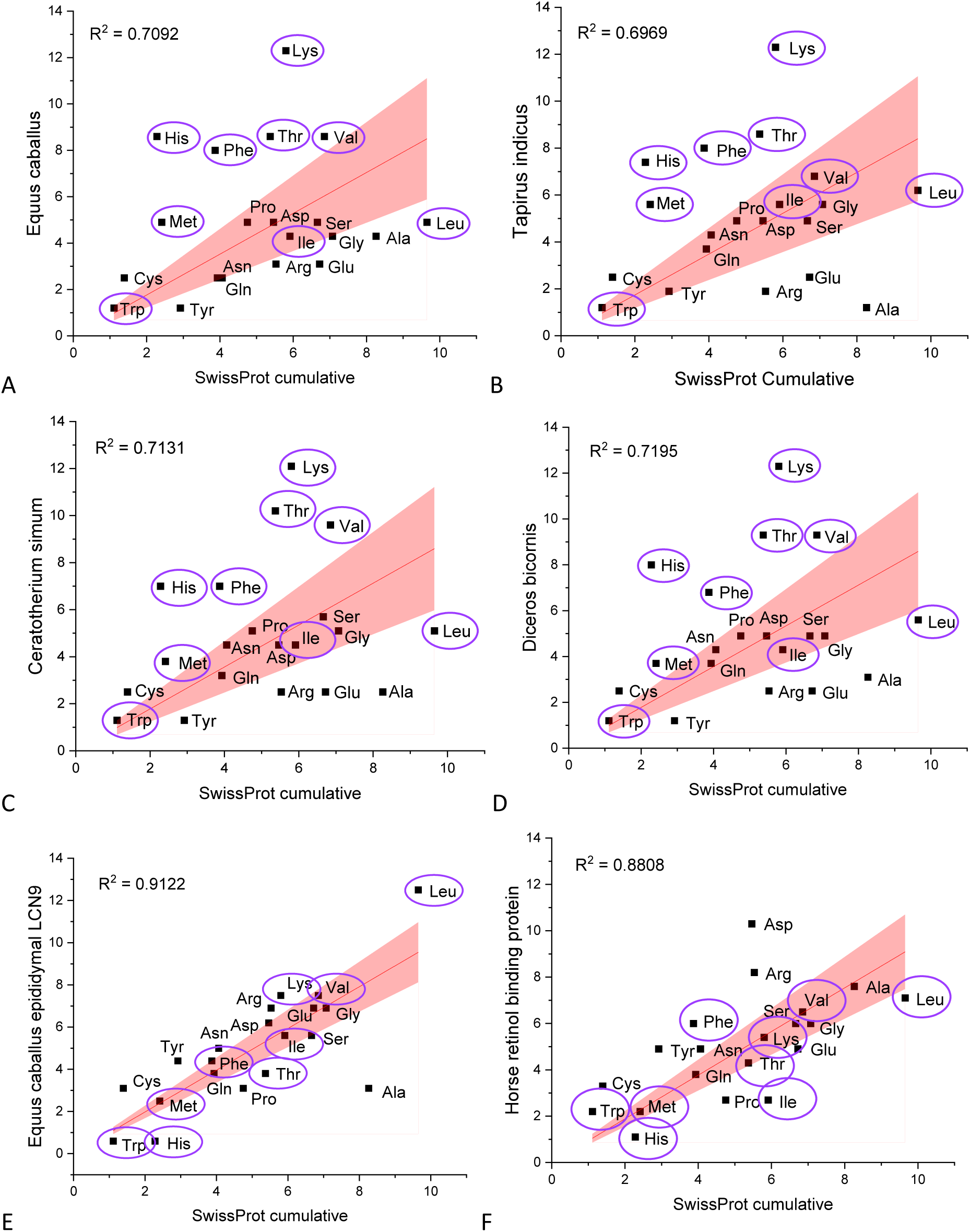
Comparison of the amino acid content of uterocalin proteins from horse, two species of rhinoceros, and the tapir. Each graph shows the percent content of each amino acid in the proteins excluding the secretory leader peptide versus the SwissProt database average for all protein entries. (A, B, C, D) The perissodactyl uterocalin proteins all exhibit excesses of selected members of the nine essential amino acids and unusually low proportions of leucines. (E, F) Similar analysis for two other lipocalins from the horse that do not exhibit the unusual scatter of essential amino acids. The linear regression fitted lines are set for intercepts to zero, and the coloured areas are the 95% confidence intervals.

For comparison, two other lipocalin proteins of horses, one associated with reproductive tissues (lipocalin 9; epididymus-specifc lipocalin (LCN9)), and a carrier of retinol in plasma (retinol binding protein), exhibit unremarkable proportions of essential amino acids (figure 2E and 2F).

Other proteins that arise close to the perissodactyl uterocalins in BLAST searches and have been labelled as “uterocalin” in databases because of sequence similarity to equine uterocalin from a range of species (exemplified here with two camel species, a pig, a feline, and a cetacean) all show no strong bias in essential amino acid content (figure 3A,B,C,D,E), and their contents of leucine are high or within norms. The overall scatter of amino acids for the perissodactyl uterocalins is greater than for all the others, as emphasised by the lower R^2^ regression values for the former and higher, closer fitting R^2^ values and tighter confidence intervals than for the others. The excess of essential amino acids in the horse, rhino and tapir proteins is further emphasised is a simple plot of the proportions of each protein comprising essential amino acids (figure 3F), this being apparent despite the depletion in leucines.

**Figure 3.**
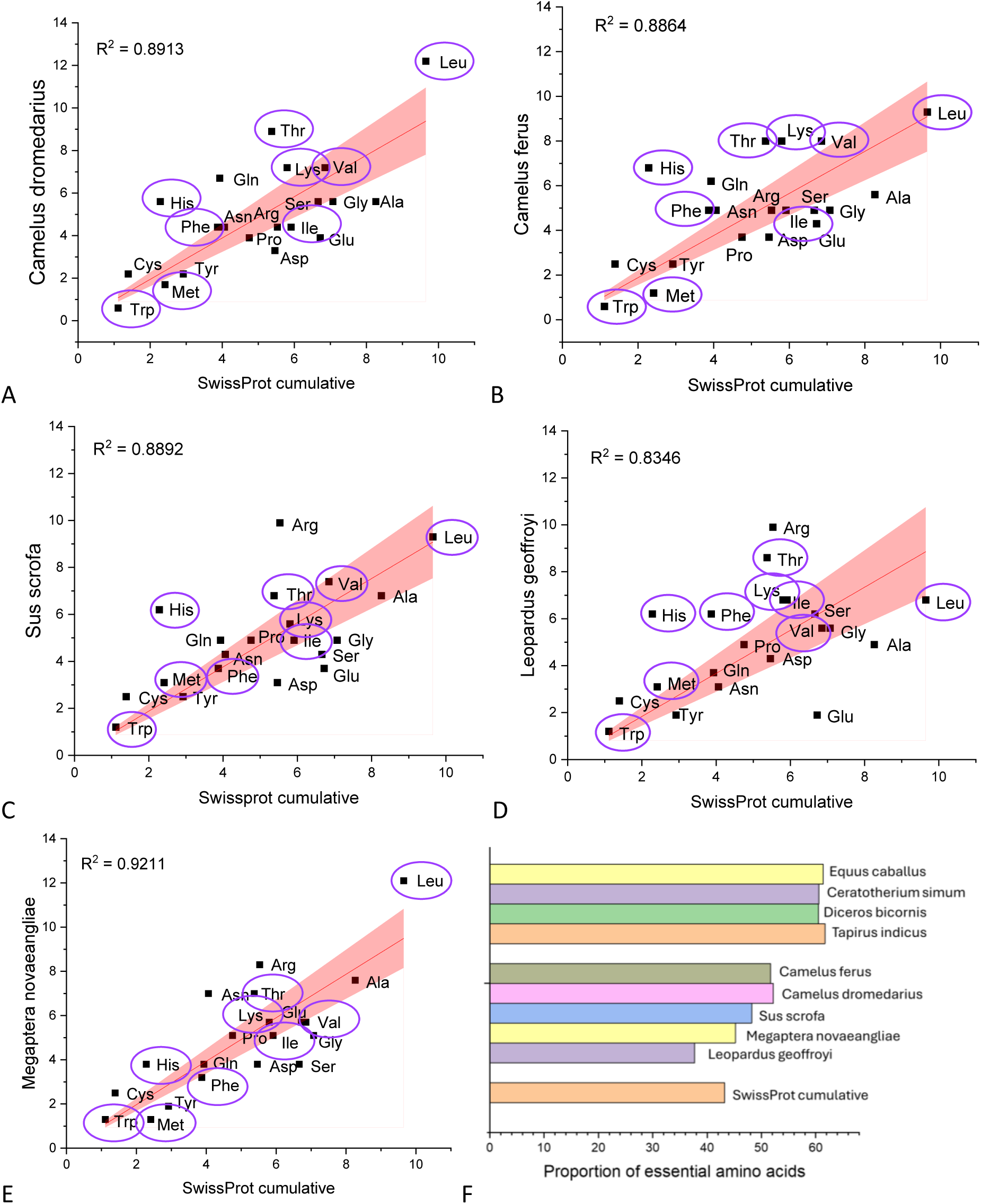
Uterocalin-like proteins from non-perissodactyls. (A to E) Comparison of the amino acid compositions of proteins that appear close to equine uterocalin by sequence similarity BLAST searches from *Camelus, Leopardus, Sus,* and *Megaptera* do not show the unusual distribution of essential amino acids. The regression R^2^ values for all six of these proteins are higher than in the perissodactyl uterocalins in figure 2, and the 95% confidence intervals are tighter, indicative of a closer adhesion to protein amino acid content norms. (F) A plot of the cumulative essential amino acid content of the perissodactylid uterocalins (upper four bars) versus the uterocalin-like proteins represented in A to E, and the SwissProt average. The perissodactyl proteins exceed the others in essential amino acid content despite the relative depletion of leucine.

When all the perissodactyl uterocalins and uterocalin-like proteins from a range of species within the Ferungulata are compared by hierarchical clustering analysis, a heatmap again emphasises the distinctive character of the perissodactyl uterocalins in that they group tightly and clearly separate from the uterocalin-like proteins (figure 4A). And this tight, coherent grouping clusters around essential amino acids, despite the straying of leucine, isoleucine, and tryptophan. An identical analysis using these proteins plus a range of unrelated lipocalins, including those associated with reproductive tissues, further emphasises the distinctiveness of the perissodactyl lipocalins (supplementary material figure S1).

**Figure 4.**
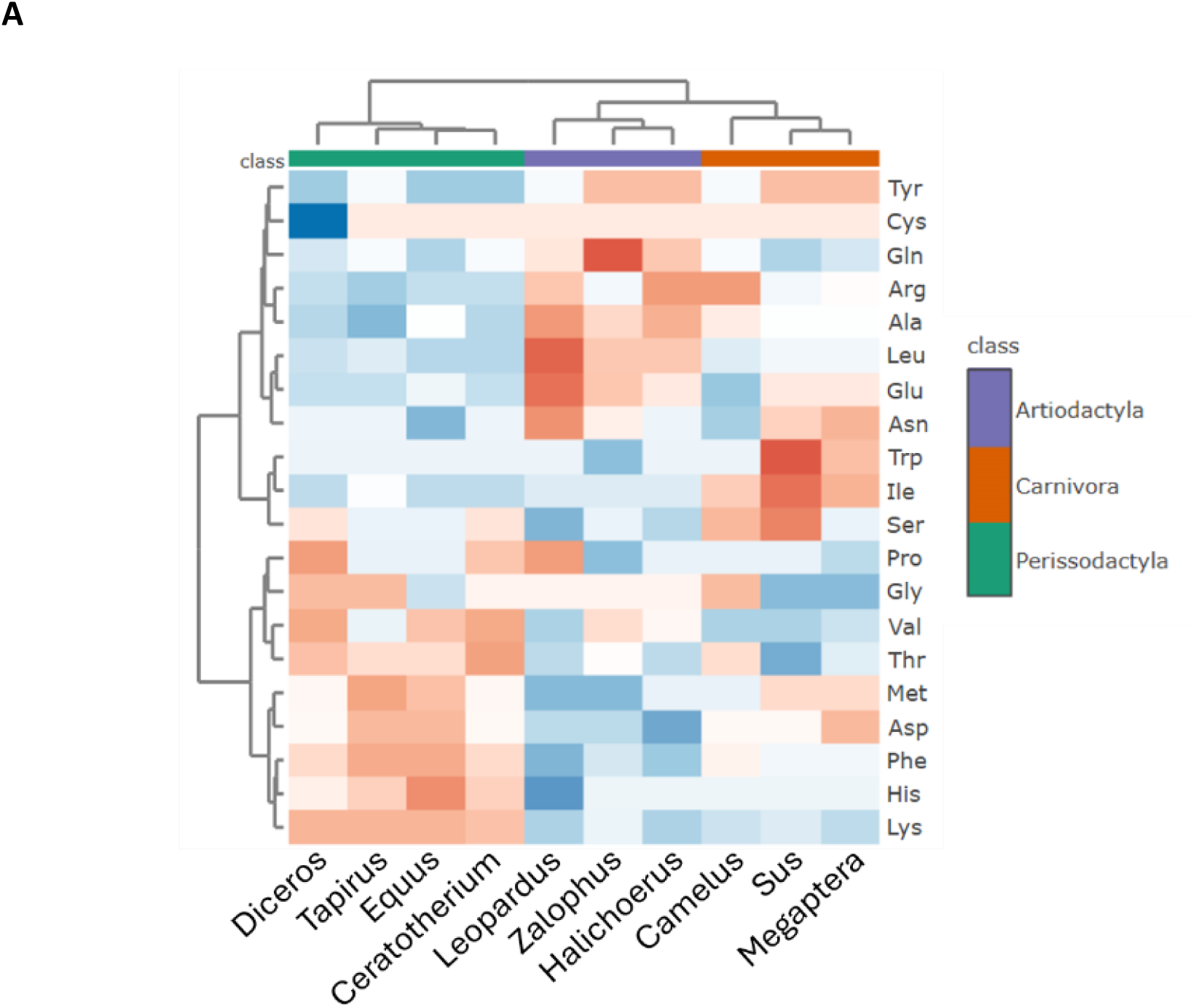
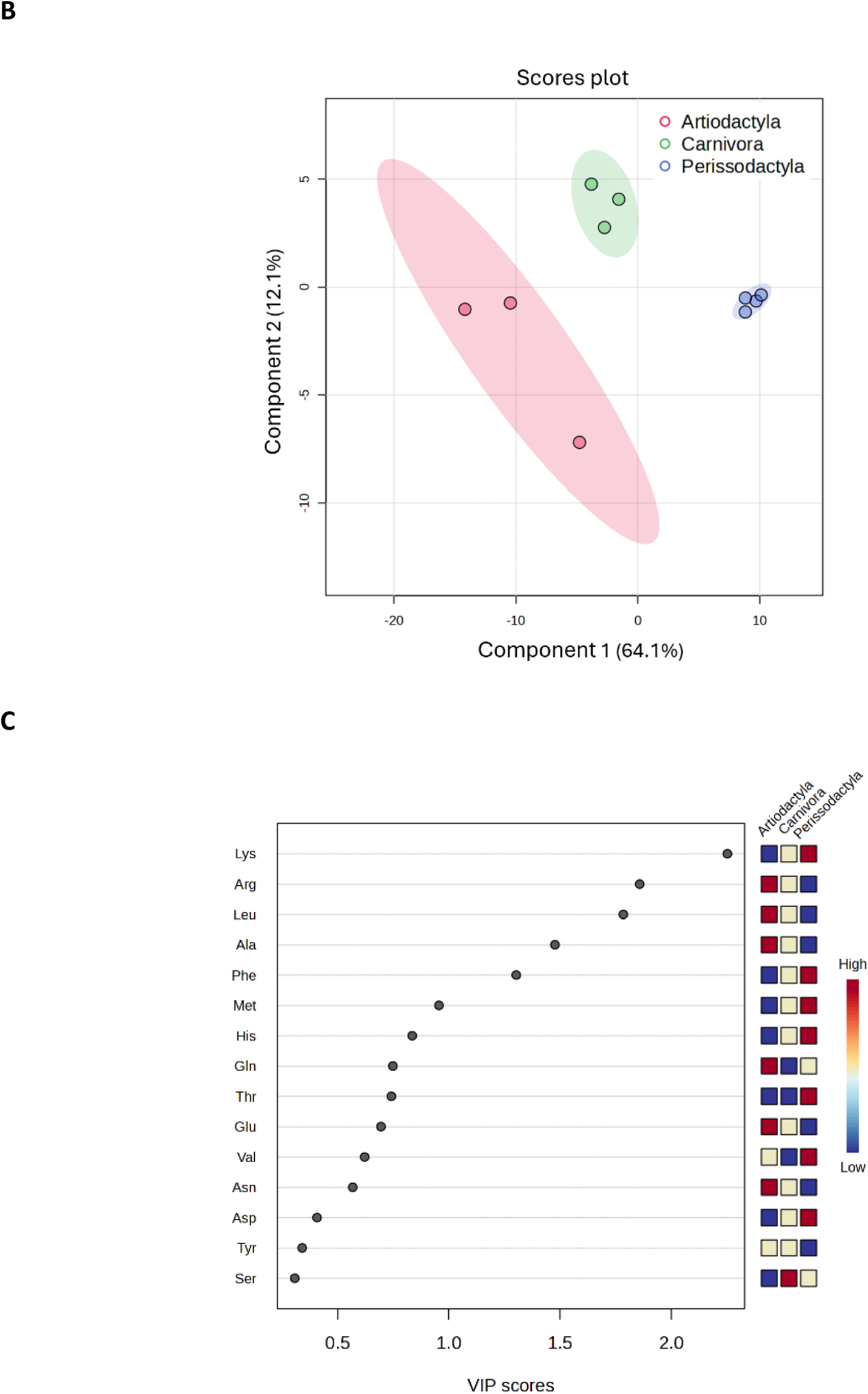
Hierarchical clustering and principal component analysis of the four Perissodactyla uterocalins against the uterocalin-like proteins from other Ferungulata. (A) Hierarchical cluster analysis of the Perissodactyl (horse, rhinoceros, tapir) uterocalins reveals distinct grouping away from the Ferungulata (here split into Artiodactyla and Carnivora) uterocalin-like proteins driven by their composition of the essential amino acids His, Thr, Lys, Met, Val, and Phe, the exceptions being Leu, Ile, and Trp. The proteins from camels (*Camelus*) are closest to the perissodactyl uterocalins in sequence similarity BLAST searches but do not cluster with them. (B) Principal component analysis carried out by Partial Least Squares Discriminant Analysis (PLS-DA) additionally emphasises the distinctiveness of the perissodactyl uterocalins. (C) Important features analysis (VIP) showing the contents of Lys, Arg, Leu and Phe of the essential amino acids, and Arg, a conditionally essential amino acid, high or low, comprise the overall distinguishing features of the perissodactyl proteins. The analyses were based on the number of each amino acid in each protein excluding the secretory signal peptide and used the original data without normalisation.

Principal component analysis of all the uterocalin-like sequences curated here similarly shows the distinctiveness of the perissodactyl proteins (figure 4B). And VIP analysis identifies lysine, leucine, arginine and phenylalanine contents as particularly important discriminators (figure 4C) between the perissodactyl uterocalins and the others.

So, by amino acid content of essential amino acids alone the perissodactyl uterocalins are distinctive within the larger family of lipocalins, including those that appear close to them in amino acid sequence similarity searches. This effect is apparent even though the normally most abundant essential amino acid in proteins, leucine, is depleted.

### (e) Regional specialisation in perissodactyl uterocalins from a putative ancestral common ancestor

Aligning amino acid sequences of uterocalin-like proteins that are closest to the perissodactyl uterocalins arising from sequence similarity searches reveals clear similarities but also distinct short regions of divergence (figure 5A). The three regions of clearest difference, positions 48 to 59, 99 to107, and 123 to 132, are mapped these onto a predicted structure of horse uterocalin reveals them to be on loops immediately adjacent to the opening to the internal cavity, or portal, of the protein (figure 5B,C). Notably, these regions show no such differences between the horse, rhinoceros and tapir sequences (figure 1). Regions of lesser differences at positions 144 to 154, and 157 to 164, map to the external helix.

**Figure 5.**
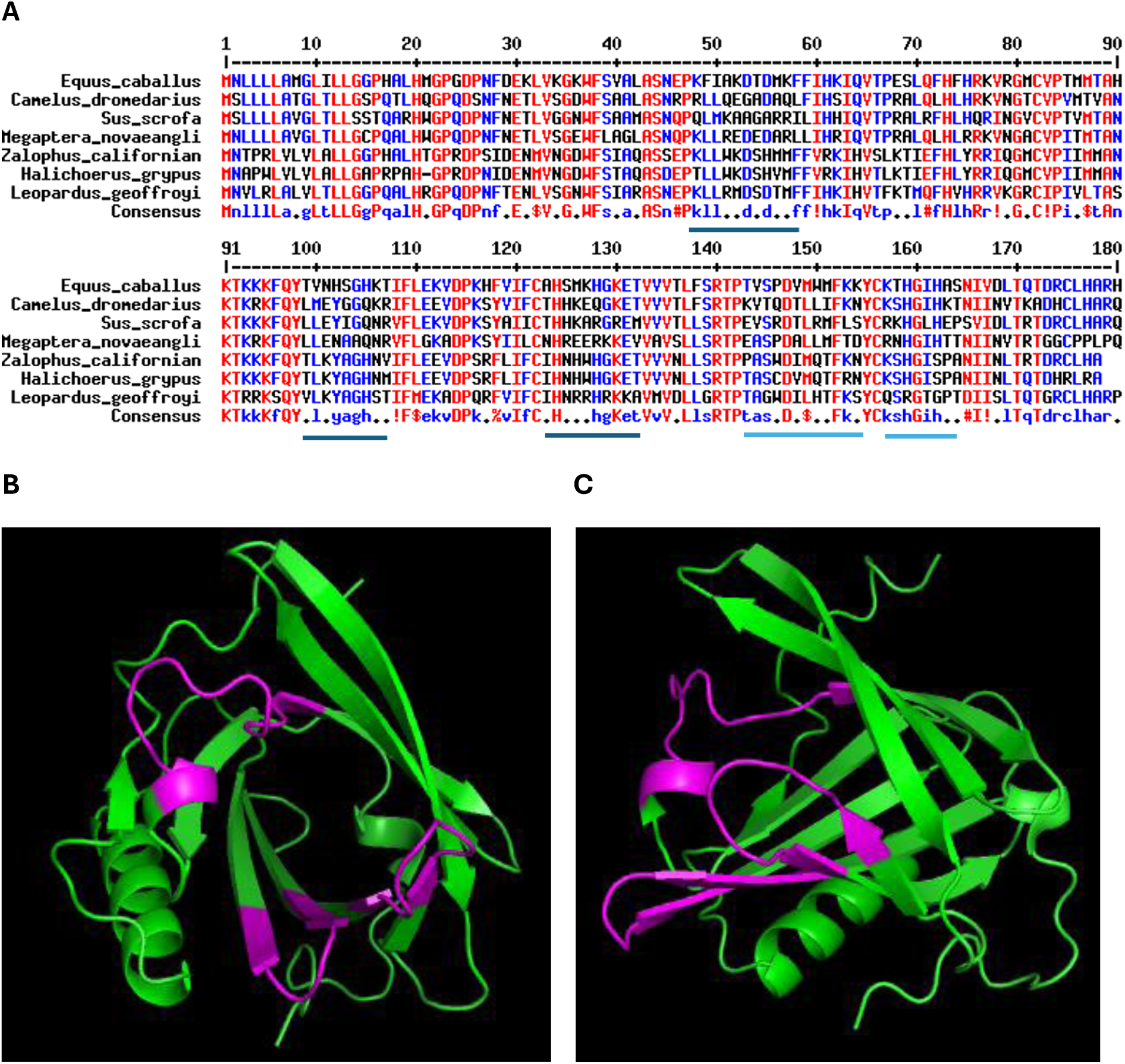
Localised structural specialisation unique to the perissodactyl uterocalins. (A) Alignment of the amino acid sequences of uterocalin-like sequences from two species of camel, a pig, a true seal, a sea lion, and a whale, along with horse uterocalin. In this alignment, the following symbols indicate substitutions of physicochemically similar or commonly inter-substituted amino acids - ! is anyone of I or V, $ is anyone of L or M, % is anyone of F or Y, and # is anyone of N, D, Q, or E. The main regions identified as most noticeably divergent in the alignment for mapping to the structure of the protein are indicated by black bars. (B) and (C) These short stretches of amino acids mapped onto the model structure of equine uterocalin (magenta). The orientation in (B) looks directly into the portal opening to the internal cavity of lipocalins. (C) Shows a side view with the opening to the lipid-binding pocket to the left. The regions of lesser differences are indicated by blue bars in *a* and map to the external helix.

Aside from these confined regions of notable differences, there is sufficient commonality between all the proteins to consider them to have a common ancestor, and that the uterocalin-like proteins from the non-perissodactyls represent an ancestor of the modified perissodactyl uterocalins. Assuming that this is the case, this alignment is useful in illustrating the replacement of leucine in the perissodactyl proteins (table 1). In all but one case leucines are replaced with another essential amino acid, and in seven out of fifteen of these, leucine was replaced with phyenylalanine.

**Table 1.** Preferential loss of leucines in horse uterocalin with biased substitution in favour of phenylalanine. Where in the alignment in figure 5 there is a leucine absent in horse uterocalin when there is one or more in the other Ferungulate sequences. Secretory signal peptides excluded. * Positions where three or four leucines are replaced; ** positions where five are so replaced. Phenylalanines are the most frequent in replacements from leucines – seven out of sixteen.

| *Equus caballus* | *Camelus dromedarius* | *Sus scrofa* | *Megaptera novaeangli* | *Zalophus californianus* | *Halichoerus grypus* | *Leopardus geoffroyi* | Replacement to essential amino acid | Position in alignment |
| --- | --- | --- | --- | --- | --- | --- | --- | --- |
| Ser | Ser | Ser | Leu | Ser | Ser | Ser | No | 39 |
| Phe | Leu | Leu | Leu | Leu | Leu | Leu | Yes | 49 ** |
| Ile | Leu | Met | Leu | Leu | Leu | Leu | Yes | 50 ** |
| Phe | Leu | Ile | Leu | Phe | Phe | Phe | Yes | 58 |
| Phe | Phe | Leu | Leu | Phe | Phe | Phe | Yes | 59 |
| Phe | Leu | Phe | Leu | Phe | Phe | Phe | Yes | 72 |
| Phe | Leu | Leu | Leu | Leu | Leu | Val | Yes | 74 ** |
| Thr | Leu | Leu | Leu | Thr | Thr | Val | Yes | 99 * |
| Val | Met | Leu | Leu | Leu | Leu | Leu | Yes | 100 ** |
| Val | Val | Ala | Ile | Leu | Leu | Val | Yes | 119 |
| Phe | Phe | Ile | Leu | Phe | Phe | Phe | Yes | 121 |
| Phe | Phe | Leu | Leu | Leu | Leu | Leu | Yes | 138 ** |
| Met | Leu | Leu | Leu | Met | Met | Leu | Yes | 149 * |
| Trp | Leu | Arg | Leu | Gln | Gln | His | Yes | 150 |
| Lys | Lys | Leu | Thr | Lys | Arg | Lys | Yes | 154 |
| Ile | Ile | Leu | Ile | Ile | Ile | Thr | Yes | 161 |

### (f) Disposition of the essential amino acids that are in excess in equine uterocalin’s protein structure

Histidine is positively charged, hydrophilic, and concentrates on the surface of the protein where it would be surrounded by polar solvent water (figure 6A). Lysines are also positively charged and, like histidines, are confined to the outside of the protein (figure 6B). Some of the lysines are predicted to project near the portal opening to the internal cavity where their positively charged sidechains could interact to tether the negatively-charged head group of an amphiphilic ligand such as the carboxylate head group of a fatty acid (figure 6B) (43–45).

**Figure 6.**
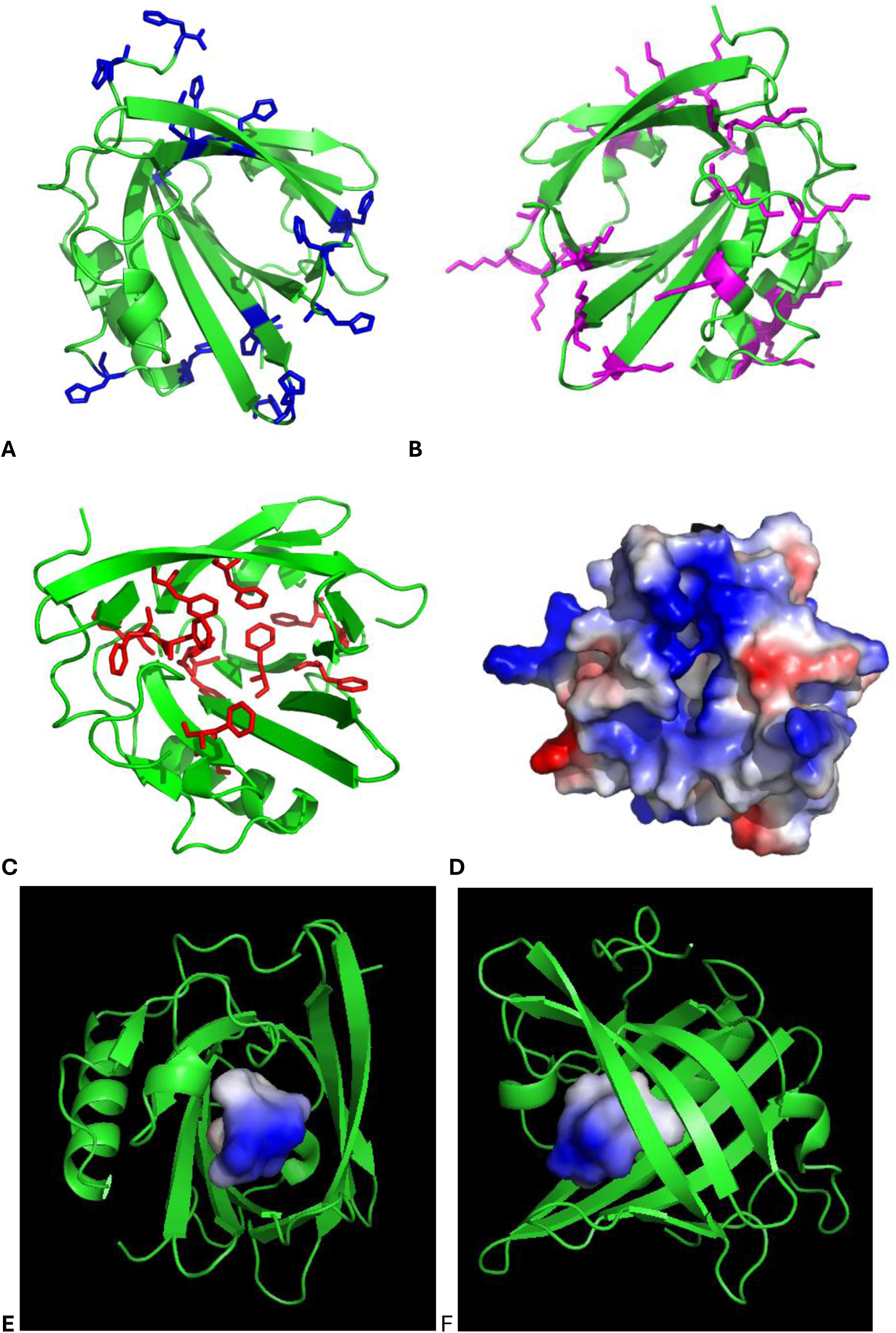
Distribution of essential amino acids in the structure of equine uterocalin. Views of (A) histidine, (B) lysines, and (C) phenylalanines in the structure viewed looking into the opening of the portal to the internal cavity. (C) A space-filling view of the protein oriented to place the entrance to the ligand-binding pocket in the centre of view. Note the ring of positively charged amino acid sidechains round the portal. (E and F) The charge distribution of the walls of the internal cavity of uterocalin viewed straight into the cavity in (E), and to one side in (F). AlphaFold model structure. Blue, positive charge; red, negative; white, neutral/apolar.

The distribution of the strongly hydrophobic essential amino acid phenylalanine is antithetical to that of lysine and histidine in being absent from the outer surface of the protein where they would encounter polar solvent water. Instead, they are either embedded in the structure of the protein or, mostly, line the inner depth of the cavity contributing to its apolar, hydrophobic nature (figure 6C).

Figure 6D provides a space-filling view of the protein coloured according to surface charge and oriented to have the entrance to the ligand-binding pocket in the centre of view and shows a predominance of positively charges (blue) surrounding the portal opening. As noted above, this ring of positive charges would be consistent with tethering the negatively-charged head groups of an amphiphilic ligand as is found in other types of lipid-binding lipocalin (43–45).

The charge distribution of the walls of the internal cavity of uterocalin as viewed straight into the cavity (figure 6E) again indicates a preponderance of positive charges around the mouth of the portal. A side view (figure 6F) shows the depth of the cavity where phenyalanines dominate to create a hydrophobic/apolar pocket where the hydrophobic hydrocarbon tail of a fatty acid or the aliphatic ring of a retinoid, could be held. This arrangement of charged head group of a ligand oriented to the opening of the cavity, and the apolar tail into an apolar core, is known from molecular structures of lipocalins holding ligands such as retinol, retinoic acid, fatty acids, and other amphipathic compounds (23, 26, 44–47).

### (g) A simple diagnostic for “true” perissodactyl uterocalins?

Taking the predicted isoelectric points (pI) of all of the uterocalin-like proteins dealt with here, and the ratios of the number of lysines versus the number of leucines in them, reveals a distinctive pattern in which all the perissodactyl uterocalins and the uterocalin-like proteins from other species have pIs above 9, but only the former have high lysine/leucine ratios (2 minimum, most higher), the remainder having low ratios (1.2 maximum, most lower) (table 2). A sample of other types of lipocalins shows that the perissodactyl uterocalins are indeed distinctive (supplementary material table S2).

**Table 2.** All uterocalin-like proteins have high predicted isoelectric points, but only perissodactyl uterocalins have high lysine/leucine number ratios. See the supplementary material table S2 for a complete list of the proteins, their database accession numbers, and current descriptors. Supplementary material table S1 also has the same details for a sample of lipocalins that show no resemblance to uterocalins, none of which have predicted pI values as high as uterocalin-like proteins, or high lysine/leucine ratios.

|  | **Isoelectric point (pI)**<br>(theoretical) | **Lysine/leucine ratio**<br>(number in protein) |
| --- | --- | --- |
| **Perissodactyl uterocalins** |  |  |
| Horse | 9.72 | 2.5 |
| Donkey | 9.72 | 2.5 |
| Przewalski's horse or takhi | 9.72 | 2.5 |
| Quagga (zebra) | 9.72 | 2.5 |
| Southern black rhinoceros | 9.72 | 2.22 |
| Southern white rhinoceros | 9.83 | 2.38 |
| Indian tapir | 9.62 | 2.0 |
| **Other uterocalin-like proteins** |  |  |
| Bactrian camel | 9.48 | 0.87 |
| Dromedary camel | 9.48 | 0.87 |
| California sea lion | 9.39 | 1.0 |
| Domestic pig | 10.33 | 0.6 |
| Harbour seal | 9.28 | 0.92 |
| Grey seal | 9.14 | 0.91 |
| Geoffroy's cat | 10.64 | 1.0 |
| Northern fur seal | 9.17 | 1.2 |
| Humpback whale | 9.51 | 0.45 |
| Grey whale | 9.62 | 0.45 |
| Pygmy blue whale | 9.37 | 0.37 |
| Rice's whale | 9.18 | 0.45 |
| Narwhal | 9.04 | 0.39 |

## 4. Discussion

The uterocalin proteins from horses and other equids, two species of rhinoceros, and a tapir, distinguish themselves amongst others within the larger lipocalin family in their remarkable enrichment in selected essential amino acids and similar signs of regional specialisation in the structure of the molecule that are associated with a very particular function. The long pre-placentation period and unusual encapsulation and mobility of the conceptus within the uteri of horses, with no direct cell:cell contact between trophoblast and maternal endometrium during that period, has seemingly necessitated the (co)evolution of a support protein to deliver development-critical lipids but also essential amino acids that are is excess in the very structure of the protein. Essential amino acids are those that animals generally cannot synthesise themselves, so require them from their diets or well-provisioned eggs. Or, in the case of mammalian embryos, from their mothers. The clear close similarities between horse uterocalin and those encoded in the genomes of the other two groups of perissodactyls leads to the contention here that the type of unusual, encapsulated nature of the early horse conceptus is a unifying, ancient characteristic of perissodactyls.

While there is no indication in the literature that tapirs have mobile, encapsulated conceptuses, there is evidence for rhinos. Ultrasonography in two species has revealed horse-like spherical embryo vesicles and evidence of forced movement within the uteri (37–41). Formal demonstration of a capsule would require extraction of embryos, which has not been reported for this stage of pregnancy, but it is unlikely that a spherical embryo could survive uterine peristaltic myometrial contractions without a protective capsule. Assuming that rhinos do have horse-like conceptuses, and given that tapirs are more closely related to rhinos than to horses (20–22), then it is reasonable to consider that horse-like conceptuses are typical of all perissodactyls. The nature of their uterocalin proteins provides direct support for this.

Equine uterocalin is secreted by endometrial glands and has been shown to concentrate in the capsule surrounding their pre-placentation conceptuses and then be transferred to, and internalised by, the developing embryo, ultimately appearing in the yolk sac (16–19, 48–50). It has been shown *in vitro* to bind small lipids such as fatty acids, including essential polyunsaturated fatty acids (PUFAs), and retinol (vitamin A) (15). Fatty acids are crucial to membrane construction and energy metabolism, while PUFAs and retinoids are precursors of a wide range of factors involved in cell:cell interactions, activation, differentiation, and signalling events, and as precursors of morphogens in development (28–32, 51–55). Crucially, polyunsaturated fatty acids such as arachidonic acid are precursors of prostaglandins that equine embryos secrete as contributing factors in maternal recognition of pregnancy (32–34, 56, 57). These small lipids can be relatively insoluble in water and are also susceptible to oxidative damage. Hence, in horses, transport within a protein that localises them to an embryonic structure (initially the capsule), appears to be a sound adaptation when there is no direct cell:cell contact between an embryo’s trophoblast and maternal uterine tissues, as distinct from most other eutherian mammals at this stage of pregnancy. It could therefore be said that this, combined with the proteins’ remarkable enrichment in essential amino acids, means that horse uterocalin (and those of all equids, rhinos and tapirs) not only carries essential nutrient compounds to the embryo, but that it is itself food.

The amino acid sequence alignments of the perissodactyl with non-perissodactyl uterocalins (figure 5A) revealed distinctive regions in the former that are confined to three loop regions in the predicted structure of the protein around the opening of the ligand exchange portal of a lipocalin (figure 5B,C). These regions differed in the sequence alignments between the uterocalin-like proteins from non-perissodactyls but not between those of perissodactyls. This indicates that the perissodactyl uterocalins have distinctive protein structural features beyond their unusual amino acid proportions that discriminate them from related proteins found in other members of the Ferungulata – the examples here traverse the clade to camels, pigs, true seals and sea lions, cetaceans and cats. This implies that the perissodactyl proteins are adapted from an ancestral protein present widely across the Ferungulata and that the circum-portal specialisation indicates a specialised function in the perissodactyls. A possibility is that these loops engage with a receptor on the surface of the trophoblast to capture the protein in a correct orientation for transfer of its lipid cargo, or to internalise the protein, cargo and all.

The two essential amino acids that are proportionately the most over-represented in the perissodactyl uterocalins are lysine and histidine. Sidechains of lysines are predicted to angle into the space surrounding the portal opening of uterocalin such that they may act to tether the negatively-charged head group of, for instance, a fatty acid or retinol (figure 6) (43–45, 58). Both are large, positively charged, and predicted to decorate the outside of the protein, thereby not interfering with the apolar internal lipid-carrying cavity. But, why so many lysines and histidines covering uterocalin’s exterior? One function for them could be through charge interaction with an abundant, negatively-charged component of the capsule, sialic acid (59–61). Sialic acid is commonly placed at the termini of glycosylations, such as the extensive glycosylations in mucins (62–64). In the equine capsule it is proposed to act as an anti-adhesion factor allowing the conceptus to be moved freely until fixation (61). But, could it also act in maintaining the structure of the capsule along with uterocalin? Histidine and lysine charge interactions with sialic acid would by themselves focus the protein and its cargo on the capsule as it is synthesised and enlarged by the embryo (61, 65). Such an activity could double to cross-link and stabilise the capsule components into a coherent, physically robust structure. This chimes with the finding that equine embryos cultured *in vitro* with uterocalin produce better capsules than those without (66). Sialic acid is progressively lost when the capsule begins to lose its structural integrity prior to emergence and fixation (59). This happens at a time when the embryo expresses increasing amounts of neuraminidase (7, 67), which could cleave sialic acid from the carbohydrate side chains of the mucus-like proteoglycans of the capsule. If uterocalin’s rich coating in histidines and lysines were to contribute to cross-linking of these to create a stable matrix, then the embryo-induced loss of sialic acids would result in the decay of such interactions and provide a direct mechanism for programmed dissolution of the capsule and emergence. When that happens, stasis and adherence to the endometrium can occur, uterocalin’s function would be retired until later in pregnancy, and its secretion from the endometrium does indeed diminish rapidly at this time (19, 48, 50, 68). The embryo itself would therefore control when it emerges from the capsule and undergo adhesion to the endometrium in preparation for placentation, coordinated with maternal uterine secretions.

If uterocalin were to stabilise the proteoglycan components of the capsule, then would this impede it in delivering its cargo and itself to the embryo? A non-covalent charge interaction would encompass dynamic exchange between bound and free forms such that the protein could ultimately diffuse to the trophoblast surface, which it may then be important to stabilise capsule material newly secreted by the embryo. The binding of fatty acids and retinol to uterocalin would also not be covalent, which may allow its cargo compounds to jump from protein to protein down a concentration gradient to the trophoblast.

Lysine is particularly abundant in the perissodactyl uterocalins and aside from its role as raw material for protein synthesis, it has many roles in physiology and metabolism (69). It is involved in modification of proteins such as in its cross-linking of collagen fibres, and a role in histone modification that could be related to gene activation and silencing crucial to embryogenesis (70, 71). Equally important could be its conversion to trimethyllysine and then carnitine, which is central to the importation of fatty acids into mitochondria and hence energy generation (69). If the energy metabolism of equine embryos is heavily dependent on fatty acids, and equine ova/embryos are notably endowed with lipids (72, 73), then an enhanced supply of lysine for carnitine is explicable. It may not be coincidental that uterocalin is both an enhanced source of lysine and a transporter of fatty acids.

The preponderance of histidines in perissodactyl uterocalins could have a bearing on another potential function of the protein. Histidines are involved in binding to divalent cations such as zinc, copper, and iron, in a wide range of protein types proteins (74–80).

Pertinently, a lipocalin present in the milk of marsupials, trichosurin, binds a zinc atom between two histidines on its surface (81). It could therefore be that an additional activity of uterocalin is to facilitate metal ion acquisition by the conceptus, and such metal ions bound externally may also affect interaction with the capsule matrix.

Also of potential relevance to the capsule is the preponderance of another essential amino acid in the uterocalins, threonine. Threonine is broken down to provide intermediates in energy metabolism and is also itself a fuel (69), but it is the second most abundant amino acid in equine capsular mucus-like proteoglycans (60) and, like the non-essential amino acid serine, is directly involved in O-linked glycosylation of proteins such as mucins (82–84).

Phenylalanine is also over-represented in the perissodactyl uterocalins. It is a highly hydrophobic amino acid with a bulky aromatic side chain. As is typical in soluble proteins, the phenylalanines of uterocalin are embedded within the structure of the protein, removed from external polar solvent, and in uterocalin their sidechains are predicted to project prominently from the walls of the apolar depth of the binding pocket where they may interact with the hydrophobic end of the amphiphilic compounds that equine uterocalin is known to bind (15). Inspection of the amino acid sequence alignment comparing perissodactyl and non-perissodactyl uterocalins (figure 5; Table 1) shows that half of the substitutions from leucines are to phenylalanines in horse uterocalin, which is regarded as an evolutionarily neutral substitution in proteins (85). But in what way can phenylalanine be so important to the embryo that leucines are sacrificed? Phenylalanine is a precursor to tyrosine, which is at slightly lower than average levels in the perissodactyl uterocalins, but this may not explain the distinct preference for phenylalanine over leucine. Phenylalanine is a precursor of the monoamine neurotransmitters dopamine, norepinephrine (noradrenaline), epinephrine (adrenaline), and melanin, all of which could be of particular importance to a developing embryo (69).

Leucine is exceptional amongst uterocalin’s essential amino acids in being in clear deficit in all of the horse, tapir and rhinoceros uterocalins. This is the more remarkable given that leucine is the most abundant amino acid in proteins as a whole. One possibility for the relative lack of leucine is that it is a branched-chain, hydrophobic amino acid with an aliphatic side chain having a high hydropathicity index, that may disrupt the structure of a relatively small soluble protein like uterocalin. This explanation is confounded, however, by the normal levels of leucines in lipocalins in general (e.g. Table S1) and in the non-perissodactylid uterocalin-like proteins dealt with here. It is thus difficult to argue for a structural explanation for the deficit in leucines.

However, leucine is involved at high levels in aerobic mitochondrial metabolism, which can consume the majority of it (69). Human embryos, for instance, degrade leucines more than they do any other amino acid (86). Which therefore deepens the mystery of a deficit in the embryo-provisioning uterocalin. The vast majority of leucine is metabolised in mitochondria contributing to the synthesis of ketones, steroids, fatty acids, acetyl-CoA and other compounds, processes that may not be so crucial to embryo metabolism in a low oxygen environment (69). Early mammalian embryos in general exist in low oxygen conditions (including humans) (2, 3, 87–89), which is likely to be the case for encapsulated equine conceptuses without close contact between the trophoblast and maternal tissue. Leucine may therefore not be required at usual levels by equine embryos, or that there is an as yet unknown other source of leucine (e.g. ref, (90)). If the low content of leucine in uterocalins reflects a reduced need for it in oxidative energy metabolism, then this appears to contradict the above suggestion for high levels of lysine – conversion to carnitine for fatty acid import into mitochondria in oxidative metabolism.

Leucine is a central indicator signal in cellular responses to nutrient availability, and a trigger for growth, protein synthesis, and developmental progression through its interaction with the cellular regulatory-associated protein protein, mTORC1 – if leucine levels are low, mTORC1 signalling induces additional broad spectrum nutrient acquisition by cells (91, 92). Equine embryos, however, may not be nutrient deficient at the encapsulated stage of development, so the leucine deficit in uterocalin may be irrelevant, or that mTORC1 is stimulated continuously to maintain continuous high level nutrient acquisition in an embryo.

Arginine, a conditionally essential amino acid, is at noticeably low levels in the perissodactyl uterocalins and is picked out in VIP analysis as a distinctive feature of perissodactyl uterocalins (figure 4C). This is mysterious because it, like leucine, has a pivotal role in regulating embryo growth and differentiation in mammals, particularly at the blastocyst stage (93). Arginine is involved in many metabolic and physiological processes involved in development, including signalling via nitric oxide synthesis (94, 95). It would be useful to know whether horse embryos can synthesise arginine, or whether it is for them an essential amino acid.

While the unusual amino acid content balance in the perissodactyl uterocalins still requires explanation, it may nevertheless provide useful clues to the metabolism of encapsulated embryos. Assuming that uterocalins are indeed as important in early-stage pregnancy in rhinos and tapirs as they appear to be in equids, might a fuller understanding of them be useful in conservation and assisted reproduction of endangered species, such as cultivation of embryos following *in vitro* fertilisation in preparation for embryo implantation? For instance, discovering why five or six of the nine essential amino acids are at unusually high levels in uterocalin could signal something critical about the metabolism of embryos of pertinence to improving culture media. Equally, it would be important to establish with which lipids or other compounds uterocalin is naturally loaded by the mothers for delivery to their embryo so that an appropriate balance can be ensured for uterocalin added to culture media. Where recombinant uterocalin is made in, for instance, bacteria or fungal protein expression systems, then the accompanying lipids may need to be replaced with more relevant lipids in order to avoid suboptimal or even deleterious *in vitro* culture outcomes.

If it is true that all perissodactyls have long, mobile, encapsulated embryo phases in pregnancy, then this character has persisted since before the Cretaceous-Paleogene extinction event. Is it therefore the case that extant perissodactyls exhibit a primitive character amongst eutherians, or a separate specialisation? Did the evolution of viviparity in all eutherians pass through a stage with horse-like encapsulated embryos that only perissodactyls have retained? Encapsulation of some kind occurs in some eutherians and metatherians (marsupials) (1), but none to the scale and persistence found in horses. And it is not known whether any of these capsules are homologous to, or have similar glycoprotein/proteoglycan components as, those of horses. So, if encapsulation were a specialisation only within perissodactyls then it indeed appears that there has been adaptation of an ancestral uterocalin-like protein (as identified here) to solve the problem presented by a robust capsule. The movement of the conceptus around and between the paired uterine horns is essential to maternal recognition of pregnancy in horses, and likely also rhinos at least, so was that the selective force behind the evolution of the uterocalin/capsule system?

## Ethics statement

This work is entirely based on free, open-access information from genome and protein sequence databases and has no ethical implications.

## Data accessibility

The original data presented in the study are included in the article or in the supplementary material.

## Declaration of AI use

No AI-assisted technologies were used in writing this article.

## Conflict of interest declaration

The author declares no competing interests.

## Author contributions

M.W.K - conceptualization, investigation, project administration, analysis, validation, visualization, writing.

## Funding

None.

## Supporting information

Supplementary to Uterocalin: a protein uniquely specialised to support mobile encapsulated embryos of horses, rhinos and tapirs

## Acknowledgements

Professor Keith Betteridge deserves particular thanks for our many conversations over this subject and his encouragement in pursuing it in understanding the evolution of an unusual type of reproduction in mammals and the particular mystery of perissodactyls. Many thanks also go to Monika Mihm-Carmichael and Neal Dawson for their extremely useful advice on mammal embryology and metabolic specialisations. Particular thanks go to Terri Roth who provided information on rhino embryos that helped make the case made in this paper the more convincing.

## Notes

### Competing Interest Statement

The authors have declared no competing interest.

## References

1. Wooding P, Burton G. Comparative Placentation. Structures, Functions and Evolution. Berlin Heidelberg: Springer Verlag; 2008.

2. Burton GJ, Jauniaux E. Placentation in the Human and Higher Primates. In: Geisert RD, Spencer T, editors. Placentation in Mammals: Tribute to EC Amoroso’s Lifetime Contributions to Viviparity. Cham: Springer International Publishing; 2021. p. 223–54.

3. Burton G, Jauniaux E. The human placenta: new perspectives on its formation and function during early pregnancy. Proceedings of the Royal Society B-Biological Sciences. 2023;290(1997).

4. Gardón JC, Velasco-Martínez MG, Satué K. Early Embryonic Development in the Mare: From Fertilization to Implantation. In: Gardón JC, Satué Ambrojo K, editors. Assisted Reproductive Technologies in Animals Volume 1: Current Trends for Reproductive Management. Cham: Springer Nature Switzerland; 2024. p. 427–71.

5. Ginther O. Equine embryo mobility. A game changer. Theriogenology. 2021;174:131–8.

6. Mcdowell K, Sharp D, Grubaugh W, Thatcher W, Wilcox C. Restricted conceptus mobility results in failure of pregnancy maintenance in mares. Biology of Reproduction. 1988;39(2):340–8.

7. Klein C. Early pregnancy in the mare: old concepts revisited. Domestic Animal Endocrinology. 2016;56:S212–S7.

8. Klein C. Pregnancy Recognition and Implantation of the Conceptus in the Mare. In: Geisert RD, Bazer FW, editors. Regulation of Implantation and Establishment of Pregnancy in Mammals: Tribute to 45 Year Anniversary of Roger V Short’s Maternal Recognition of Pregnancy. Advances in Anatomy Embryology and Cell Biology. 216 2015. p. 165–88.

9. Waelchli R, Betteridge K. Osmolality of equine blastocyst fluid from day 11 to day 25 of pregnancy. Reproduction Fertility and Development. 1996;8(6):981–8.

10. Budik S, Walter I, Tschulenk W, HeIrnreich M, Deichsel K, Pittner F, et al. Significance of aquaporins and sodium potassium ATPase subunits for expansion of the early equine conceptus. Reproduction. 2008;135(4):497–508.

11. Crews L, Waelchli R, Betteridge K. Developmental changes in the distribution of electrolytes within the horse conceptus during the second week of pregnancy. Theriogenology. 2002;58(2-4):745–8.

12. Crews L, Waelchli R, Huang C, Canny M, McCully M, Betteridge K. Electrolyte distribution and yolk sac morphology in frozen hydrated equine conceptuses during the second week of pregnancy. Reproduction Fertility and Development. 2007;19(7):804– 14.

13. Alvarez E, Ortiz-Rodríguez J, Martín-Cano F, Alvarez-Barrientos A, Becerro-Rey L, Gil M, et al. Matched embryo-endometrium RNA-seq reveals coordinated but asymmetric transcriptomic reprogramming at the onset of early equine pregnancy. Frontiers in Cell and Developmental Biology. 2026;14.

14. Amorim M, Bramer S, Rajamanickam G, Klein C, Card C. Endometrial and luteal gene expression of putative gene regulators of the equine maternal recognition of pregnancy. Animal Reproduction Science. 2022;245.

15. Suire S, Stewart F, Beauchamp J, Kennedy M. Uterocalin, a lipocalin provisioning the preattachment equine conceptus: fatty acid and retinol binding properties, and structural characterization. Biochemical Journal. 2001;356:369–76.

16. Crossett B, Allen W, Stewart F. A 19 kDa protein secreted by the endometrium of the mare is a novel member of the lipocalin family. Biochemical Journal. 1996;320:137–43.

17. Crossett B, Suire S, Herrler A, Allen W, Stewart F. Transfer of a uterine lipocalin from the endometrium of the mare to the developing equine conceptus. Biology of Reproduction. 1998;59(3):483–90.

18. Stewart F, Charleston B, Crossett B, Barker P, Allen W. A novel uterine protein that associates with the embryonic capsule in equids. Journal of Reproduction and Fertility. 1995;105(1):65–70.

19. Quinn BA, Hayes MA, Waelchli R, Kennedy MW, Betteridge KJ. Changes in major proteins in the embryonic capsule during immobilization (fixation) of the conceptus in the third week of pregnancy in the mare. Reproduction. 2007;134(1):161–70.

20. Foley N, Mason V, Harris A, Bredemeyer K, Damas J, Lewin H, et al. A genomic timescale for placental mammal evolution. Science. 2023;380(6643):365.

21. Liu G, Pan Q, Du J, Zhu P, Liu W, Li Z, et al. Improved mammalian family phylogeny using gap-rare multiple sequence alignment: A timetree of extant placentals and marsupials. Zoological Research. 2023;44(6):1064–79.

22. Liu S, Westbury M, Dussex N, Mitchell K, Sinding M, Heintzman P, et al. Ancient and modem genomes unravel the evolutionary history of the rhinoceros family. Cell. 2021;184(19):4874–+.

23. Grzyb J, Latowski D, Strzalka K. Lipocalins - a family portrait. Journal of Plant Physiology. 2006;163(9):895–915.

24. Chandrasekaran P, Weiskirchen S, Weiskirchen R. Structure, functions, and implications of selected lipocalins in human disease. International Journal of Molecular Sciences. 2024;25(8).

25. Redl B, Habeler M. The diversity of lipocalin receptors. Biochimie. 2022;192:22– 9.

26. Stopková R, Otcenásková T, Matejková T, Kuntová B, Stopka P. Biological roles of lipocalins in chemical communication, reproduction, and regulation of microbiota. Frontiers in Physiology. 2021;12.

27. Yang H, Wang X, Li S, Liu Y, Akbar R, Fan G. Lipocalin family proteins and their diverse roles in cardiovascular disease. Pharmacology & Therapeutics. 2023;244.

28. Schilling T, Nie Q, Lander A. Dynamics and precision in retinoic acid morphogen gradients. Current Opinion in Genetics & Development. 2012;22(6):562–9.

29. Dollé P, Vermot J, Gallego-Llamas J, Fraulob V, Niederreither K, Chambon P. Retinoic acid controls the bilateral symmetry of somite formation in the mouse embryo. Mechanisms of Development. 2005;122:S7–S.

30. Vilhais-Neto G, Maruhashi M, Smith K, Vasseur-Cognet M, Peterson A, Workman J, et al. Rere controls retinoic acid signalling and somite bilateral symmetry. Nature. 2010;463(7283):953–7.

31. Berry E, Liu Y, Chen L, Guo A. Eicosanoids: Emerging contributors in stem cell-mediated wound healing. Prostaglandins & Other Lipid Mediators. 2017;132:17–24.

32. Kalish B, Kieran M, Puder M, Panigrahy D. The growing role of eicosanoids in tissue regeneration, repair, and wound healing. Prostaglandins & Other Lipid Mediators. 2013;104:130–8.

33. Pelus L, Hoggatt J. Pleiotropic effects of prostaglandin E2 in hematopoiesis; prostaglandin E2 and other eicosanoids regulate hematopoietic stem and progenitor cell function. Prostaglandins & Other Lipid Mediators. 2011;96(1-4):3–9.

34. Budik S, Walter I, Leitner M, Ertl R, Aurich C. Expression of enzymes associated with prostaglandin synthesis in equine conceptuses. Animals. 2021;11(4).

35. Stout T, Allen W. Role of prostaglandins in intrauterine migration of the equine conceptus. Reproduction. 2001;121(5):771–5.

36. Cross D, Ginther O. The effect of estrogen, progesterone and prostaglandin-F2-alpha on uterine contractions in seasonally anovulatory mares. Domestic Animal Endocrinology. 1987;4(4):271–8.

37. Roth T. That was then, this is now - Over two decades of progress in rhinoceros reproductive science and technology. Theriogenology Wild. 2024;4.

38. Stoops M, Campbell M, DeChant C, Hauser J, Kottwitz J, Pairan R, et al. Enhancing captive Indian rhinoceros genetics via artificial insemination of cryopreserved sperm. Animal Reproduction Science. 2016;172:60–75.

39. Roth T, O’Brien J, McRae M, Bellem A, Romo S, Kroll J, et al. Ultrasound and endocrine evaluation of the ovarian cycle and early pregnancy in the Sumatran rhinoceros, Dicerorhinus sumatrensis. Reproduction. 2001;121(1):139–49.

40. Adams G, Plotka E, Asa C, Ginther O. Feasibility of characterizing reproductive events in large nondomestic species by transrectal ultrasonic-imaging. Zoo Biology. 1991;10(3):247–59.

41. Ginther O. How ultrasound technologies have expanded and revolutionized research in reproduction in large animals. Theriogenology. 2014;81(1):112–25.

42. Paterson R, Mackie M, Capobianco A, Heckeberg N, Fraser D, Demarchi B, et al. Phylogenetically informative proteins from an Early Miocene rhinocerotid. Nature. 2025;643(8072).

43. Meenan N, Ball G, Bromek K, Uhrín D, Cooper A, Kennedy M, et al. Solution structure of a repeated unit of the ABA-1 nematode polyprotein allergen of ascaris reveals a novel fold and two discrete lipid-binding sites. Plos Neglected Tropical Diseases. 2011;5(4).

44. Papiz M, Sawyer L, Eliopoulos E, North A, Findlay J, Sivaprasadarao R, et al. The structure of beta-lactoglobulin and its similarity to plasma retinol-binding protein. Nature. 1986;324(6095):383–5.

45. Wu S, Pèrez M, Puyol P, Sawyer L. β-lactoglobulin binds palmitate within its central cavity. Journal of Biological Chemistry. 1999;274(1):170–4.

46. Breustedt DA, Schonfeld DL, Skerra A. Comparative ligand-binding analysis of ten human lipocalins. Biochimica Et Biophysica Acta-Proteins and Proteomics. 2006;1764(2):161–73.

47. Newcomer M, Jones T, Aqvist J, Sundelin J, Eriksson U, Rask L, et al. The 3-dimensional structure of retinol-binding protein. Embo Journal. 1984;3(7):1451–4.

48. Hayes M, Quinn B, Keirstead N, Katavolos P, Waelchli R, Betteridge K. Proteins associated with the early intrauterine equine conceptus. Reproduction in Domestic Animals. 2008;43:232–7.

49. Klein C, Bruce P, Hammermueller J, Hayes T, Lillie B, Betteridge K. Transcriptional profiling of equine endometrium before, during and after capsule disintegration during normal pregnancy and after oxytocin-induced luteostasis in non-pregnant mares. Plos One. 2021;16(10).

50. Ellenberger C, Wilsher S, Allen W, Hoffmann C, Kölling M, Bazer F, et al. Immunolocalisation of the uterine secretory proteins uterocalin, uteroferrin and uteroglobin in the mare’s uterus and placenta throughout pregnancy. Theriogenology. 2008;70(5):746–57.

51. Yang C, Mai D, Pan Z, Xue Y, Wang Y, Yan C. Recent advances in the methods and applications used to analyze eicosanoids. Chinese Journal of Chromatography. 2016;34(5):449–55.

52. Kane M. Retinoic acid homeostasis and disease. In: Duester G, Ghyselinck NB, editors. Retinoids in Development and Disease. Current Topics in Developmental Biology. 1612025. p. 201–33.

53. Oliveira L, Teixeira F, Sato M. Impact of retinoic acid on immune cells and inflammatory diseases. Mediators of Inflammation. 2018;2018:3067126.

54. Polcz M, Barbul A. The role of vitamin a in wound healing. Nutrition in Clinical Practice. 2019;34(5):695–700.

55. Vermot J, Llamas J, Fraulob V, Niederreither K, Chambon P, Dollé P. Retinoic acid controls the bilateral symmetry of somite formation in the mouse embryo. Science. 2005;308(5721):563–6.

56. Klein C. Maternal recognition of pregnancy in the context of equine embryo transfer. Journal of Equine Veterinary Science. 2016;41:22–8.

57. Calder P. Eicosanoids. Lipid Mediators. 2020;64(3):423–41.

58. Curry S, Mandelkow H, Brick P, Franks N. Crystal structure of human serum albumin complexed with fatty acid reveals an asymmetric distribution of binding sites. Nature Structural Biology. 1998;5(9):827–35.

59. Arar S, Chan K, Quinn B, Waelchli R, Hayes M, Betteridge K, et al. Desialylation of core type 1 O-glycan in the equine embryonic capsule coincides with immobilization of the conceptus in the uterus. Carbohydrate Research. 2007;342(8):1110–5.

60. Oriol J, Betteridge K, Clarke A, Sharom F. Mucin-like glycoproteins in the equine embryonic capsule. Molecular Reproduction and Development. 1993;34(3):255–65.

61. Oriol J, Sharom F, Betteridge K. Developmentally-regulated changes in the glycoproteins of the equine embryonic capsule. Journal of Reproduction and Fertility. 1993;99(2):653–64.

62. Chatterjee M, van Putten J, Strijbis K. Defensive properties of mucin glycoproteins during respiratory infections-relevance for SARS-CoV-2. Mbio. 2020;11(6).

63. Lillehoj E, Luzina I, Atamas S. Mammalian Neuraminidases in Immune-Mediated Diseases: Mucins and Beyond. Frontiers in Immunology. 2022;13.

64. Ma X, Li M, Wang X, Qi G, Wei L, Zhang D. Sialylation in the gut: From mucosal protection to disease pathogenesis. Carbohydrate Polymers. 2024;343.

65. Albihn A, Waelchli R, Samper J, Oriol J, Croy B, Betteridge K. Production of capsular material by equine trophoblast transplanted into immunodeficient mice. Reproduction. 2003;125(6):855–63.

66. Smits K, Govaere J, Peelman L, Goossens K, de Graaf D, Vercauteren D, et al. Influence of the uterine environment on the development of in vitro-produced equine embryos. Reproduction. 2012;143(2):173–81.

67. Klein C, Troedsson M. Equine pre-implantation conceptuses express neuraminidase 2 - A potential mechanism for desialylation of the equine capsule. Reproduction in Domestic Animals. 2012;47(3):449–54.

68. Hoffmann C, Bazer FW, Klug J, Aupperle H, Ellenberger C, Schoon HA. Immunohistochemical and histochemical identification of proteins and carbohydrates in the equine endometrium Expression patterns for mares suffering from endometrosis. Theriogenology. 2009;71(2):264–74.

69. Kohlmeier M. Amino acids and nitrogen compounds. In: Kohlmeier M, editor. Nutrient Metabolism: Structures, Functions, and Genes. Second ed. Amsterdam, Boston: Academic Press; 2015. p. 266–477.

70. Dambacher S, Hahn M, Schotta G. Epigenetic regulation of development by histone lysine methylation. Heredity. 2010;105(1):24–37.

71. Martin C, Zhang Y. The diverse functions of histone lysine methylation. Nature Reviews Molecular Cell Biology. 2005;6(11):838–49.

72. Lawson E, Pickford R, Aitken R, Gibb Z, Grupen C, Swegen A. Mapping the lipidomic secretome of the early equine embryo. Frontiers in Veterinary Science. 2024;11.

73. Lawson EF, Grupen CG, Baker MA, Aitken RJ, Swegen A, Pollard CL, et al. Conception and early pregnancy in the mare: lipidomics the unexplored frontier. Reproduction and Fertility. 2022;3(1):R1–R18.

74. Eriksson A, Jones T, Liljas A. Refined structure of human carbonic anhydrase-II at 2.0-Å resolution. Proteins-Structure Function and Bioinformatics. 1988;4(4):274–82.

75. Ferraroni M, Tilli S, Briganti F, Chegwidden W, Supuran C, Wiebauer K, et al. Crystal structure of a zinc-activated variant of human carbonic anhydrase I, CA I Michigan 1: Evidence for a second zinc binding site involving arginine coordination. Biochemistry. 2002;41(20):6237–44.

76. Jones A, Hulett M, Parish C. Histidine-rich glycoprotein: A novel adaptor protein in plasma that modulates the immune, vascular and coagulation systems. Immunology and Cell Biology. 2005;83(2):106–18.

77. Pandey A, Babbarwal V, Okoyeh J, Joshi R, Puri S, Singh R, et al. Hemozoin formation in malaria: a two-step process involving histidine-rich proteins and lipids. Biochemical and Biophysical Research Communications. 2003;308(4):736–43.

78. Yang Y, Tang T, Feng B, Li S, Hou N, Ma X, et al. Disruption of Plasmodium falciparum histidine-rich protein 2 may affect haem metabolism in the blood stage. Parasites & Vectors. 2020;13(1).

79. Lukin J, Ho C. The structure-function relationship of hemoglobin in solution at atomic resolution. Chemical Reviews. 2004;104(3):1219–30.

80. Oke M, Ching R, Carter L, Johnson K, Liu H, McMahon S, et al. Unusual chromophore and cross-links in ranasmurfin: A blue protein from the foam nests of a tropical frog. Angewandte Chemie-International Edition. 2008;47(41):7853–6.

81. Watson R, Demmer J, Baker E, Arcus V. Three-dimensional structure and ligand binding properties of trichosurin, a metatherian lipocalin from the milk whey of the common brushtail possum Trichosurus vulpecula. Biochemical Journal. 2007;408:29– 38.

82. Bansil R, Turner B. The biology of mucus: Composition, synthesis and organization. Advanced Drug Delivery Reviews. 2018;124:3–15.

83. Hanisch F. O-glycosylation of the mucin type. Biological Chemistry. 2001;382(2):143–9.

84. Peter-Katalinic J. Methods in enzymology: O-glycosylation of proteins. In: Burlingame AL, editor. Mass Spectrometry: Modified Proteins and Glycoconjugates. Methods in Enzymology. 405 2005. p. 139–71.

85. Dayhoff M, Schwartz R, Orcutt B. Atlas of Protein Sequence and Structure. Washington, USA: National Biomedical Research Foundation; 1979.

86. Houghton F, Hawkhead J, Humpherson P, Hogg J, Balen A, Rutherford A, et al. Non-invasive amino acid turnover predicts human embryo developmental capacity. Human Reproduction. 2002;17(4):999–1005.

87. García-Martínez S, Hurtado M, Gutiérrez H, Margallo F, Romar R, Latorre R, et al. Mimicking physiological O2 tension in the female reproductive tract improves assisted reproduction outcomes in pig. Molecular Human Reproduction. 2018;24(5):260–70.

88. Jauniaux E, Poston L, Burton G. Placental-related diseases of pregnancy: involvement of oxidative stress and implications in human evolution. Human Reproduction Update. 2006;12(6):747–55.

89. Lewis N, Hinrichs K, Leese H, Argo C, Brison D, Sturmey R. Energy metabolism of the equine cumulus oocyte complex during in vitro maturation. Scientific Reports. 2020;10(1).

90. Liu BM, Duan LK, Liu XJ, Bazer FW, Wang XQ. Uterine histotroph and conceptus development. IV. Metabolomic analyses of uterine luminal fluid reveals regulatory landscapes during the peri-implantation period of pregnancy in pigs. Biology of Reproduction. 2025;113(6):1523–38.

91. Koundouros N, Nagiec M, Bullen N, Noch E, Burgos-Barragan G, Li Z, et al. Direct sensing of dietary ω-6 linoleic acid through FABP5-mTORC1 signaling. Science. 2025;387(6739).

92. Yang S, Bian T, Zoncu R. Molecular and structural mechanisms of nutrient sensing in the mTORC1 pathway. Trends in Biochemical Sciences. 2026;51(5):457–73.

93. Leese H, McKeegan P, Sturmey R. Amino acids and the early mammalian embryo: origin, fate, function and life-long legacy. International Journal of Environmental Research and Public Health. 2021;18(18).

94. Mehmood T, Ali M, Javed M, Mateen R, Nouman M, Knani S, et al. Multifaceted roles of arginine in cellular function, signaling, and disease. Frontiers in Chemistry. 2026;14.

95. Wu G, Bazer FW, Satterfield MC, Li X, Wang X, Johnson GA, et al. Impacts of arginine nutrition on embryonic and fetal development in mammals. Amino Acids. 2013;45(2):241–56.

