## Supplementary to Uterocalin: a protein uniquely specialised to support mobile encapsulated embryos of horses, rhinos and tapirs for "Uterocalin – a protein uniquely specialised to support mobile encapsulated embryos of horses, rhinos and tapirs"

#### **Contents**

**Table S1. Amino acid sequences of all the proteins dealt with in the paper along with database accession codes, how some were edited, and their amino acid compositions and theoretical isoelectric points (pI).**

**Figure S1. Heatmap, Principal Component Analysis, and VIP analysis of amino acid content of perissodactyl uterocalins, uterocalin-like proteins from non-perissodactyls, and a selection of unrelated lipocalins.**

**Table S2. A simple diagnostic for “true” horse, rhinoceros and tapir uterocalins? These have high predicted isoelectric points (pI) combined with high lysine/leucine content ratios.**

**References.**

**Table S1.** Amino acid sequences of all the proteins dealt with in the paper along with database accession codes, how some were edited, and their amino acid compositions and theoretical isoelectric points (pI).

All but one of the sequences were obtained from NCBI (https://www.ncbi.nlm.nih.gov/), the remainder being from the *Tapirus indicus* genome resource, as indicated. Beside each entry is the amino acid composition and theoretical pI of each protein calculated by ProtParam (https://web.expasy.org/protparam/).

The “uterocalin-like” protein sequence were derived from simple protein-protein BLAST searches of the NCBI database using the horse sequence as query, and those chosen for examination were those that appeared in the report list immediately following the perissodactyl sequences. Some of the uterocalin-like proteins predicted from genomic data are longer (sometimes in both N- and COOH-terminal directions) than expected of typical lipocalins, although their cores align well with perissodactyl uterocalins. These extensions (indicated in blue) may be due to errors in gene transcription/translation predictions. None of the extensions align with any other protein sequences other than themselves by BLAST, so are likely artifacts. For this reason the extended sequences were excluded from all the calculations used, but also to allow comparisons between similar lengths of core sequence. This was also done to the sequence for horse retinol binding protein since it also has an extended sequence. Sequences that are excluded for any reason are indicated in blue.

The N-terminal secretory signal peptides were predicted by SignalP 6.0 (https://services.healthtech.dtu.dk/services/SignalP-6.0/) set for Eukaryota and are indicated in red. For most of the lipocalins here this left “mature” proteins of 162 amino acids in length, this number varying slightly because of where signal peptides ended, or abbreviation of the mature protein lengths.

Note that all the proteins under the heading “Uterocalin-like” that align well with the equine/perissodactyl forms exhibit high predicted isoelectric point (pI) values, even if they do not possess the high ratios of essential amino acids or the divergent regions noted in the paper that appear to be structural specialisations in the perissodactyl mammals (equids, rhinoceros, tapir).

| Protein<br>(sequences in single letter codes for amino acids) | ProtParam output |  |  |  |  |  |  |  |  |  |  |  |  |  |  |  |  |  |  |  |  |  |  |  |  |  |  |  |  |  |  |  |  |  |  |  |  |  |  |  |  |  |  |  |  |  |  |  |  |  |  |  |  |  |  |  |  |  |  |  |  |  |  |  |  |  |  |
| --- | --- | --- | --- | --- | --- | --- | --- | --- | --- | --- | --- | --- | --- | --- | --- | --- | --- | --- | --- | --- | --- | --- | --- | --- | --- | --- | --- | --- | --- | --- | --- | --- | --- | --- | --- | --- | --- | --- | --- | --- | --- | --- | --- | --- | --- | --- | --- | --- | --- | --- | --- | --- | --- | --- | --- | --- | --- | --- | --- | --- | --- | --- | --- | --- | --- | --- | --- |
| Perissodactyl uterocalins<br>(with current database names/designations) |  |  |  |  |  |  |  |  |  |  |  |  |  |  |  |  |  |  |  |  |  |  |  |  |  |  |  |  |  |  |  |  |  |  |  |  |  |  |  |  |  |  |  |  |  |  |  |  |  |  |  |  |  |  |  |  |  |  |  |  |  |  |  |  |  |  |  |
| uterocalin precursor [Equus caballus]<br>NCBI Reference Sequence: NP_001075978.2<br>mnllllamgl illggpha lh mgpgdpnfde klvkgkwfsv<br>alasnepkfi akdtdmkffi<br>hkiqvtpesl qfhfhrkvrg mcvptmmtah ktkkkfqytv<br>nhsghtifl ekvdpkhfvi<br>fcahsmkhgk etvvvtlfsr tptvspdvmw mfkkyckthg<br>ihasnivdlt qtdrclharh | Number of amino acids: 162<br>Theoretical pI: 9.72<br>Molecular weight: 18763.00<br>Amino acid composition:<br>CSV format<br>Ala (A)<br>7<br>4.3%<br>Arg (R)<br>5<br>3.1%<br>Asn (N)<br>4<br>2.5%<br>Asp (D)<br>8<br>4.9%<br>Cys (C)<br>4<br>2.5%<br>Gln (Q)<br>4<br>2.5%<br>Glu (E)<br>5<br>3.1%<br>Gly (G)<br>7<br>4.3%<br>His (H)<br>14<br>8.6%<br>Ile (I)<br>7<br>4.3%<br>Leu (L)<br>8<br>4.9%<br>Lys (K)<br>20<br>12.3%<br>Met (M)<br>8<br>4.9%<br>Phe (F)<br>13<br>8.0%<br>Pro (P)<br>8<br>4.9%<br>Ser (S)<br>8<br>4.9%<br>Thr (T)<br>14<br>8.6%<br>Trp (W)<br>2<br>1.2%<br>Tyr (Y)<br>2<br>1.2%<br>Val (V)<br>14<br>8.6%<br>Pyl (O)<br>0<br>0.0%<br>Sec (U)<br>0<br>0.0% | Ala (A) | 7 | 4.3% | Arg (R) | 5 | 3.1% | Asn (N) | 4 | 2.5% | Asp (D) | 8 | 4.9% | Cys (C) | 4 | 2.5% | Gln (Q) | 4 | 2.5% | Glu (E) | 5 | 3.1% | Gly (G) | 7 | 4.3% | His (H) | 14 | 8.6% | Ile (I) | 7 | 4.3% | Leu (L) | 8 | 4.9% | Lys (K) | 20 | 12.3% | Met (M) | 8 | 4.9% | Phe (F) | 13 | 8.0% | Pro (P) | 8 | 4.9% | Ser (S) | 8 | 4.9% | Thr (T) | 14 | 8.6% | Trp (W) | 2 | 1.2% | Tyr (Y) | 2 | 1.2% | Val (V) | 14 | 8.6% | Pyl (O) | 0 | 0.0% | Sec (U) | 0 | 0.0% |
| Ala (A) | 7 | 4.3% |  |  |  |  |  |  |  |  |  |  |  |  |  |  |  |  |  |  |  |  |  |  |  |  |  |  |  |  |  |  |  |  |  |  |  |  |  |  |  |  |  |  |  |  |  |  |  |  |  |  |  |  |  |  |  |  |  |  |  |  |  |  |  |  |  |
| Arg (R) | 5 | 3.1% |  |  |  |  |  |  |  |  |  |  |  |  |  |  |  |  |  |  |  |  |  |  |  |  |  |  |  |  |  |  |  |  |  |  |  |  |  |  |  |  |  |  |  |  |  |  |  |  |  |  |  |  |  |  |  |  |  |  |  |  |  |  |  |  |  |
| Asn (N) | 4 | 2.5% |  |  |  |  |  |  |  |  |  |  |  |  |  |  |  |  |  |  |  |  |  |  |  |  |  |  |  |  |  |  |  |  |  |  |  |  |  |  |  |  |  |  |  |  |  |  |  |  |  |  |  |  |  |  |  |  |  |  |  |  |  |  |  |  |  |
| Asp (D) | 8 | 4.9% |  |  |  |  |  |  |  |  |  |  |  |  |  |  |  |  |  |  |  |  |  |  |  |  |  |  |  |  |  |  |  |  |  |  |  |  |  |  |  |  |  |  |  |  |  |  |  |  |  |  |  |  |  |  |  |  |  |  |  |  |  |  |  |  |  |
| Cys (C) | 4 | 2.5% |  |  |  |  |  |  |  |  |  |  |  |  |  |  |  |  |  |  |  |  |  |  |  |  |  |  |  |  |  |  |  |  |  |  |  |  |  |  |  |  |  |  |  |  |  |  |  |  |  |  |  |  |  |  |  |  |  |  |  |  |  |  |  |  |  |
| Gln (Q) | 4 | 2.5% |  |  |  |  |  |  |  |  |  |  |  |  |  |  |  |  |  |  |  |  |  |  |  |  |  |  |  |  |  |  |  |  |  |  |  |  |  |  |  |  |  |  |  |  |  |  |  |  |  |  |  |  |  |  |  |  |  |  |  |  |  |  |  |  |  |
| Glu (E) | 5 | 3.1% |  |  |  |  |  |  |  |  |  |  |  |  |  |  |  |  |  |  |  |  |  |  |  |  |  |  |  |  |  |  |  |  |  |  |  |  |  |  |  |  |  |  |  |  |  |  |  |  |  |  |  |  |  |  |  |  |  |  |  |  |  |  |  |  |  |
| Gly (G) | 7 | 4.3% |  |  |  |  |  |  |  |  |  |  |  |  |  |  |  |  |  |  |  |  |  |  |  |  |  |  |  |  |  |  |  |  |  |  |  |  |  |  |  |  |  |  |  |  |  |  |  |  |  |  |  |  |  |  |  |  |  |  |  |  |  |  |  |  |  |
| His (H) | 14 | 8.6% |  |  |  |  |  |  |  |  |  |  |  |  |  |  |  |  |  |  |  |  |  |  |  |  |  |  |  |  |  |  |  |  |  |  |  |  |  |  |  |  |  |  |  |  |  |  |  |  |  |  |  |  |  |  |  |  |  |  |  |  |  |  |  |  |  |
| Ile (I) | 7 | 4.3% |  |  |  |  |  |  |  |  |  |  |  |  |  |  |  |  |  |  |  |  |  |  |  |  |  |  |  |  |  |  |  |  |  |  |  |  |  |  |  |  |  |  |  |  |  |  |  |  |  |  |  |  |  |  |  |  |  |  |  |  |  |  |  |  |  |
| Leu (L) | 8 | 4.9% |  |  |  |  |  |  |  |  |  |  |  |  |  |  |  |  |  |  |  |  |  |  |  |  |  |  |  |  |  |  |  |  |  |  |  |  |  |  |  |  |  |  |  |  |  |  |  |  |  |  |  |  |  |  |  |  |  |  |  |  |  |  |  |  |  |
| Lys (K) | 20 | 12.3% |  |  |  |  |  |  |  |  |  |  |  |  |  |  |  |  |  |  |  |  |  |  |  |  |  |  |  |  |  |  |  |  |  |  |  |  |  |  |  |  |  |  |  |  |  |  |  |  |  |  |  |  |  |  |  |  |  |  |  |  |  |  |  |  |  |
| Met (M) | 8 | 4.9% |  |  |  |  |  |  |  |  |  |  |  |  |  |  |  |  |  |  |  |  |  |  |  |  |  |  |  |  |  |  |  |  |  |  |  |  |  |  |  |  |  |  |  |  |  |  |  |  |  |  |  |  |  |  |  |  |  |  |  |  |  |  |  |  |  |
| Phe (F) | 13 | 8.0% |  |  |  |  |  |  |  |  |  |  |  |  |  |  |  |  |  |  |  |  |  |  |  |  |  |  |  |  |  |  |  |  |  |  |  |  |  |  |  |  |  |  |  |  |  |  |  |  |  |  |  |  |  |  |  |  |  |  |  |  |  |  |  |  |  |
| Pro (P) | 8 | 4.9% |  |  |  |  |  |  |  |  |  |  |  |  |  |  |  |  |  |  |  |  |  |  |  |  |  |  |  |  |  |  |  |  |  |  |  |  |  |  |  |  |  |  |  |  |  |  |  |  |  |  |  |  |  |  |  |  |  |  |  |  |  |  |  |  |  |
| Ser (S) | 8 | 4.9% |  |  |  |  |  |  |  |  |  |  |  |  |  |  |  |  |  |  |  |  |  |  |  |  |  |  |  |  |  |  |  |  |  |  |  |  |  |  |  |  |  |  |  |  |  |  |  |  |  |  |  |  |  |  |  |  |  |  |  |  |  |  |  |  |  |
| Thr (T) | 14 | 8.6% |  |  |  |  |  |  |  |  |  |  |  |  |  |  |  |  |  |  |  |  |  |  |  |  |  |  |  |  |  |  |  |  |  |  |  |  |  |  |  |  |  |  |  |  |  |  |  |  |  |  |  |  |  |  |  |  |  |  |  |  |  |  |  |  |  |
| Trp (W) | 2 | 1.2% |  |  |  |  |  |  |  |  |  |  |  |  |  |  |  |  |  |  |  |  |  |  |  |  |  |  |  |  |  |  |  |  |  |  |  |  |  |  |  |  |  |  |  |  |  |  |  |  |  |  |  |  |  |  |  |  |  |  |  |  |  |  |  |  |  |
| Tyr (Y) | 2 | 1.2% |  |  |  |  |  |  |  |  |  |  |  |  |  |  |  |  |  |  |  |  |  |  |  |  |  |  |  |  |  |  |  |  |  |  |  |  |  |  |  |  |  |  |  |  |  |  |  |  |  |  |  |  |  |  |  |  |  |  |  |  |  |  |  |  |  |
| Val (V) | 14 | 8.6% |  |  |  |  |  |  |  |  |  |  |  |  |  |  |  |  |  |  |  |  |  |  |  |  |  |  |  |  |  |  |  |  |  |  |  |  |  |  |  |  |  |  |  |  |  |  |  |  |  |  |  |  |  |  |  |  |  |  |  |  |  |  |  |  |  |
| Pyl (O) | 0 | 0.0% |  |  |  |  |  |  |  |  |  |  |  |  |  |  |  |  |  |  |  |  |  |  |  |  |  |  |  |  |  |  |  |  |  |  |  |  |  |  |  |  |  |  |  |  |  |  |  |  |  |  |  |  |  |  |  |  |  |  |  |  |  |  |  |  |  |
| Sec (U) | 0 | 0.0% |  |  |  |  |  |  |  |  |  |  |  |  |  |  |  |  |  |  |  |  |  |  |  |  |  |  |  |  |  |  |  |  |  |  |  |  |  |  |  |  |  |  |  |  |  |  |  |  |  |  |  |  |  |  |  |  |  |  |  |  |  |  |  |  |  |
| uterocalin isoform X2 [Equus asinus]<br>NCBI Reference Sequence: XP_070376020.1<br>Wrong start methionine. Alignment and Signal P prediction gives start as below<br>mvivnrrgp vtachvarnp llaedkratv kgseplsevp<br>lvslshsclt<br>mnllllamgl illggpha lh mgpgdpnfde klvkgkwfsv<br>alasnepkfi akdtdmkffi hkiqvtpesl<br>qfhfhrkvrg mcvptmmtah ktkkkfqytv nhsghtifl<br>ekvdpkhfvi fcahsmkhgk<br>etvvvtlfsr tptvspdvmw mfkkyckthg ihasnivdlt<br>qtdrclharh | Number of amino acids: 162<br>Theoretical pI: 9.72<br>Molecular weight: 18763.00<br>Amino acid composition:<br>CSV format<br>Ala (A)<br>7<br>4.3%<br>Arg (R)<br>5<br>3.1%<br>Asn (N)<br>4<br>2.5%<br>Asp (D)<br>8<br>4.9%<br>Cys (C)<br>4<br>2.5%<br>Gln (Q)<br>4<br>2.5%<br>Glu (E)<br>5<br>3.1%<br>Gly (G)<br>7<br>4.3%<br>His (H)<br>14<br>8.6%<br>Ile (I)<br>7<br>4.3%<br>Leu (L)<br>8<br>4.9%<br>Lys (K)<br>20<br>12.3%<br>Met (M)<br>8<br>4.9%<br>Phe (F)<br>13<br>8.0%<br>Pro (P)<br>8<br>4.9%<br>Ser (S)<br>8<br>4.9%<br>Thr (T)<br>14<br>8.6%<br>Trp (W)<br>2<br>1.2%<br>Tyr (Y)<br>2<br>1.2%<br>Val (V)<br>14<br>8.6%<br>Pyl (O)<br>0<br>0.0%<br>Sec (U)<br>0<br>0.0% | Ala (A) | 7 | 4.3% | Arg (R) | 5 | 3.1% | Asn (N) | 4 | 2.5% | Asp (D) | 8 | 4.9% | Cys (C) | 4 | 2.5% | Gln (Q) | 4 | 2.5% | Glu (E) | 5 | 3.1% | Gly (G) | 7 | 4.3% | His (H) | 14 | 8.6% | Ile (I) | 7 | 4.3% | Leu (L) | 8 | 4.9% | Lys (K) | 20 | 12.3% | Met (M) | 8 | 4.9% | Phe (F) | 13 | 8.0% | Pro (P) | 8 | 4.9% | Ser (S) | 8 | 4.9% | Thr (T) | 14 | 8.6% | Trp (W) | 2 | 1.2% | Tyr (Y) | 2 | 1.2% | Val (V) | 14 | 8.6% | Pyl (O) | 0 | 0.0% | Sec (U) | 0 | 0.0% |
| Ala (A) | 7 | 4.3% |  |  |  |  |  |  |  |  |  |  |  |  |  |  |  |  |  |  |  |  |  |  |  |  |  |  |  |  |  |  |  |  |  |  |  |  |  |  |  |  |  |  |  |  |  |  |  |  |  |  |  |  |  |  |  |  |  |  |  |  |  |  |  |  |  |
| Arg (R) | 5 | 3.1% |  |  |  |  |  |  |  |  |  |  |  |  |  |  |  |  |  |  |  |  |  |  |  |  |  |  |  |  |  |  |  |  |  |  |  |  |  |  |  |  |  |  |  |  |  |  |  |  |  |  |  |  |  |  |  |  |  |  |  |  |  |  |  |  |  |
| Asn (N) | 4 | 2.5% |  |  |  |  |  |  |  |  |  |  |  |  |  |  |  |  |  |  |  |  |  |  |  |  |  |  |  |  |  |  |  |  |  |  |  |  |  |  |  |  |  |  |  |  |  |  |  |  |  |  |  |  |  |  |  |  |  |  |  |  |  |  |  |  |  |
| Asp (D) | 8 | 4.9% |  |  |  |  |  |  |  |  |  |  |  |  |  |  |  |  |  |  |  |  |  |  |  |  |  |  |  |  |  |  |  |  |  |  |  |  |  |  |  |  |  |  |  |  |  |  |  |  |  |  |  |  |  |  |  |  |  |  |  |  |  |  |  |  |  |
| Cys (C) | 4 | 2.5% |  |  |  |  |  |  |  |  |  |  |  |  |  |  |  |  |  |  |  |  |  |  |  |  |  |  |  |  |  |  |  |  |  |  |  |  |  |  |  |  |  |  |  |  |  |  |  |  |  |  |  |  |  |  |  |  |  |  |  |  |  |  |  |  |  |
| Gln (Q) | 4 | 2.5% |  |  |  |  |  |  |  |  |  |  |  |  |  |  |  |  |  |  |  |  |  |  |  |  |  |  |  |  |  |  |  |  |  |  |  |  |  |  |  |  |  |  |  |  |  |  |  |  |  |  |  |  |  |  |  |  |  |  |  |  |  |  |  |  |  |
| Glu (E) | 5 | 3.1% |  |  |  |  |  |  |  |  |  |  |  |  |  |  |  |  |  |  |  |  |  |  |  |  |  |  |  |  |  |  |  |  |  |  |  |  |  |  |  |  |  |  |  |  |  |  |  |  |  |  |  |  |  |  |  |  |  |  |  |  |  |  |  |  |  |
| Gly (G) | 7 | 4.3% |  |  |  |  |  |  |  |  |  |  |  |  |  |  |  |  |  |  |  |  |  |  |  |  |  |  |  |  |  |  |  |  |  |  |  |  |  |  |  |  |  |  |  |  |  |  |  |  |  |  |  |  |  |  |  |  |  |  |  |  |  |  |  |  |  |
| His (H) | 14 | 8.6% |  |  |  |  |  |  |  |  |  |  |  |  |  |  |  |  |  |  |  |  |  |  |  |  |  |  |  |  |  |  |  |  |  |  |  |  |  |  |  |  |  |  |  |  |  |  |  |  |  |  |  |  |  |  |  |  |  |  |  |  |  |  |  |  |  |
| Ile (I) | 7 | 4.3% |  |  |  |  |  |  |  |  |  |  |  |  |  |  |  |  |  |  |  |  |  |  |  |  |  |  |  |  |  |  |  |  |  |  |  |  |  |  |  |  |  |  |  |  |  |  |  |  |  |  |  |  |  |  |  |  |  |  |  |  |  |  |  |  |  |
| Leu (L) | 8 | 4.9% |  |  |  |  |  |  |  |  |  |  |  |  |  |  |  |  |  |  |  |  |  |  |  |  |  |  |  |  |  |  |  |  |  |  |  |  |  |  |  |  |  |  |  |  |  |  |  |  |  |  |  |  |  |  |  |  |  |  |  |  |  |  |  |  |  |
| Lys (K) | 20 | 12.3% |  |  |  |  |  |  |  |  |  |  |  |  |  |  |  |  |  |  |  |  |  |  |  |  |  |  |  |  |  |  |  |  |  |  |  |  |  |  |  |  |  |  |  |  |  |  |  |  |  |  |  |  |  |  |  |  |  |  |  |  |  |  |  |  |  |
| Met (M) | 8 | 4.9% |  |  |  |  |  |  |  |  |  |  |  |  |  |  |  |  |  |  |  |  |  |  |  |  |  |  |  |  |  |  |  |  |  |  |  |  |  |  |  |  |  |  |  |  |  |  |  |  |  |  |  |  |  |  |  |  |  |  |  |  |  |  |  |  |  |
| Phe (F) | 13 | 8.0% |  |  |  |  |  |  |  |  |  |  |  |  |  |  |  |  |  |  |  |  |  |  |  |  |  |  |  |  |  |  |  |  |  |  |  |  |  |  |  |  |  |  |  |  |  |  |  |  |  |  |  |  |  |  |  |  |  |  |  |  |  |  |  |  |  |
| Pro (P) | 8 | 4.9% |  |  |  |  |  |  |  |  |  |  |  |  |  |  |  |  |  |  |  |  |  |  |  |  |  |  |  |  |  |  |  |  |  |  |  |  |  |  |  |  |  |  |  |  |  |  |  |  |  |  |  |  |  |  |  |  |  |  |  |  |  |  |  |  |  |
| Ser (S) | 8 | 4.9% |  |  |  |  |  |  |  |  |  |  |  |  |  |  |  |  |  |  |  |  |  |  |  |  |  |  |  |  |  |  |  |  |  |  |  |  |  |  |  |  |  |  |  |  |  |  |  |  |  |  |  |  |  |  |  |  |  |  |  |  |  |  |  |  |  |
| Thr (T) | 14 | 8.6% |  |  |  |  |  |  |  |  |  |  |  |  |  |  |  |  |  |  |  |  |  |  |  |  |  |  |  |  |  |  |  |  |  |  |  |  |  |  |  |  |  |  |  |  |  |  |  |  |  |  |  |  |  |  |  |  |  |  |  |  |  |  |  |  |  |
| Trp (W) | 2 | 1.2% |  |  |  |  |  |  |  |  |  |  |  |  |  |  |  |  |  |  |  |  |  |  |  |  |  |  |  |  |  |  |  |  |  |  |  |  |  |  |  |  |  |  |  |  |  |  |  |  |  |  |  |  |  |  |  |  |  |  |  |  |  |  |  |  |  |
| Tyr (Y) | 2 | 1.2% |  |  |  |  |  |  |  |  |  |  |  |  |  |  |  |  |  |  |  |  |  |  |  |  |  |  |  |  |  |  |  |  |  |  |  |  |  |  |  |  |  |  |  |  |  |  |  |  |  |  |  |  |  |  |  |  |  |  |  |  |  |  |  |  |  |
| Val (V) | 14 | 8.6% |  |  |  |  |  |  |  |  |  |  |  |  |  |  |  |  |  |  |  |  |  |  |  |  |  |  |  |  |  |  |  |  |  |  |  |  |  |  |  |  |  |  |  |  |  |  |  |  |  |  |  |  |  |  |  |  |  |  |  |  |  |  |  |  |  |
| Pyl (O) | 0 | 0.0% |  |  |  |  |  |  |  |  |  |  |  |  |  |  |  |  |  |  |  |  |  |  |  |  |  |  |  |  |  |  |  |  |  |  |  |  |  |  |  |  |  |  |  |  |  |  |  |  |  |  |  |  |  |  |  |  |  |  |  |  |  |  |  |  |  |
| Sec (U) | 0 | 0.0% |  |  |  |  |  |  |  |  |  |  |  |  |  |  |  |  |  |  |  |  |  |  |  |  |  |  |  |  |  |  |  |  |  |  |  |  |  |  |  |  |  |  |  |  |  |  |  |  |  |  |  |  |  |  |  |  |  |  |  |  |  |  |  |  |  |

**uterocalin [Equus przewalskii]**

NCBI Reference Sequence: XP\_070450822.1

**mnllwlamgl illggpha** lh mgpgdpnfde klvkgkwfsv
alasnepkfi akdtdmkffi
hkiqvtpesl qfhfhrkvrq mcvptmmtah ktkkkfqtyv
nhsghtkift ekvdpkhfvi
fcahsmkhgk etvvvtlfsr tptvspdvmw mfkkyckthg
ihasnivdlt qtdrcldharh

**Number of amino acids:** 162**Theoretical pI:** 9.72**Molecular weight:** 18763.00**Amino acid composition:**CSV format

|  |  |  |
| --- | --- | --- |
| Ala (A) | 7 | 4.3% |
| Arg (R) | 5 | 3.1% |
| Asn (N) | 4 | 2.5% |
| Asp (D) | 8 | 4.9% |
| Cys (C) | 4 | 2.5% |
| Gln (Q) | 4 | 2.5% |
| Glu (E) | 5 | 3.1% |
| Gly (G) | 7 | 4.3% |
| His (H) | 14 | 8.6% |
| Ile (I) | 7 | 4.3% |
| Leu (L) | 8 | 4.9% |
| Lys (K) | 20 | 12.3% |
| Met (M) | 8 | 4.9% |
| Phe (F) | 13 | 8.0% |
| Pro (P) | 8 | 4.9% |
| Ser (S) | 8 | 4.9% |
| Thr (T) | 14 | 8.6% |
| Trp (W) | 2 | 1.2% |
| Tyr (Y) | 2 | 1.2% |
| Val (V) | 14 | 8.6% |
| Pyl (O) | 0 | 0.0% |
| Sec (U) | 0 | 0.0% |

**uterocalin isoform X2 [Equus quagga]**

NCBI Reference Sequence: XP\_046538582.1

**mvivnrrgp vtachvarnp llaedkrati kgseplsevp**
**lvslshsclt**

**mnlllamgl illggpha** lh mgpgdpnfde klvkgkwfsv
alasnepkfi akdtdmkffi hkiqvtpesl
qfhfhrkvrq mcvptmmtah ktkkkfqtyv nhsghtkift
ekvdpkhfvi fcahsmkhgk
etvvvtlfsr tptvspdvmw mfkkyckthg ihtsnivdlt
qtdrcldharh

N-terminal extension here excluded finds BLAST matches with –
uterocalin isoform X2 [Equus quagga]
Sequence ID: XP\_046538582.1 – identical - itself
uterocalin isoform X1 [Equus quagga]
Sequence ID: XP\_046538574.1 - identical
uterocalin isoform X2 [Equus asinus]
Sequence ID: XP\_070376020.1 – one substitution
uterocalin isoform X1 [Equus asinus]
Sequence ID: XP\_070376019.1 – one substitution
And a 59% identity match with
uncharacterized protein LOC131490937 [Neofelis nebulosa]
Sequence ID: XP\_058549321.1

**Number of amino acids:** 162**Theoretical pI:** 9.72**Molecular weight:** 18793.02**Amino acid composition:**CSV format

|  |  |  |
| --- | --- | --- |
| Ala (A) | 6 | 3.7% |
| Arg (R) | 5 | 3.1% |
| Asn (N) | 4 | 2.5% |
| Asp (D) | 8 | 4.9% |
| Cys (C) | 4 | 2.5% |
| Gln (Q) | 4 | 2.5% |
| Glu (E) | 5 | 3.1% |
| Gly (G) | 7 | 4.3% |
| His (H) | 14 | 8.6% |
| Ile (I) | 7 | 4.3% |
| Leu (L) | 8 | 4.9% |
| Lys (K) | 20 | 12.3% |
| Met (M) | 8 | 4.9% |
| Phe (F) | 13 | 8.0% |
| Pro (P) | 8 | 4.9% |
| Ser (S) | 8 | 4.9% |
| Thr (T) | 15 | 9.3% |
| Trp (W) | 2 | 1.2% |
| Tyr (Y) | 2 | 1.2% |
| Val (V) | 14 | 8.6% |
| Pyl (O) | 0 | 0.0% |
| Sec (U) | 0 | 0.0% |

**epididymal-specific lipocalin-9 [Diceror bicornis minor] Southern black rhinoceros**

NCBI Reference Sequence: XP\_058380674.1

msllllamgl tllgglqa lh kgpgdpnfde klvkgkwfsv
amasnepkfi tkdtdmkffi
hniqvtpksl qlhvhrkvkg vcvptmtan ktgkkfytv
nhsgkhkifl ekvdpqhfa
fcthsvkghgk emmvvnlftr tptvspdvlw mfrkyckthg
ihstnivdlt qtdrclhahq

wqkapfpwld hlggphaacp fhvlsiqrw ssgspnhsth
yvllpgnrpf wrretvalrt
qeevtlklp qtpktkpvit dvkasdslln avkannsgpa
vtgpgqegg gggtavp

C-terminal extension finds BLAST matches with –
epididymal-specific lipocalin-9 [Diceros bicornis minor]

Sequence ID: XP\_058380674.1 – itself

epididymal-specific lipocalin-9 [Ceratotherium simum simum]

Sequence ID: XP\_014637465.1 – 75% identities

Number of amino acids: 162

Theoretical pI: 9.72

Molecular weight: 18610.68

Amino acid composition: CSV format

|  |  |  |
| --- | --- | --- |
| Ala (A) | 5 | 3.1% |
| Arg (R) | 4 | 2.5% |
| Asn (N) | 7 | 4.3% |
| Asp (D) | 8 | 4.9% |
| Cys (C) | 4 | 2.5% |
| Gln (Q) | 6 | 3.7% |
| Glu (E) | 4 | 2.5% |
| Gly (G) | 8 | 4.9% |
| His (H) | 13 | 8.0% |
| Ile (I) | 7 | 4.3% |
| Leu (L) | 9 | 5.6% |
| Lys (K) | 20 | 12.3% |
| Met (M) | 6 | 3.7% |
| Phe (F) | 11 | 6.8% |
| Pro (P) | 8 | 4.9% |
| Ser (S) | 8 | 4.9% |
| Thr (T) | 15 | 9.3% |
| Trp (W) | 2 | 1.2% |
| Tyr (Y) | 2 | 1.2% |
| Val (V) | 15 | 9.3% |
| Pyl (O) | 0 | 0.0% |
| Sec (U) | 0 | 0.0% |

**epididymal-specific lipocalin-9 [Ceratotherium simum simum] southern white rhinoceros**

NCBI Reference Sequence: XP\_014637465.1

msllllavgl tllgglqa lh kgpgdpnfde klvkgkwfsv
amasnepkfi tkdtdmkffi
hniqvtpksl qlhvhrkvkg vcvptmtan ktgkkfytv
nhsskhmifl ekvdpqhfa
fcthsvkghgk etmvvnlftr tptvspdvlw mfrkyckthg
ihstnivdlt qtdrcpphrq

pqeaeglpgd acsahlaplc sslhrslspr ppvdsthyvl
lpgnrpfwwg ervtlrtqge
vtlklpqt ktkpvitdvk asdsllnavk annsgpavqt
gpgrvvgrql fpeppwllpa
301 paaatataeg petaeahsps lpqggnpals gccrdrdkpv
stasglqvqp tggpgpgqwn
3 ptapshrlsp plpgpafnqp llspqa

Extended C-terminal sequence BLAST matches to –
epididymal-specific lipocalin-9 [Ceratotherium simum simum]

Sequence ID: XP\_014637465.1 – itself

epididymal-specific lipocalin-9 [Diceror bicornis minor]

Sequence ID: XP\_058380674.1 – 77% identities
with large blocks of identical

Number of amino acids: 162

Theoretical pI: 9.83

Molecular weight: 18584.64

Amino acid composition: CSV format

|  |  |  |
| --- | --- | --- |
| Ala (A) | 4 | 2.5% |
| Arg (R) | 5 | 3.1% |
| Asn (N) | 7 | 4.3% |
| Asp (D) | 7 | 4.3% |
| Cys (C) | 4 | 2.5% |
| Gln (Q) | 6 | 3.7% |
| Glu (E) | 4 | 2.5% |
| Gly (G) | 8 | 4.9% |
| His (H) | 12 | 7.4% |
| Ile (I) | 7 | 4.3% |
| Leu (L) | 8 | 4.9% |
| Lys (K) | 19 | 11.7% |
| Met (M) | 6 | 3.7% |
| Phe (F) | 11 | 6.8% |
| Pro (P) | 10 | 6.2% |
| Ser (S) | 9 | 5.6% |
| Thr (T) | 16 | 9.9% |
| Trp (W) | 2 | 1.2% |
| Tyr (Y) | 2 | 1.2% |
| Val (V) | 15 | 9.3% |
| Pyl (O) | 0 | 0.0% |
| Sec (U) | 0 | 0.0% |

|  |  |  |  |  |  |  |  |  |  |  |  |  |  |  |  |  |  |  |  |  |  |  |  |  |  |  |  |  |  |  |  |  |  |  |  |  |  |  |  |  |  |  |  |  |  |  |  |  |  |  |  |  |  |  |  |  |  |  |  |  |  |  |  |  |  |  |  |
| --- | --- | --- | --- | --- | --- | --- | --- | --- | --- | --- | --- | --- | --- | --- | --- | --- | --- | --- | --- | --- | --- | --- | --- | --- | --- | --- | --- | --- | --- | --- | --- | --- | --- | --- | --- | --- | --- | --- | --- | --- | --- | --- | --- | --- | --- | --- | --- | --- | --- | --- | --- | --- | --- | --- | --- | --- | --- | --- | --- | --- | --- | --- | --- | --- | --- | --- | --- |
| Uterocalin [Tapirus indicus]<br>Tapir indicus genome reference<br>Tapirus_indicus_015923-RA<br>MNVLLLA**VGLTLLGGLQA**<br>LHMGPGDPNPFDEKLIKGWYSVALGSNEPKFLTKDTD<br>MKFFIHKIQVTPKSLQFHFHKKVKGMCVPTMMTVNKT<br>GKKFQYTVNHFGHKMIFLEKVDQHFVIFCTHSMKHG<br>KETMVVNLFSTRPTVSPDILWMFKNYCKSHGIHSTNII<br>DLTQTDRCLHARQ | Number of amino acids: 162<br>Theoretical pI: 9.62<br>Molecular weight: 18854.12<br>Amino acid composition:<br>CSV format<br>Ala (A)<br>2<br>1.2%<br>Arg (R)<br>3<br>1.9%<br>Asn (N)<br>7<br>4.3%<br>Asp (D)<br>8<br>4.9%<br>Cys (C)<br>4<br>2.5%<br>Gln (Q)<br>6<br>3.7%<br>Glu (E)<br>4<br>2.5%<br>Gly (G)<br>9<br>5.6%<br>His (H)<br>12<br>7.4%<br>Ile (I)<br>9<br>5.6%<br>Leu (L)<br>10<br>6.2%<br>Lys (K)<br>20<br>12.3%<br>Met (M)<br>9<br>5.6%<br>Phe (F)<br>13<br>8.0%<br>Pro (P)<br>8<br>4.9%<br>Ser (S)<br>8<br>4.9%<br>Thr (T)<br>14<br>8.6%<br>Trp (W)<br>2<br>1.2%<br>Tyr (Y)<br>3<br>1.9%<br>Val (V)<br>11<br>6.8%<br>Pyl (O)<br>0<br>0.0%<br>Sec (U)<br>0<br>0.0% | Ala (A) | 2 | 1.2% | Arg (R) | 3 | 1.9% | Asn (N) | 7 | 4.3% | Asp (D) | 8 | 4.9% | Cys (C) | 4 | 2.5% | Gln (Q) | 6 | 3.7% | Glu (E) | 4 | 2.5% | Gly (G) | 9 | 5.6% | His (H) | 12 | 7.4% | Ile (I) | 9 | 5.6% | Leu (L) | 10 | 6.2% | Lys (K) | 20 | 12.3% | Met (M) | 9 | 5.6% | Phe (F) | 13 | 8.0% | Pro (P) | 8 | 4.9% | Ser (S) | 8 | 4.9% | Thr (T) | 14 | 8.6% | Trp (W) | 2 | 1.2% | Tyr (Y) | 3 | 1.9% | Val (V) | 11 | 6.8% | Pyl (O) | 0 | 0.0% | Sec (U) | 0 | 0.0% |
| Ala (A) | 2 | 1.2% |  |  |  |  |  |  |  |  |  |  |  |  |  |  |  |  |  |  |  |  |  |  |  |  |  |  |  |  |  |  |  |  |  |  |  |  |  |  |  |  |  |  |  |  |  |  |  |  |  |  |  |  |  |  |  |  |  |  |  |  |  |  |  |  |  |
| Arg (R) | 3 | 1.9% |  |  |  |  |  |  |  |  |  |  |  |  |  |  |  |  |  |  |  |  |  |  |  |  |  |  |  |  |  |  |  |  |  |  |  |  |  |  |  |  |  |  |  |  |  |  |  |  |  |  |  |  |  |  |  |  |  |  |  |  |  |  |  |  |  |
| Asn (N) | 7 | 4.3% |  |  |  |  |  |  |  |  |  |  |  |  |  |  |  |  |  |  |  |  |  |  |  |  |  |  |  |  |  |  |  |  |  |  |  |  |  |  |  |  |  |  |  |  |  |  |  |  |  |  |  |  |  |  |  |  |  |  |  |  |  |  |  |  |  |
| Asp (D) | 8 | 4.9% |  |  |  |  |  |  |  |  |  |  |  |  |  |  |  |  |  |  |  |  |  |  |  |  |  |  |  |  |  |  |  |  |  |  |  |  |  |  |  |  |  |  |  |  |  |  |  |  |  |  |  |  |  |  |  |  |  |  |  |  |  |  |  |  |  |
| Cys (C) | 4 | 2.5% |  |  |  |  |  |  |  |  |  |  |  |  |  |  |  |  |  |  |  |  |  |  |  |  |  |  |  |  |  |  |  |  |  |  |  |  |  |  |  |  |  |  |  |  |  |  |  |  |  |  |  |  |  |  |  |  |  |  |  |  |  |  |  |  |  |
| Gln (Q) | 6 | 3.7% |  |  |  |  |  |  |  |  |  |  |  |  |  |  |  |  |  |  |  |  |  |  |  |  |  |  |  |  |  |  |  |  |  |  |  |  |  |  |  |  |  |  |  |  |  |  |  |  |  |  |  |  |  |  |  |  |  |  |  |  |  |  |  |  |  |
| Glu (E) | 4 | 2.5% |  |  |  |  |  |  |  |  |  |  |  |  |  |  |  |  |  |  |  |  |  |  |  |  |  |  |  |  |  |  |  |  |  |  |  |  |  |  |  |  |  |  |  |  |  |  |  |  |  |  |  |  |  |  |  |  |  |  |  |  |  |  |  |  |  |
| Gly (G) | 9 | 5.6% |  |  |  |  |  |  |  |  |  |  |  |  |  |  |  |  |  |  |  |  |  |  |  |  |  |  |  |  |  |  |  |  |  |  |  |  |  |  |  |  |  |  |  |  |  |  |  |  |  |  |  |  |  |  |  |  |  |  |  |  |  |  |  |  |  |
| His (H) | 12 | 7.4% |  |  |  |  |  |  |  |  |  |  |  |  |  |  |  |  |  |  |  |  |  |  |  |  |  |  |  |  |  |  |  |  |  |  |  |  |  |  |  |  |  |  |  |  |  |  |  |  |  |  |  |  |  |  |  |  |  |  |  |  |  |  |  |  |  |
| Ile (I) | 9 | 5.6% |  |  |  |  |  |  |  |  |  |  |  |  |  |  |  |  |  |  |  |  |  |  |  |  |  |  |  |  |  |  |  |  |  |  |  |  |  |  |  |  |  |  |  |  |  |  |  |  |  |  |  |  |  |  |  |  |  |  |  |  |  |  |  |  |  |
| Leu (L) | 10 | 6.2% |  |  |  |  |  |  |  |  |  |  |  |  |  |  |  |  |  |  |  |  |  |  |  |  |  |  |  |  |  |  |  |  |  |  |  |  |  |  |  |  |  |  |  |  |  |  |  |  |  |  |  |  |  |  |  |  |  |  |  |  |  |  |  |  |  |
| Lys (K) | 20 | 12.3% |  |  |  |  |  |  |  |  |  |  |  |  |  |  |  |  |  |  |  |  |  |  |  |  |  |  |  |  |  |  |  |  |  |  |  |  |  |  |  |  |  |  |  |  |  |  |  |  |  |  |  |  |  |  |  |  |  |  |  |  |  |  |  |  |  |
| Met (M) | 9 | 5.6% |  |  |  |  |  |  |  |  |  |  |  |  |  |  |  |  |  |  |  |  |  |  |  |  |  |  |  |  |  |  |  |  |  |  |  |  |  |  |  |  |  |  |  |  |  |  |  |  |  |  |  |  |  |  |  |  |  |  |  |  |  |  |  |  |  |
| Phe (F) | 13 | 8.0% |  |  |  |  |  |  |  |  |  |  |  |  |  |  |  |  |  |  |  |  |  |  |  |  |  |  |  |  |  |  |  |  |  |  |  |  |  |  |  |  |  |  |  |  |  |  |  |  |  |  |  |  |  |  |  |  |  |  |  |  |  |  |  |  |  |
| Pro (P) | 8 | 4.9% |  |  |  |  |  |  |  |  |  |  |  |  |  |  |  |  |  |  |  |  |  |  |  |  |  |  |  |  |  |  |  |  |  |  |  |  |  |  |  |  |  |  |  |  |  |  |  |  |  |  |  |  |  |  |  |  |  |  |  |  |  |  |  |  |  |
| Ser (S) | 8 | 4.9% |  |  |  |  |  |  |  |  |  |  |  |  |  |  |  |  |  |  |  |  |  |  |  |  |  |  |  |  |  |  |  |  |  |  |  |  |  |  |  |  |  |  |  |  |  |  |  |  |  |  |  |  |  |  |  |  |  |  |  |  |  |  |  |  |  |
| Thr (T) | 14 | 8.6% |  |  |  |  |  |  |  |  |  |  |  |  |  |  |  |  |  |  |  |  |  |  |  |  |  |  |  |  |  |  |  |  |  |  |  |  |  |  |  |  |  |  |  |  |  |  |  |  |  |  |  |  |  |  |  |  |  |  |  |  |  |  |  |  |  |
| Trp (W) | 2 | 1.2% |  |  |  |  |  |  |  |  |  |  |  |  |  |  |  |  |  |  |  |  |  |  |  |  |  |  |  |  |  |  |  |  |  |  |  |  |  |  |  |  |  |  |  |  |  |  |  |  |  |  |  |  |  |  |  |  |  |  |  |  |  |  |  |  |  |
| Tyr (Y) | 3 | 1.9% |  |  |  |  |  |  |  |  |  |  |  |  |  |  |  |  |  |  |  |  |  |  |  |  |  |  |  |  |  |  |  |  |  |  |  |  |  |  |  |  |  |  |  |  |  |  |  |  |  |  |  |  |  |  |  |  |  |  |  |  |  |  |  |  |  |
| Val (V) | 11 | 6.8% |  |  |  |  |  |  |  |  |  |  |  |  |  |  |  |  |  |  |  |  |  |  |  |  |  |  |  |  |  |  |  |  |  |  |  |  |  |  |  |  |  |  |  |  |  |  |  |  |  |  |  |  |  |  |  |  |  |  |  |  |  |  |  |  |  |
| Pyl (O) | 0 | 0.0% |  |  |  |  |  |  |  |  |  |  |  |  |  |  |  |  |  |  |  |  |  |  |  |  |  |  |  |  |  |  |  |  |  |  |  |  |  |  |  |  |  |  |  |  |  |  |  |  |  |  |  |  |  |  |  |  |  |  |  |  |  |  |  |  |  |
| Sec (U) | 0 | 0.0% |  |  |  |  |  |  |  |  |  |  |  |  |  |  |  |  |  |  |  |  |  |  |  |  |  |  |  |  |  |  |  |  |  |  |  |  |  |  |  |  |  |  |  |  |  |  |  |  |  |  |  |  |  |  |  |  |  |  |  |  |  |  |  |  |  |
| Uterocalin-like proteins<br>(by sequence similarity) |  |  |  |  |  |  |  |  |  |  |  |  |  |  |  |  |  |  |  |  |  |  |  |  |  |  |  |  |  |  |  |  |  |  |  |  |  |  |  |  |  |  |  |  |  |  |  |  |  |  |  |  |  |  |  |  |  |  |  |  |  |  |  |  |  |  |  |
| uterocalin isoform X2 [Camelus ferus]<br>NCBI Reference Sequence: XP_006172841.1<br>msllllatgl tllgspqt lh qgpqdsnfne tlvsqdwfsa<br>alasnrrprll qegadaqlfi<br>hsiqvtpral qlhlhrkvng tcvpvmtnvan ktkrkfqylm<br>eyggqkrifl eevdpksyvi<br>fcthhkehkgk etvvvtlfsr tpkvtqdtll ifknyckshg<br>ihktniinvk kadhclharq<br>Use this one – X1 and X2 are identical until position 140 and then don't match at all. X2 aligns well with horse uterocalin along the whole length. So, X1 is deviant for last 40 amino acids. | Number of amino acids: 162<br>Theoretical pI: 9.48<br>Molecular weight: 18503.29<br>Amino acid composition:<br>CSV format<br>Ala (A)<br>9<br>5.6%<br>Arg (R)<br>8<br>4.9%<br>Asn (N)<br>8<br>4.9%<br>Asp (D)<br>6<br>3.7%<br>Cys (C)<br>4<br>2.5%<br>Gln (Q)<br>10<br>6.2%<br>Glu (E)<br>7<br>4.3%<br>Gly (G)<br>8<br>4.9%<br>His (H)<br>11<br>6.8%<br>Ile (I)<br>8<br>4.9%<br>Leu (L)<br>15<br>9.3%<br>Lys (K)<br>13<br>8.0%<br>Met (M)<br>2<br>1.2%<br>Phe (F)<br>8<br>4.9%<br>Pro (P)<br>6<br>3.7%<br>Ser (S)<br>8<br>4.9%<br>Thr (T)<br>13<br>8.0%<br>Trp (W)<br>1<br>0.6%<br>Tyr (Y)<br>4<br>2.5%<br>Val (V)<br>13<br>8.0%<br>Pyl (O)<br>0<br>0.0%<br>Sec (U)<br>0<br>0.0% | Ala (A) | 9 | 5.6% | Arg (R) | 8 | 4.9% | Asn (N) | 8 | 4.9% | Asp (D) | 6 | 3.7% | Cys (C) | 4 | 2.5% | Gln (Q) | 10 | 6.2% | Glu (E) | 7 | 4.3% | Gly (G) | 8 | 4.9% | His (H) | 11 | 6.8% | Ile (I) | 8 | 4.9% | Leu (L) | 15 | 9.3% | Lys (K) | 13 | 8.0% | Met (M) | 2 | 1.2% | Phe (F) | 8 | 4.9% | Pro (P) | 6 | 3.7% | Ser (S) | 8 | 4.9% | Thr (T) | 13 | 8.0% | Trp (W) | 1 | 0.6% | Tyr (Y) | 4 | 2.5% | Val (V) | 13 | 8.0% | Pyl (O) | 0 | 0.0% | Sec (U) | 0 | 0.0% |
| Ala (A) | 9 | 5.6% |  |  |  |  |  |  |  |  |  |  |  |  |  |  |  |  |  |  |  |  |  |  |  |  |  |  |  |  |  |  |  |  |  |  |  |  |  |  |  |  |  |  |  |  |  |  |  |  |  |  |  |  |  |  |  |  |  |  |  |  |  |  |  |  |  |
| Arg (R) | 8 | 4.9% |  |  |  |  |  |  |  |  |  |  |  |  |  |  |  |  |  |  |  |  |  |  |  |  |  |  |  |  |  |  |  |  |  |  |  |  |  |  |  |  |  |  |  |  |  |  |  |  |  |  |  |  |  |  |  |  |  |  |  |  |  |  |  |  |  |
| Asn (N) | 8 | 4.9% |  |  |  |  |  |  |  |  |  |  |  |  |  |  |  |  |  |  |  |  |  |  |  |  |  |  |  |  |  |  |  |  |  |  |  |  |  |  |  |  |  |  |  |  |  |  |  |  |  |  |  |  |  |  |  |  |  |  |  |  |  |  |  |  |  |
| Asp (D) | 6 | 3.7% |  |  |  |  |  |  |  |  |  |  |  |  |  |  |  |  |  |  |  |  |  |  |  |  |  |  |  |  |  |  |  |  |  |  |  |  |  |  |  |  |  |  |  |  |  |  |  |  |  |  |  |  |  |  |  |  |  |  |  |  |  |  |  |  |  |
| Cys (C) | 4 | 2.5% |  |  |  |  |  |  |  |  |  |  |  |  |  |  |  |  |  |  |  |  |  |  |  |  |  |  |  |  |  |  |  |  |  |  |  |  |  |  |  |  |  |  |  |  |  |  |  |  |  |  |  |  |  |  |  |  |  |  |  |  |  |  |  |  |  |
| Gln (Q) | 10 | 6.2% |  |  |  |  |  |  |  |  |  |  |  |  |  |  |  |  |  |  |  |  |  |  |  |  |  |  |  |  |  |  |  |  |  |  |  |  |  |  |  |  |  |  |  |  |  |  |  |  |  |  |  |  |  |  |  |  |  |  |  |  |  |  |  |  |  |
| Glu (E) | 7 | 4.3% |  |  |  |  |  |  |  |  |  |  |  |  |  |  |  |  |  |  |  |  |  |  |  |  |  |  |  |  |  |  |  |  |  |  |  |  |  |  |  |  |  |  |  |  |  |  |  |  |  |  |  |  |  |  |  |  |  |  |  |  |  |  |  |  |  |
| Gly (G) | 8 | 4.9% |  |  |  |  |  |  |  |  |  |  |  |  |  |  |  |  |  |  |  |  |  |  |  |  |  |  |  |  |  |  |  |  |  |  |  |  |  |  |  |  |  |  |  |  |  |  |  |  |  |  |  |  |  |  |  |  |  |  |  |  |  |  |  |  |  |
| His (H) | 11 | 6.8% |  |  |  |  |  |  |  |  |  |  |  |  |  |  |  |  |  |  |  |  |  |  |  |  |  |  |  |  |  |  |  |  |  |  |  |  |  |  |  |  |  |  |  |  |  |  |  |  |  |  |  |  |  |  |  |  |  |  |  |  |  |  |  |  |  |
| Ile (I) | 8 | 4.9% |  |  |  |  |  |  |  |  |  |  |  |  |  |  |  |  |  |  |  |  |  |  |  |  |  |  |  |  |  |  |  |  |  |  |  |  |  |  |  |  |  |  |  |  |  |  |  |  |  |  |  |  |  |  |  |  |  |  |  |  |  |  |  |  |  |
| Leu (L) | 15 | 9.3% |  |  |  |  |  |  |  |  |  |  |  |  |  |  |  |  |  |  |  |  |  |  |  |  |  |  |  |  |  |  |  |  |  |  |  |  |  |  |  |  |  |  |  |  |  |  |  |  |  |  |  |  |  |  |  |  |  |  |  |  |  |  |  |  |  |
| Lys (K) | 13 | 8.0% |  |  |  |  |  |  |  |  |  |  |  |  |  |  |  |  |  |  |  |  |  |  |  |  |  |  |  |  |  |  |  |  |  |  |  |  |  |  |  |  |  |  |  |  |  |  |  |  |  |  |  |  |  |  |  |  |  |  |  |  |  |  |  |  |  |
| Met (M) | 2 | 1.2% |  |  |  |  |  |  |  |  |  |  |  |  |  |  |  |  |  |  |  |  |  |  |  |  |  |  |  |  |  |  |  |  |  |  |  |  |  |  |  |  |  |  |  |  |  |  |  |  |  |  |  |  |  |  |  |  |  |  |  |  |  |  |  |  |  |
| Phe (F) | 8 | 4.9% |  |  |  |  |  |  |  |  |  |  |  |  |  |  |  |  |  |  |  |  |  |  |  |  |  |  |  |  |  |  |  |  |  |  |  |  |  |  |  |  |  |  |  |  |  |  |  |  |  |  |  |  |  |  |  |  |  |  |  |  |  |  |  |  |  |
| Pro (P) | 6 | 3.7% |  |  |  |  |  |  |  |  |  |  |  |  |  |  |  |  |  |  |  |  |  |  |  |  |  |  |  |  |  |  |  |  |  |  |  |  |  |  |  |  |  |  |  |  |  |  |  |  |  |  |  |  |  |  |  |  |  |  |  |  |  |  |  |  |  |
| Ser (S) | 8 | 4.9% |  |  |  |  |  |  |  |  |  |  |  |  |  |  |  |  |  |  |  |  |  |  |  |  |  |  |  |  |  |  |  |  |  |  |  |  |  |  |  |  |  |  |  |  |  |  |  |  |  |  |  |  |  |  |  |  |  |  |  |  |  |  |  |  |  |
| Thr (T) | 13 | 8.0% |  |  |  |  |  |  |  |  |  |  |  |  |  |  |  |  |  |  |  |  |  |  |  |  |  |  |  |  |  |  |  |  |  |  |  |  |  |  |  |  |  |  |  |  |  |  |  |  |  |  |  |  |  |  |  |  |  |  |  |  |  |  |  |  |  |
| Trp (W) | 1 | 0.6% |  |  |  |  |  |  |  |  |  |  |  |  |  |  |  |  |  |  |  |  |  |  |  |  |  |  |  |  |  |  |  |  |  |  |  |  |  |  |  |  |  |  |  |  |  |  |  |  |  |  |  |  |  |  |  |  |  |  |  |  |  |  |  |  |  |
| Tyr (Y) | 4 | 2.5% |  |  |  |  |  |  |  |  |  |  |  |  |  |  |  |  |  |  |  |  |  |  |  |  |  |  |  |  |  |  |  |  |  |  |  |  |  |  |  |  |  |  |  |  |  |  |  |  |  |  |  |  |  |  |  |  |  |  |  |  |  |  |  |  |  |
| Val (V) | 13 | 8.0% |  |  |  |  |  |  |  |  |  |  |  |  |  |  |  |  |  |  |  |  |  |  |  |  |  |  |  |  |  |  |  |  |  |  |  |  |  |  |  |  |  |  |  |  |  |  |  |  |  |  |  |  |  |  |  |  |  |  |  |  |  |  |  |  |  |
| Pyl (O) | 0 | 0.0% |  |  |  |  |  |  |  |  |  |  |  |  |  |  |  |  |  |  |  |  |  |  |  |  |  |  |  |  |  |  |  |  |  |  |  |  |  |  |  |  |  |  |  |  |  |  |  |  |  |  |  |  |  |  |  |  |  |  |  |  |  |  |  |  |  |
| Sec (U) | 0 | 0.0% |  |  |  |  |  |  |  |  |  |  |  |  |  |  |  |  |  |  |  |  |  |  |  |  |  |  |  |  |  |  |  |  |  |  |  |  |  |  |  |  |  |  |  |  |  |  |  |  |  |  |  |  |  |  |  |  |  |  |  |  |  |  |  |  |  |

**uterocalin isoform X1 [Camelus ferus]**

NCBI Reference Sequence: XP\_032333258.1

mslllatgl tllgspqt lh qgpqdsnfne tlvsqdwfsa
alasnrprll qegadaqlfi
hsiqvtpral qlhlhrkvng tcvpvmtnvan ktkrkfqylm
eyggqkrifl eevdpksyvi
fcthhkehkgk etvvvtlfsk crrphppghh phppdragga
prqghparcg spgktpghsq

qdpggtpgtw rplgsrraln tr

This one not used – see note for X2 above.

**Number of amino acids:** 162**Theoretical pI:** 9.73**Molecular weight:** 18144.65**Amino acid composition:** CSV format

|  |  |  |
| --- | --- | --- |
| Ala (A) | 10 | 6.2% |
| Arg (R) | 11 | 6.8% |
| Asn (N) | 5 | 3.1% |
| Asp (D) | 5 | 3.1% |
| Cys (C) | 4 | 2.5% |
| Gln (Q) | 11 | 6.8% |
| Glu (E) | 7 | 4.3% |
| Gly (G) | 13 | 8.0% |
| His (H) | 13 | 8.0% |
| Ile (I) | 4 | 2.5% |
| Leu (L) | 12 | 7.4% |
| Lys (K) | 10 | 6.2% |
| Met (M) | 2 | 1.2% |
| Phe (F) | 7 | 4.3% |
| Pro (P) | 15 | 9.3% |
| Ser (S) | 9 | 5.6% |
| Thr (T) | 9 | 5.6% |
| Trp (W) | 1 | 0.6% |
| Tyr (Y) | 3 | 1.9% |
| Val (V) | 11 | 6.8% |
| Pyl (O) | 0 | 0.0% |
| Sec (U) | 0 | 0.0% |

**uterocalin [Camelus dromedarius]**

NCBI Reference Sequence: XP\_031305434.2

mslllatgl tllgspqt lh qgpqdsnfne tlvsqdwfsa
alasnrprll qegadaqlfi
hsiqvtpral qlhlhrkvng tcvpvmtnvan ktkrkfqylm
eyggqkrifl eevdpksyvi
fcthhkeqgk etvvvtlfsr tpkvtqdtll ifknyckshg
ihktniinvk kadhclharq

sapsapipip vppsplsitg vsprhsatfp egpedlgqpe
epfslpdplg aattlleaf
gvsnkrlprt qcalgaspvh epgw

Only one substitution with Camelus ferus.

BLAST search finds –

**uterocalin [Camelus dromedarius]**

**Sequence ID:** XP\_031305434.2 itself**protein FAM193B-like [Balaenoptera acutorostrata]**

**Sequence ID:** XP\_007194825.2 - 57% with short blocks of identical**uterocalin-like [Balaenoptera physalus]**

**Sequence ID:** XP\_082325636.1 – 53% with short blocks of identical**hypothetical protein E2I00\_001780, partial [Balaenoptera physalus]**

**Number of amino acids:** 162**Theoretical pI:** 9.48**Molecular weight:** 18494.27**Amino acid composition:** CSV format

|  |  |  |
| --- | --- | --- |
| Ala (A) | 9 | 5.6% |
| Arg (R) | 8 | 4.9% |
| Asn (N) | 8 | 4.9% |
| Asp (D) | 6 | 3.7% |
| Cys (C) | 4 | 2.5% |
| Gln (Q) | 11 | 6.8% |
| Glu (E) | 7 | 4.3% |
| Gly (G) | 8 | 4.9% |
| His (H) | 10 | 6.2% |
| Ile (I) | 8 | 4.9% |
| Leu (L) | 15 | 9.3% |
| Lys (K) | 13 | 8.0% |
| Met (M) | 2 | 1.2% |
| Phe (F) | 8 | 4.9% |
| Pro (P) | 6 | 3.7% |
| Ser (S) | 8 | 4.9% |
| Thr (T) | 13 | 8.0% |
| Trp (W) | 1 | 0.6% |
| Tyr (Y) | 4 | 2.5% |
| Val (V) | 13 | 8.0% |
| Pyl (O) | 0 | 0.0% |
| Sec (U) | 0 | 0.0% |

|  |  |  |  |  |  |  |  |  |  |  |  |  |  |  |  |  |  |  |  |  |  |  |  |  |  |  |  |  |  |  |  |  |  |  |  |  |  |  |  |  |  |  |  |  |  |  |  |  |  |  |  |  |  |  |  |  |  |  |  |  |  |  |  |  |  |  |  |
| --- | --- | --- | --- | --- | --- | --- | --- | --- | --- | --- | --- | --- | --- | --- | --- | --- | --- | --- | --- | --- | --- | --- | --- | --- | --- | --- | --- | --- | --- | --- | --- | --- | --- | --- | --- | --- | --- | --- | --- | --- | --- | --- | --- | --- | --- | --- | --- | --- | --- | --- | --- | --- | --- | --- | --- | --- | --- | --- | --- | --- | --- | --- | --- | --- | --- | --- | --- |
| Sequence ID: KAB0394076.1 – 52% identical with short blocks identical |  |  |  |  |  |  |  |  |  |  |  |  |  |  |  |  |  |  |  |  |  |  |  |  |  |  |  |  |  |  |  |  |  |  |  |  |  |  |  |  |  |  |  |  |  |  |  |  |  |  |  |  |  |  |  |  |  |  |  |  |  |  |  |  |  |  |  |
| uterocalin-like isoform X1 [Zalophus californianus] California sea lion<br>NCBI Reference Sequence: XP_027471829.1<br>mntprlvvl allggpha<br>lh tgprdp side nmvngdwfsi<br>aqassepkll wkdshmmffv<br>rkihvs lkti efhlyrriqg mcvpiimman ktkkkf qytI<br>kyaghnvifl eevdpsrfli<br>fcihnhwhgk etvvvnllsr tppaswdimq tfknyckshg<br>ispaniinlt rtdrcIha | Number of amino acids: 160<br>Theoretical pI: 9.39<br>Molecular weight: 18668.78<br>Amino acid composition:<br>CSV format<br>Ala (A)<br>7<br>4.4%<br>Arg (R)<br>8<br>5.0%<br>Asn (N)<br>9<br>5.6%<br>Asp (D)<br>7<br>4.4%<br>Cys (C)<br>4<br>2.5%<br>Gln (Q)<br>4<br>2.5%<br>Glu (E)<br>6<br>3.8%<br>Gly (G)<br>6<br>3.8%<br>His (H)<br>10<br>6.2%<br>Ile (I)<br>14<br>8.8%<br>Leu (L)<br>12<br>7.5%<br>Lys (K)<br>12<br>7.5%<br>Met (M)<br>7<br>4.4%<br>Phe (F)<br>9<br>5.6%<br>Pro (P)<br>8<br>5.0%<br>Ser (S)<br>11<br>6.9%<br>Thr (T)<br>9<br>5.6%<br>Trp (W)<br>4<br>2.5%<br>Tyr (Y)<br>4<br>2.5%<br>Val (V)<br>9<br>5.6%<br>Pyl (O)<br>0<br>0.0%<br>Sec (U)<br>0<br>0.0% | Ala (A) | 7 | 4.4% | Arg (R) | 8 | 5.0% | Asn (N) | 9 | 5.6% | Asp (D) | 7 | 4.4% | Cys (C) | 4 | 2.5% | Gln (Q) | 4 | 2.5% | Glu (E) | 6 | 3.8% | Gly (G) | 6 | 3.8% | His (H) | 10 | 6.2% | Ile (I) | 14 | 8.8% | Leu (L) | 12 | 7.5% | Lys (K) | 12 | 7.5% | Met (M) | 7 | 4.4% | Phe (F) | 9 | 5.6% | Pro (P) | 8 | 5.0% | Ser (S) | 11 | 6.9% | Thr (T) | 9 | 5.6% | Trp (W) | 4 | 2.5% | Tyr (Y) | 4 | 2.5% | Val (V) | 9 | 5.6% | Pyl (O) | 0 | 0.0% | Sec (U) | 0 | 0.0% |
| Ala (A) | 7 | 4.4% |  |  |  |  |  |  |  |  |  |  |  |  |  |  |  |  |  |  |  |  |  |  |  |  |  |  |  |  |  |  |  |  |  |  |  |  |  |  |  |  |  |  |  |  |  |  |  |  |  |  |  |  |  |  |  |  |  |  |  |  |  |  |  |  |  |
| Arg (R) | 8 | 5.0% |  |  |  |  |  |  |  |  |  |  |  |  |  |  |  |  |  |  |  |  |  |  |  |  |  |  |  |  |  |  |  |  |  |  |  |  |  |  |  |  |  |  |  |  |  |  |  |  |  |  |  |  |  |  |  |  |  |  |  |  |  |  |  |  |  |
| Asn (N) | 9 | 5.6% |  |  |  |  |  |  |  |  |  |  |  |  |  |  |  |  |  |  |  |  |  |  |  |  |  |  |  |  |  |  |  |  |  |  |  |  |  |  |  |  |  |  |  |  |  |  |  |  |  |  |  |  |  |  |  |  |  |  |  |  |  |  |  |  |  |
| Asp (D) | 7 | 4.4% |  |  |  |  |  |  |  |  |  |  |  |  |  |  |  |  |  |  |  |  |  |  |  |  |  |  |  |  |  |  |  |  |  |  |  |  |  |  |  |  |  |  |  |  |  |  |  |  |  |  |  |  |  |  |  |  |  |  |  |  |  |  |  |  |  |
| Cys (C) | 4 | 2.5% |  |  |  |  |  |  |  |  |  |  |  |  |  |  |  |  |  |  |  |  |  |  |  |  |  |  |  |  |  |  |  |  |  |  |  |  |  |  |  |  |  |  |  |  |  |  |  |  |  |  |  |  |  |  |  |  |  |  |  |  |  |  |  |  |  |
| Gln (Q) | 4 | 2.5% |  |  |  |  |  |  |  |  |  |  |  |  |  |  |  |  |  |  |  |  |  |  |  |  |  |  |  |  |  |  |  |  |  |  |  |  |  |  |  |  |  |  |  |  |  |  |  |  |  |  |  |  |  |  |  |  |  |  |  |  |  |  |  |  |  |
| Glu (E) | 6 | 3.8% |  |  |  |  |  |  |  |  |  |  |  |  |  |  |  |  |  |  |  |  |  |  |  |  |  |  |  |  |  |  |  |  |  |  |  |  |  |  |  |  |  |  |  |  |  |  |  |  |  |  |  |  |  |  |  |  |  |  |  |  |  |  |  |  |  |
| Gly (G) | 6 | 3.8% |  |  |  |  |  |  |  |  |  |  |  |  |  |  |  |  |  |  |  |  |  |  |  |  |  |  |  |  |  |  |  |  |  |  |  |  |  |  |  |  |  |  |  |  |  |  |  |  |  |  |  |  |  |  |  |  |  |  |  |  |  |  |  |  |  |
| His (H) | 10 | 6.2% |  |  |  |  |  |  |  |  |  |  |  |  |  |  |  |  |  |  |  |  |  |  |  |  |  |  |  |  |  |  |  |  |  |  |  |  |  |  |  |  |  |  |  |  |  |  |  |  |  |  |  |  |  |  |  |  |  |  |  |  |  |  |  |  |  |
| Ile (I) | 14 | 8.8% |  |  |  |  |  |  |  |  |  |  |  |  |  |  |  |  |  |  |  |  |  |  |  |  |  |  |  |  |  |  |  |  |  |  |  |  |  |  |  |  |  |  |  |  |  |  |  |  |  |  |  |  |  |  |  |  |  |  |  |  |  |  |  |  |  |
| Leu (L) | 12 | 7.5% |  |  |  |  |  |  |  |  |  |  |  |  |  |  |  |  |  |  |  |  |  |  |  |  |  |  |  |  |  |  |  |  |  |  |  |  |  |  |  |  |  |  |  |  |  |  |  |  |  |  |  |  |  |  |  |  |  |  |  |  |  |  |  |  |  |
| Lys (K) | 12 | 7.5% |  |  |  |  |  |  |  |  |  |  |  |  |  |  |  |  |  |  |  |  |  |  |  |  |  |  |  |  |  |  |  |  |  |  |  |  |  |  |  |  |  |  |  |  |  |  |  |  |  |  |  |  |  |  |  |  |  |  |  |  |  |  |  |  |  |
| Met (M) | 7 | 4.4% |  |  |  |  |  |  |  |  |  |  |  |  |  |  |  |  |  |  |  |  |  |  |  |  |  |  |  |  |  |  |  |  |  |  |  |  |  |  |  |  |  |  |  |  |  |  |  |  |  |  |  |  |  |  |  |  |  |  |  |  |  |  |  |  |  |
| Phe (F) | 9 | 5.6% |  |  |  |  |  |  |  |  |  |  |  |  |  |  |  |  |  |  |  |  |  |  |  |  |  |  |  |  |  |  |  |  |  |  |  |  |  |  |  |  |  |  |  |  |  |  |  |  |  |  |  |  |  |  |  |  |  |  |  |  |  |  |  |  |  |
| Pro (P) | 8 | 5.0% |  |  |  |  |  |  |  |  |  |  |  |  |  |  |  |  |  |  |  |  |  |  |  |  |  |  |  |  |  |  |  |  |  |  |  |  |  |  |  |  |  |  |  |  |  |  |  |  |  |  |  |  |  |  |  |  |  |  |  |  |  |  |  |  |  |
| Ser (S) | 11 | 6.9% |  |  |  |  |  |  |  |  |  |  |  |  |  |  |  |  |  |  |  |  |  |  |  |  |  |  |  |  |  |  |  |  |  |  |  |  |  |  |  |  |  |  |  |  |  |  |  |  |  |  |  |  |  |  |  |  |  |  |  |  |  |  |  |  |  |
| Thr (T) | 9 | 5.6% |  |  |  |  |  |  |  |  |  |  |  |  |  |  |  |  |  |  |  |  |  |  |  |  |  |  |  |  |  |  |  |  |  |  |  |  |  |  |  |  |  |  |  |  |  |  |  |  |  |  |  |  |  |  |  |  |  |  |  |  |  |  |  |  |  |
| Trp (W) | 4 | 2.5% |  |  |  |  |  |  |  |  |  |  |  |  |  |  |  |  |  |  |  |  |  |  |  |  |  |  |  |  |  |  |  |  |  |  |  |  |  |  |  |  |  |  |  |  |  |  |  |  |  |  |  |  |  |  |  |  |  |  |  |  |  |  |  |  |  |
| Tyr (Y) | 4 | 2.5% |  |  |  |  |  |  |  |  |  |  |  |  |  |  |  |  |  |  |  |  |  |  |  |  |  |  |  |  |  |  |  |  |  |  |  |  |  |  |  |  |  |  |  |  |  |  |  |  |  |  |  |  |  |  |  |  |  |  |  |  |  |  |  |  |  |
| Val (V) | 9 | 5.6% |  |  |  |  |  |  |  |  |  |  |  |  |  |  |  |  |  |  |  |  |  |  |  |  |  |  |  |  |  |  |  |  |  |  |  |  |  |  |  |  |  |  |  |  |  |  |  |  |  |  |  |  |  |  |  |  |  |  |  |  |  |  |  |  |  |
| Pyl (O) | 0 | 0.0% |  |  |  |  |  |  |  |  |  |  |  |  |  |  |  |  |  |  |  |  |  |  |  |  |  |  |  |  |  |  |  |  |  |  |  |  |  |  |  |  |  |  |  |  |  |  |  |  |  |  |  |  |  |  |  |  |  |  |  |  |  |  |  |  |  |
| Sec (U) | 0 | 0.0% |  |  |  |  |  |  |  |  |  |  |  |  |  |  |  |  |  |  |  |  |  |  |  |  |  |  |  |  |  |  |  |  |  |  |  |  |  |  |  |  |  |  |  |  |  |  |  |  |  |  |  |  |  |  |  |  |  |  |  |  |  |  |  |  |  |
| uterocalin P19 precursor [Sus scrofa]<br>NCBI Reference Sequence: NP_001431516.1<br>msllllavgl tllsstqa<br>rh wgpqdpnfne tlvggnwfsa<br>amasnqpqlm kaagarrili<br>hhiqvtpral rfhlhqrinq vcvptvmtan ktkkkf qyll<br>eyigqnr vfl ekvdpksyai<br>icthhkargr emvvvtllsr tpevsrdtIrl mflsycrkhg<br>lhpsvidlt rtdrcIharq | Number of amino acids: 162<br>Theoretical pI: 10.33<br>Molecular weight: 18721.79<br>Amino acid composition:<br>CSV format<br>Ala (A)<br>11<br>6.8%<br>Arg (R)<br>16<br>9.9%<br>Asn (N)<br>7<br>4.3%<br>Asp (D)<br>5<br>3.1%<br>Cys (C)<br>4<br>2.5%<br>Gln (Q)<br>8<br>4.9%<br>Glu (E)<br>6<br>3.7%<br>Gly (G)<br>8<br>4.9%<br>His (H)<br>10<br>6.2%<br>Ile (I)<br>8<br>4.9%<br>Leu (L)<br>15<br>9.3%<br>Lys (K)<br>9<br>5.6%<br>Met (M)<br>5<br>3.1%<br>Phe (F)<br>6<br>3.7%<br>Pro (P)<br>8<br>4.9%<br>Ser (S)<br>7<br>4.3%<br>Thr (T)<br>11<br>6.8%<br>Trp (W)<br>2<br>1.2%<br>Tyr (Y)<br>4<br>2.5%<br>Val (V)<br>12<br>7.4%<br>Pyl (O)<br>0<br>0.0%<br>Sec (U)<br>0<br>0.0% | Ala (A) | 11 | 6.8% | Arg (R) | 16 | 9.9% | Asn (N) | 7 | 4.3% | Asp (D) | 5 | 3.1% | Cys (C) | 4 | 2.5% | Gln (Q) | 8 | 4.9% | Glu (E) | 6 | 3.7% | Gly (G) | 8 | 4.9% | His (H) | 10 | 6.2% | Ile (I) | 8 | 4.9% | Leu (L) | 15 | 9.3% | Lys (K) | 9 | 5.6% | Met (M) | 5 | 3.1% | Phe (F) | 6 | 3.7% | Pro (P) | 8 | 4.9% | Ser (S) | 7 | 4.3% | Thr (T) | 11 | 6.8% | Trp (W) | 2 | 1.2% | Tyr (Y) | 4 | 2.5% | Val (V) | 12 | 7.4% | Pyl (O) | 0 | 0.0% | Sec (U) | 0 | 0.0% |
| Ala (A) | 11 | 6.8% |  |  |  |  |  |  |  |  |  |  |  |  |  |  |  |  |  |  |  |  |  |  |  |  |  |  |  |  |  |  |  |  |  |  |  |  |  |  |  |  |  |  |  |  |  |  |  |  |  |  |  |  |  |  |  |  |  |  |  |  |  |  |  |  |  |
| Arg (R) | 16 | 9.9% |  |  |  |  |  |  |  |  |  |  |  |  |  |  |  |  |  |  |  |  |  |  |  |  |  |  |  |  |  |  |  |  |  |  |  |  |  |  |  |  |  |  |  |  |  |  |  |  |  |  |  |  |  |  |  |  |  |  |  |  |  |  |  |  |  |
| Asn (N) | 7 | 4.3% |  |  |  |  |  |  |  |  |  |  |  |  |  |  |  |  |  |  |  |  |  |  |  |  |  |  |  |  |  |  |  |  |  |  |  |  |  |  |  |  |  |  |  |  |  |  |  |  |  |  |  |  |  |  |  |  |  |  |  |  |  |  |  |  |  |
| Asp (D) | 5 | 3.1% |  |  |  |  |  |  |  |  |  |  |  |  |  |  |  |  |  |  |  |  |  |  |  |  |  |  |  |  |  |  |  |  |  |  |  |  |  |  |  |  |  |  |  |  |  |  |  |  |  |  |  |  |  |  |  |  |  |  |  |  |  |  |  |  |  |
| Cys (C) | 4 | 2.5% |  |  |  |  |  |  |  |  |  |  |  |  |  |  |  |  |  |  |  |  |  |  |  |  |  |  |  |  |  |  |  |  |  |  |  |  |  |  |  |  |  |  |  |  |  |  |  |  |  |  |  |  |  |  |  |  |  |  |  |  |  |  |  |  |  |
| Gln (Q) | 8 | 4.9% |  |  |  |  |  |  |  |  |  |  |  |  |  |  |  |  |  |  |  |  |  |  |  |  |  |  |  |  |  |  |  |  |  |  |  |  |  |  |  |  |  |  |  |  |  |  |  |  |  |  |  |  |  |  |  |  |  |  |  |  |  |  |  |  |  |
| Glu (E) | 6 | 3.7% |  |  |  |  |  |  |  |  |  |  |  |  |  |  |  |  |  |  |  |  |  |  |  |  |  |  |  |  |  |  |  |  |  |  |  |  |  |  |  |  |  |  |  |  |  |  |  |  |  |  |  |  |  |  |  |  |  |  |  |  |  |  |  |  |  |
| Gly (G) | 8 | 4.9% |  |  |  |  |  |  |  |  |  |  |  |  |  |  |  |  |  |  |  |  |  |  |  |  |  |  |  |  |  |  |  |  |  |  |  |  |  |  |  |  |  |  |  |  |  |  |  |  |  |  |  |  |  |  |  |  |  |  |  |  |  |  |  |  |  |
| His (H) | 10 | 6.2% |  |  |  |  |  |  |  |  |  |  |  |  |  |  |  |  |  |  |  |  |  |  |  |  |  |  |  |  |  |  |  |  |  |  |  |  |  |  |  |  |  |  |  |  |  |  |  |  |  |  |  |  |  |  |  |  |  |  |  |  |  |  |  |  |  |
| Ile (I) | 8 | 4.9% |  |  |  |  |  |  |  |  |  |  |  |  |  |  |  |  |  |  |  |  |  |  |  |  |  |  |  |  |  |  |  |  |  |  |  |  |  |  |  |  |  |  |  |  |  |  |  |  |  |  |  |  |  |  |  |  |  |  |  |  |  |  |  |  |  |
| Leu (L) | 15 | 9.3% |  |  |  |  |  |  |  |  |  |  |  |  |  |  |  |  |  |  |  |  |  |  |  |  |  |  |  |  |  |  |  |  |  |  |  |  |  |  |  |  |  |  |  |  |  |  |  |  |  |  |  |  |  |  |  |  |  |  |  |  |  |  |  |  |  |
| Lys (K) | 9 | 5.6% |  |  |  |  |  |  |  |  |  |  |  |  |  |  |  |  |  |  |  |  |  |  |  |  |  |  |  |  |  |  |  |  |  |  |  |  |  |  |  |  |  |  |  |  |  |  |  |  |  |  |  |  |  |  |  |  |  |  |  |  |  |  |  |  |  |
| Met (M) | 5 | 3.1% |  |  |  |  |  |  |  |  |  |  |  |  |  |  |  |  |  |  |  |  |  |  |  |  |  |  |  |  |  |  |  |  |  |  |  |  |  |  |  |  |  |  |  |  |  |  |  |  |  |  |  |  |  |  |  |  |  |  |  |  |  |  |  |  |  |
| Phe (F) | 6 | 3.7% |  |  |  |  |  |  |  |  |  |  |  |  |  |  |  |  |  |  |  |  |  |  |  |  |  |  |  |  |  |  |  |  |  |  |  |  |  |  |  |  |  |  |  |  |  |  |  |  |  |  |  |  |  |  |  |  |  |  |  |  |  |  |  |  |  |
| Pro (P) | 8 | 4.9% |  |  |  |  |  |  |  |  |  |  |  |  |  |  |  |  |  |  |  |  |  |  |  |  |  |  |  |  |  |  |  |  |  |  |  |  |  |  |  |  |  |  |  |  |  |  |  |  |  |  |  |  |  |  |  |  |  |  |  |  |  |  |  |  |  |
| Ser (S) | 7 | 4.3% |  |  |  |  |  |  |  |  |  |  |  |  |  |  |  |  |  |  |  |  |  |  |  |  |  |  |  |  |  |  |  |  |  |  |  |  |  |  |  |  |  |  |  |  |  |  |  |  |  |  |  |  |  |  |  |  |  |  |  |  |  |  |  |  |  |
| Thr (T) | 11 | 6.8% |  |  |  |  |  |  |  |  |  |  |  |  |  |  |  |  |  |  |  |  |  |  |  |  |  |  |  |  |  |  |  |  |  |  |  |  |  |  |  |  |  |  |  |  |  |  |  |  |  |  |  |  |  |  |  |  |  |  |  |  |  |  |  |  |  |
| Trp (W) | 2 | 1.2% |  |  |  |  |  |  |  |  |  |  |  |  |  |  |  |  |  |  |  |  |  |  |  |  |  |  |  |  |  |  |  |  |  |  |  |  |  |  |  |  |  |  |  |  |  |  |  |  |  |  |  |  |  |  |  |  |  |  |  |  |  |  |  |  |  |
| Tyr (Y) | 4 | 2.5% |  |  |  |  |  |  |  |  |  |  |  |  |  |  |  |  |  |  |  |  |  |  |  |  |  |  |  |  |  |  |  |  |  |  |  |  |  |  |  |  |  |  |  |  |  |  |  |  |  |  |  |  |  |  |  |  |  |  |  |  |  |  |  |  |  |
| Val (V) | 12 | 7.4% |  |  |  |  |  |  |  |  |  |  |  |  |  |  |  |  |  |  |  |  |  |  |  |  |  |  |  |  |  |  |  |  |  |  |  |  |  |  |  |  |  |  |  |  |  |  |  |  |  |  |  |  |  |  |  |  |  |  |  |  |  |  |  |  |  |
| Pyl (O) | 0 | 0.0% |  |  |  |  |  |  |  |  |  |  |  |  |  |  |  |  |  |  |  |  |  |  |  |  |  |  |  |  |  |  |  |  |  |  |  |  |  |  |  |  |  |  |  |  |  |  |  |  |  |  |  |  |  |  |  |  |  |  |  |  |  |  |  |  |  |
| Sec (U) | 0 | 0.0% |  |  |  |  |  |  |  |  |  |  |  |  |  |  |  |  |  |  |  |  |  |  |  |  |  |  |  |  |  |  |  |  |  |  |  |  |  |  |  |  |  |  |  |  |  |  |  |  |  |  |  |  |  |  |  |  |  |  |  |  |  |  |  |  |  |

**uterocalin-like [Phoca vitulina]**

NCBI Reference Sequence: XP\_032274977.1

**mnapwlvvl allggpha** lh tgprdpnide nmqvngdwfs
taqasdepkl lwkdshvmff
vrkihvtlkt iefhlyrriq gmcvpiimma ntkkkkfyt
lkyaghnmiif leevdpsrfl
ifcihnhwhg ketvvvnlls rtptascdvm qtfrnycksh
gispaniinl tqtdhrlra

This has an unusual insert at its position 33. The near-identical sequence from Halichoerus grypus used instead.

**Number of amino acids:** 161**Theoretical pI:** 9.28**Molecular weight:** 18813.81**Amino acid composition:**CSV format

|  |  |  |
| --- | --- | --- |
| Ala (A) | 7 | 4.3% |
| Arg (R) | 9 | 5.6% |
| Asn (N) | 10 | 6.2% |
| Asp (D) | 8 | 5.0% |
| Cys (C) | 4 | 2.5% |
| Gln (Q) | 6 | 3.7% |
| Glu (E) | 6 | 3.7% |
| Gly (G) | 6 | 3.7% |
| His (H) | 10 | 6.2% |
| Ile (I) | 12 | 7.5% |
| Leu (L) | 12 | 7.5% |
| Lys (K) | 11 | 6.8% |
| Met (M) | 7 | 4.3% |
| Phe (F) | 9 | 5.6% |
| Pro (P) | 7 | 4.3% |
| Ser (S) | 8 | 5.0% |
| Thr (T) | 12 | 7.5% |
| Trp (W) | 3 | 1.9% |
| Tyr (Y) | 4 | 2.5% |
| Val (V) | 10 | 6.2% |
| Pyl (O) | 0 | 0.0% |
| Sec (U) | 0 | 0.0% |

**uterocalin-like [Halichoerus grypus]**

NCBI Reference Sequence: XP\_077917433.1

**mnapwlvvl allgaprpa** h gprdpniden mvngdwfsta
qasdeptllw kdshvmffvr
kihvtlktie fhlyrriqgm cvpiimmank tkkkfqytlk
yaghnmiifle evdpsrflif
cihnhwhgke tvvnllsrt ptascdvmt frnyckshgi
spaniintlq tdhrlra

**Number of amino acids:** 158**Theoretical pI:** 9.14**Molecular weight:** 18444.34**Amino acid composition:**CSV format

|  |  |  |
| --- | --- | --- |
| Ala (A) | 7 | 4.4% |
| Arg (R) | 9 | 5.7% |
| Asn (N) | 10 | 6.3% |
| Asp (D) | 8 | 5.1% |
| Cys (C) | 4 | 2.5% |
| Gln (Q) | 5 | 3.2% |
| Glu (E) | 6 | 3.8% |
| Gly (G) | 6 | 3.8% |
| His (H) | 10 | 6.3% |
| Ile (I) | 12 | 7.6% |
| Leu (L) | 11 | 7.0% |
| Lys (K) | 10 | 6.3% |
| Met (M) | 7 | 4.4% |
| Phe (F) | 9 | 5.7% |
| Pro (P) | 7 | 4.4% |
| Ser (S) | 8 | 5.1% |
| Thr (T) | 12 | 7.6% |
| Trp (W) | 3 | 1.9% |
| Tyr (Y) | 4 | 2.5% |
| Val (V) | 10 | 6.3% |
| Pyl (O) | 0 | 0.0% |
| Sec (U) | 0 | 0.0% |

**uterocalin-like isoform X2 [Leopardus geoffroyi]**

**Geoffroy's cat**

NCBI Reference Sequence: XP\_045323916.1

**mnvrlalvl tllggpqa** lh rgpqdpnfte nlvsgnwfsi
arasnepkll rmdsdtmffi
hkihvtfktm qfhvhrrvkg rcipivltas ktrrksqyvl
kyaghstifm ekadpqrivi
fcihnrhrk kavmvdllgr tptagwdilh tfksycqsrq
tgptdiisl qtgrclharp

**Number of amino acids:** 162

**Theoretical pI:** 10.64

**Molecular weight:** 18759.83

**Amino acid composition:** CSV format

|  |  |  |
| --- | --- | --- |
| Ala (A) | 8 | 4.9% |
| Arg (R) | 16 | 9.9% |
| Asn (N) | 5 | 3.1% |
| Asp (D) | 7 | 4.3% |
| Cys (C) | 4 | 2.5% |
| Gln (Q) | 6 | 3.7% |
| Glu (E) | 3 | 1.9% |
| Gly (G) | 9 | 5.6% |
| His (H) | 10 | 6.2% |
| Ile (I) | 11 | 6.8% |
| Leu (L) | 11 | 6.8% |
| Lys (K) | 11 | 6.8% |
| Met (M) | 5 | 3.1% |
| Phe (F) | 10 | 6.2% |
| Pro (P) | 8 | 4.9% |
| Ser (S) | 10 | 6.2% |
| Thr (T) | 14 | 8.6% |
| Trp (W) | 2 | 1.2% |
| Tyr (Y) | 3 | 1.9% |
| Val (V) | 9 | 5.6% |
| Pyl (O) | 0 | 0.0% |
| Sec (U) | 0 | 0.0% |

**uterocalin-like [Callorhinus ursinus] northern fur seal**

NCBI Reference Sequence: XP\_073746189.1

**mntprlvvl allrgpha** lh tpggdpside nmvngdwfsi
aqassepkll wkdshmmffv
rkihvsikti efhlyrriqg mcvpiimman ktkkkfqymf
kyaghsvifp eevapsrfif
cihnhwhgke tvvnllsrt ppaswdvmqt fknyckshgi
spaniinl trdcclha

**Number of amino acids:** 159

**Theoretical pI:** 9.17

**Molecular weight:** 18332.41

**Amino acid composition:** CSV format

|  |  |  |
| --- | --- | --- |
| Ala (A) | 8 | 5.0% |
| Arg (R) | 6 | 3.8% |
| Asn (N) | 8 | 5.0% |
| Asp (D) | 6 | 3.8% |
| Cys (C) | 5 | 3.1% |
| Gln (Q) | 4 | 2.5% |
| Glu (E) | 6 | 3.8% |
| Gly (G) | 7 | 4.4% |
| His (H) | 10 | 6.3% |
| Ile (I) | 13 | 8.2% |
| Leu (L) | 10 | 6.3% |
| Lys (K) | 12 | 7.5% |
| Met (M) | 8 | 5.0% |
| Phe (F) | 9 | 5.7% |
| Pro (P) | 9 | 5.7% |
| Ser (S) | 12 | 7.5% |
| Thr (T) | 8 | 5.0% |
| Trp (W) | 4 | 2.5% |
| Tyr (Y) | 4 | 2.5% |
| Val (V) | 10 | 6.3% |
| Pyl (O) | 0 | 0.0% |
| Sec (U) | 0 | 0.0% |

|  |  |  |  |  |  |  |  |  |  |  |  |  |  |  |  |  |  |  |  |  |  |  |  |  |  |  |  |  |  |  |  |  |  |  |  |  |  |  |  |  |  |  |  |  |  |  |  |  |  |  |  |  |  |  |  |  |  |  |  |  |  |  |  |  |  |  |  |
| --- | --- | --- | --- | --- | --- | --- | --- | --- | --- | --- | --- | --- | --- | --- | --- | --- | --- | --- | --- | --- | --- | --- | --- | --- | --- | --- | --- | --- | --- | --- | --- | --- | --- | --- | --- | --- | --- | --- | --- | --- | --- | --- | --- | --- | --- | --- | --- | --- | --- | --- | --- | --- | --- | --- | --- | --- | --- | --- | --- | --- | --- | --- | --- | --- | --- | --- | --- |
| **uterocalin-like isoform 1-T1 [Megaptera novaeangliae] humpback whale**<br>GenBank: KAM9082391.1<br>mnllllavgl tllgcpqa lh wgpqdpnfne tlvsgewfla<br>glasnqpkll rededarlli<br>hriqvtpral qlhlrrkvng acvpitmtan ktkrkfyll<br>enaaqnravl gkdpksyii<br>lcnhreerkk evvavslsr tpeaspdall mftdycrnhg<br>ihttniinvrt rtggcpplpq<br>prgaegtart strlatdrsl ra | **Number of amino acids:** 162<br>**Theoretical pI:** 9.51<br>**Molecular weight:** 18375.20<br>**Amino acid composition:** CSV format<br>Ala (A)<br>12<br>7.4%<br>Arg (R)<br>13<br>8.0%<br>Asn (N)<br>11<br>6.8%<br>Asp (D)<br>6<br>3.7%<br>Cys (C)<br>4<br>2.5%<br>Gln (Q)<br>7<br>4.3%<br>Glu (E)<br>9<br>5.6%<br>Gly (G)<br>8<br>4.9%<br>His (H)<br>6<br>3.7%<br>Ile (I)<br>8<br>4.9%<br>Leu (L)<br>20<br>12.3%<br>Lys (K)<br>9<br>5.6%<br>Met (M)<br>2<br>1.2%<br>Phe (F)<br>5<br>3.1%<br>Pro (P)<br>11<br>6.8%<br>Ser (S)<br>6<br>3.7%<br>Thr (T)<br>11<br>6.8%<br>Trp (W)<br>2<br>1.2%<br>Tyr (Y)<br>3<br>1.9%<br>Val (V)<br>9<br>5.6%<br>Pyl (O)<br>0<br>0.0%<br>Sec (U)<br>0<br>0.0% | Ala (A) | 12 | 7.4% | Arg (R) | 13 | 8.0% | Asn (N) | 11 | 6.8% | Asp (D) | 6 | 3.7% | Cys (C) | 4 | 2.5% | Gln (Q) | 7 | 4.3% | Glu (E) | 9 | 5.6% | Gly (G) | 8 | 4.9% | His (H) | 6 | 3.7% | Ile (I) | 8 | 4.9% | Leu (L) | 20 | 12.3% | Lys (K) | 9 | 5.6% | Met (M) | 2 | 1.2% | Phe (F) | 5 | 3.1% | Pro (P) | 11 | 6.8% | Ser (S) | 6 | 3.7% | Thr (T) | 11 | 6.8% | Trp (W) | 2 | 1.2% | Tyr (Y) | 3 | 1.9% | Val (V) | 9 | 5.6% | Pyl (O) | 0 | 0.0% | Sec (U) | 0 | 0.0% |
| Ala (A) | 12 | 7.4% |  |  |  |  |  |  |  |  |  |  |  |  |  |  |  |  |  |  |  |  |  |  |  |  |  |  |  |  |  |  |  |  |  |  |  |  |  |  |  |  |  |  |  |  |  |  |  |  |  |  |  |  |  |  |  |  |  |  |  |  |  |  |  |  |  |
| Arg (R) | 13 | 8.0% |  |  |  |  |  |  |  |  |  |  |  |  |  |  |  |  |  |  |  |  |  |  |  |  |  |  |  |  |  |  |  |  |  |  |  |  |  |  |  |  |  |  |  |  |  |  |  |  |  |  |  |  |  |  |  |  |  |  |  |  |  |  |  |  |  |
| Asn (N) | 11 | 6.8% |  |  |  |  |  |  |  |  |  |  |  |  |  |  |  |  |  |  |  |  |  |  |  |  |  |  |  |  |  |  |  |  |  |  |  |  |  |  |  |  |  |  |  |  |  |  |  |  |  |  |  |  |  |  |  |  |  |  |  |  |  |  |  |  |  |
| Asp (D) | 6 | 3.7% |  |  |  |  |  |  |  |  |  |  |  |  |  |  |  |  |  |  |  |  |  |  |  |  |  |  |  |  |  |  |  |  |  |  |  |  |  |  |  |  |  |  |  |  |  |  |  |  |  |  |  |  |  |  |  |  |  |  |  |  |  |  |  |  |  |
| Cys (C) | 4 | 2.5% |  |  |  |  |  |  |  |  |  |  |  |  |  |  |  |  |  |  |  |  |  |  |  |  |  |  |  |  |  |  |  |  |  |  |  |  |  |  |  |  |  |  |  |  |  |  |  |  |  |  |  |  |  |  |  |  |  |  |  |  |  |  |  |  |  |
| Gln (Q) | 7 | 4.3% |  |  |  |  |  |  |  |  |  |  |  |  |  |  |  |  |  |  |  |  |  |  |  |  |  |  |  |  |  |  |  |  |  |  |  |  |  |  |  |  |  |  |  |  |  |  |  |  |  |  |  |  |  |  |  |  |  |  |  |  |  |  |  |  |  |
| Glu (E) | 9 | 5.6% |  |  |  |  |  |  |  |  |  |  |  |  |  |  |  |  |  |  |  |  |  |  |  |  |  |  |  |  |  |  |  |  |  |  |  |  |  |  |  |  |  |  |  |  |  |  |  |  |  |  |  |  |  |  |  |  |  |  |  |  |  |  |  |  |  |
| Gly (G) | 8 | 4.9% |  |  |  |  |  |  |  |  |  |  |  |  |  |  |  |  |  |  |  |  |  |  |  |  |  |  |  |  |  |  |  |  |  |  |  |  |  |  |  |  |  |  |  |  |  |  |  |  |  |  |  |  |  |  |  |  |  |  |  |  |  |  |  |  |  |
| His (H) | 6 | 3.7% |  |  |  |  |  |  |  |  |  |  |  |  |  |  |  |  |  |  |  |  |  |  |  |  |  |  |  |  |  |  |  |  |  |  |  |  |  |  |  |  |  |  |  |  |  |  |  |  |  |  |  |  |  |  |  |  |  |  |  |  |  |  |  |  |  |
| Ile (I) | 8 | 4.9% |  |  |  |  |  |  |  |  |  |  |  |  |  |  |  |  |  |  |  |  |  |  |  |  |  |  |  |  |  |  |  |  |  |  |  |  |  |  |  |  |  |  |  |  |  |  |  |  |  |  |  |  |  |  |  |  |  |  |  |  |  |  |  |  |  |
| Leu (L) | 20 | 12.3% |  |  |  |  |  |  |  |  |  |  |  |  |  |  |  |  |  |  |  |  |  |  |  |  |  |  |  |  |  |  |  |  |  |  |  |  |  |  |  |  |  |  |  |  |  |  |  |  |  |  |  |  |  |  |  |  |  |  |  |  |  |  |  |  |  |
| Lys (K) | 9 | 5.6% |  |  |  |  |  |  |  |  |  |  |  |  |  |  |  |  |  |  |  |  |  |  |  |  |  |  |  |  |  |  |  |  |  |  |  |  |  |  |  |  |  |  |  |  |  |  |  |  |  |  |  |  |  |  |  |  |  |  |  |  |  |  |  |  |  |
| Met (M) | 2 | 1.2% |  |  |  |  |  |  |  |  |  |  |  |  |  |  |  |  |  |  |  |  |  |  |  |  |  |  |  |  |  |  |  |  |  |  |  |  |  |  |  |  |  |  |  |  |  |  |  |  |  |  |  |  |  |  |  |  |  |  |  |  |  |  |  |  |  |
| Phe (F) | 5 | 3.1% |  |  |  |  |  |  |  |  |  |  |  |  |  |  |  |  |  |  |  |  |  |  |  |  |  |  |  |  |  |  |  |  |  |  |  |  |  |  |  |  |  |  |  |  |  |  |  |  |  |  |  |  |  |  |  |  |  |  |  |  |  |  |  |  |  |
| Pro (P) | 11 | 6.8% |  |  |  |  |  |  |  |  |  |  |  |  |  |  |  |  |  |  |  |  |  |  |  |  |  |  |  |  |  |  |  |  |  |  |  |  |  |  |  |  |  |  |  |  |  |  |  |  |  |  |  |  |  |  |  |  |  |  |  |  |  |  |  |  |  |
| Ser (S) | 6 | 3.7% |  |  |  |  |  |  |  |  |  |  |  |  |  |  |  |  |  |  |  |  |  |  |  |  |  |  |  |  |  |  |  |  |  |  |  |  |  |  |  |  |  |  |  |  |  |  |  |  |  |  |  |  |  |  |  |  |  |  |  |  |  |  |  |  |  |
| Thr (T) | 11 | 6.8% |  |  |  |  |  |  |  |  |  |  |  |  |  |  |  |  |  |  |  |  |  |  |  |  |  |  |  |  |  |  |  |  |  |  |  |  |  |  |  |  |  |  |  |  |  |  |  |  |  |  |  |  |  |  |  |  |  |  |  |  |  |  |  |  |  |
| Trp (W) | 2 | 1.2% |  |  |  |  |  |  |  |  |  |  |  |  |  |  |  |  |  |  |  |  |  |  |  |  |  |  |  |  |  |  |  |  |  |  |  |  |  |  |  |  |  |  |  |  |  |  |  |  |  |  |  |  |  |  |  |  |  |  |  |  |  |  |  |  |  |
| Tyr (Y) | 3 | 1.9% |  |  |  |  |  |  |  |  |  |  |  |  |  |  |  |  |  |  |  |  |  |  |  |  |  |  |  |  |  |  |  |  |  |  |  |  |  |  |  |  |  |  |  |  |  |  |  |  |  |  |  |  |  |  |  |  |  |  |  |  |  |  |  |  |  |
| Val (V) | 9 | 5.6% |  |  |  |  |  |  |  |  |  |  |  |  |  |  |  |  |  |  |  |  |  |  |  |  |  |  |  |  |  |  |  |  |  |  |  |  |  |  |  |  |  |  |  |  |  |  |  |  |  |  |  |  |  |  |  |  |  |  |  |  |  |  |  |  |  |
| Pyl (O) | 0 | 0.0% |  |  |  |  |  |  |  |  |  |  |  |  |  |  |  |  |  |  |  |  |  |  |  |  |  |  |  |  |  |  |  |  |  |  |  |  |  |  |  |  |  |  |  |  |  |  |  |  |  |  |  |  |  |  |  |  |  |  |  |  |  |  |  |  |  |
| Sec (U) | 0 | 0.0% |  |  |  |  |  |  |  |  |  |  |  |  |  |  |  |  |  |  |  |  |  |  |  |  |  |  |  |  |  |  |  |  |  |  |  |  |  |  |  |  |  |  |  |  |  |  |  |  |  |  |  |  |  |  |  |  |  |  |  |  |  |  |  |  |  |
| **hypothetical protein J1605_017252 [Eschrichtius robustus] grey whale**<br>GenBank: KAJ8797520.1<br>mnllllavgl tllgcpqa lh wgpqdpnfne tlvsgewfla<br>glasnqpkll kedaralli<br>hriqvtpral qlhlrrkvng acvpitmtan ktkrkfyll<br>enaaqnravl gkdpksyii<br>lcnhreerek evvavslsr tpeaspdall mftdycrnhg<br>ihttniinvrt rtgrlraqvg<br>dl | **Number of amino acids:** 162<br>**Theoretical pI:** 9.62<br>**Molecular weight:** 18436.22<br>**Amino acid composition:** CSV format<br>Ala (A)<br>13<br>8.0%<br>Arg (R)<br>14<br>8.6%<br>Asn (N)<br>11<br>6.8%<br>Asp (D)<br>6<br>3.7%<br>Cys (C)<br>3<br>1.9%<br>Gln (Q)<br>7<br>4.3%<br>Glu (E)<br>10<br>6.2%<br>Gly (G)<br>8<br>4.9%<br>His (H)<br>6<br>3.7%<br>Ile (I)<br>8<br>4.9%<br>Leu (L)<br>20<br>12.3%<br>Lys (K)<br>9<br>5.6%<br>Met (M)<br>2<br>1.2%<br>Phe (F)<br>5<br>3.1%<br>Pro (P)<br>8<br>4.9%<br>Ser (S)<br>6<br>3.7%<br>Thr (T)<br>11<br>6.8%<br>Trp (W)<br>2<br>1.2%<br>Tyr (Y)<br>3<br>1.9%<br>Val (V)<br>10<br>6.2%<br>Pyl (O)<br>0<br>0.0%<br>Sec (U)<br>0<br>0.0% | Ala (A) | 13 | 8.0% | Arg (R) | 14 | 8.6% | Asn (N) | 11 | 6.8% | Asp (D) | 6 | 3.7% | Cys (C) | 3 | 1.9% | Gln (Q) | 7 | 4.3% | Glu (E) | 10 | 6.2% | Gly (G) | 8 | 4.9% | His (H) | 6 | 3.7% | Ile (I) | 8 | 4.9% | Leu (L) | 20 | 12.3% | Lys (K) | 9 | 5.6% | Met (M) | 2 | 1.2% | Phe (F) | 5 | 3.1% | Pro (P) | 8 | 4.9% | Ser (S) | 6 | 3.7% | Thr (T) | 11 | 6.8% | Trp (W) | 2 | 1.2% | Tyr (Y) | 3 | 1.9% | Val (V) | 10 | 6.2% | Pyl (O) | 0 | 0.0% | Sec (U) | 0 | 0.0% |
| Ala (A) | 13 | 8.0% |  |  |  |  |  |  |  |  |  |  |  |  |  |  |  |  |  |  |  |  |  |  |  |  |  |  |  |  |  |  |  |  |  |  |  |  |  |  |  |  |  |  |  |  |  |  |  |  |  |  |  |  |  |  |  |  |  |  |  |  |  |  |  |  |  |
| Arg (R) | 14 | 8.6% |  |  |  |  |  |  |  |  |  |  |  |  |  |  |  |  |  |  |  |  |  |  |  |  |  |  |  |  |  |  |  |  |  |  |  |  |  |  |  |  |  |  |  |  |  |  |  |  |  |  |  |  |  |  |  |  |  |  |  |  |  |  |  |  |  |
| Asn (N) | 11 | 6.8% |  |  |  |  |  |  |  |  |  |  |  |  |  |  |  |  |  |  |  |  |  |  |  |  |  |  |  |  |  |  |  |  |  |  |  |  |  |  |  |  |  |  |  |  |  |  |  |  |  |  |  |  |  |  |  |  |  |  |  |  |  |  |  |  |  |
| Asp (D) | 6 | 3.7% |  |  |  |  |  |  |  |  |  |  |  |  |  |  |  |  |  |  |  |  |  |  |  |  |  |  |  |  |  |  |  |  |  |  |  |  |  |  |  |  |  |  |  |  |  |  |  |  |  |  |  |  |  |  |  |  |  |  |  |  |  |  |  |  |  |
| Cys (C) | 3 | 1.9% |  |  |  |  |  |  |  |  |  |  |  |  |  |  |  |  |  |  |  |  |  |  |  |  |  |  |  |  |  |  |  |  |  |  |  |  |  |  |  |  |  |  |  |  |  |  |  |  |  |  |  |  |  |  |  |  |  |  |  |  |  |  |  |  |  |
| Gln (Q) | 7 | 4.3% |  |  |  |  |  |  |  |  |  |  |  |  |  |  |  |  |  |  |  |  |  |  |  |  |  |  |  |  |  |  |  |  |  |  |  |  |  |  |  |  |  |  |  |  |  |  |  |  |  |  |  |  |  |  |  |  |  |  |  |  |  |  |  |  |  |
| Glu (E) | 10 | 6.2% |  |  |  |  |  |  |  |  |  |  |  |  |  |  |  |  |  |  |  |  |  |  |  |  |  |  |  |  |  |  |  |  |  |  |  |  |  |  |  |  |  |  |  |  |  |  |  |  |  |  |  |  |  |  |  |  |  |  |  |  |  |  |  |  |  |
| Gly (G) | 8 | 4.9% |  |  |  |  |  |  |  |  |  |  |  |  |  |  |  |  |  |  |  |  |  |  |  |  |  |  |  |  |  |  |  |  |  |  |  |  |  |  |  |  |  |  |  |  |  |  |  |  |  |  |  |  |  |  |  |  |  |  |  |  |  |  |  |  |  |
| His (H) | 6 | 3.7% |  |  |  |  |  |  |  |  |  |  |  |  |  |  |  |  |  |  |  |  |  |  |  |  |  |  |  |  |  |  |  |  |  |  |  |  |  |  |  |  |  |  |  |  |  |  |  |  |  |  |  |  |  |  |  |  |  |  |  |  |  |  |  |  |  |
| Ile (I) | 8 | 4.9% |  |  |  |  |  |  |  |  |  |  |  |  |  |  |  |  |  |  |  |  |  |  |  |  |  |  |  |  |  |  |  |  |  |  |  |  |  |  |  |  |  |  |  |  |  |  |  |  |  |  |  |  |  |  |  |  |  |  |  |  |  |  |  |  |  |
| Leu (L) | 20 | 12.3% |  |  |  |  |  |  |  |  |  |  |  |  |  |  |  |  |  |  |  |  |  |  |  |  |  |  |  |  |  |  |  |  |  |  |  |  |  |  |  |  |  |  |  |  |  |  |  |  |  |  |  |  |  |  |  |  |  |  |  |  |  |  |  |  |  |
| Lys (K) | 9 | 5.6% |  |  |  |  |  |  |  |  |  |  |  |  |  |  |  |  |  |  |  |  |  |  |  |  |  |  |  |  |  |  |  |  |  |  |  |  |  |  |  |  |  |  |  |  |  |  |  |  |  |  |  |  |  |  |  |  |  |  |  |  |  |  |  |  |  |
| Met (M) | 2 | 1.2% |  |  |  |  |  |  |  |  |  |  |  |  |  |  |  |  |  |  |  |  |  |  |  |  |  |  |  |  |  |  |  |  |  |  |  |  |  |  |  |  |  |  |  |  |  |  |  |  |  |  |  |  |  |  |  |  |  |  |  |  |  |  |  |  |  |
| Phe (F) | 5 | 3.1% |  |  |  |  |  |  |  |  |  |  |  |  |  |  |  |  |  |  |  |  |  |  |  |  |  |  |  |  |  |  |  |  |  |  |  |  |  |  |  |  |  |  |  |  |  |  |  |  |  |  |  |  |  |  |  |  |  |  |  |  |  |  |  |  |  |
| Pro (P) | 8 | 4.9% |  |  |  |  |  |  |  |  |  |  |  |  |  |  |  |  |  |  |  |  |  |  |  |  |  |  |  |  |  |  |  |  |  |  |  |  |  |  |  |  |  |  |  |  |  |  |  |  |  |  |  |  |  |  |  |  |  |  |  |  |  |  |  |  |  |
| Ser (S) | 6 | 3.7% |  |  |  |  |  |  |  |  |  |  |  |  |  |  |  |  |  |  |  |  |  |  |  |  |  |  |  |  |  |  |  |  |  |  |  |  |  |  |  |  |  |  |  |  |  |  |  |  |  |  |  |  |  |  |  |  |  |  |  |  |  |  |  |  |  |
| Thr (T) | 11 | 6.8% |  |  |  |  |  |  |  |  |  |  |  |  |  |  |  |  |  |  |  |  |  |  |  |  |  |  |  |  |  |  |  |  |  |  |  |  |  |  |  |  |  |  |  |  |  |  |  |  |  |  |  |  |  |  |  |  |  |  |  |  |  |  |  |  |  |
| Trp (W) | 2 | 1.2% |  |  |  |  |  |  |  |  |  |  |  |  |  |  |  |  |  |  |  |  |  |  |  |  |  |  |  |  |  |  |  |  |  |  |  |  |  |  |  |  |  |  |  |  |  |  |  |  |  |  |  |  |  |  |  |  |  |  |  |  |  |  |  |  |  |
| Tyr (Y) | 3 | 1.9% |  |  |  |  |  |  |  |  |  |  |  |  |  |  |  |  |  |  |  |  |  |  |  |  |  |  |  |  |  |  |  |  |  |  |  |  |  |  |  |  |  |  |  |  |  |  |  |  |  |  |  |  |  |  |  |  |  |  |  |  |  |  |  |  |  |
| Val (V) | 10 | 6.2% |  |  |  |  |  |  |  |  |  |  |  |  |  |  |  |  |  |  |  |  |  |  |  |  |  |  |  |  |  |  |  |  |  |  |  |  |  |  |  |  |  |  |  |  |  |  |  |  |  |  |  |  |  |  |  |  |  |  |  |  |  |  |  |  |  |
| Pyl (O) | 0 | 0.0% |  |  |  |  |  |  |  |  |  |  |  |  |  |  |  |  |  |  |  |  |  |  |  |  |  |  |  |  |  |  |  |  |  |  |  |  |  |  |  |  |  |  |  |  |  |  |  |  |  |  |  |  |  |  |  |  |  |  |  |  |  |  |  |  |  |
| Sec (U) | 0 | 0.0% |  |  |  |  |  |  |  |  |  |  |  |  |  |  |  |  |  |  |  |  |  |  |  |  |  |  |  |  |  |  |  |  |  |  |  |  |  |  |  |  |  |  |  |  |  |  |  |  |  |  |  |  |  |  |  |  |  |  |  |  |  |  |  |  |  |

**uterocalin-like [Balaenoptera musculus] pygmy blue whale**

NCBI Reference Sequence: XP\_036712196.1

mnllllavgl tllgcpqa lh wgpqdpnfne tlvsgewfla
glasnqpkll rededarlli
hriqvtpal qlhlrrkvng acvpitmtan ktkrrfqyll
enaaqnravl ekvdpksyii
lcnhheerek evvavslsr tpeaspdall mftdycrngh
ihttniinvrt rgtvsprhqr

trrsrltqga lqlphplgva salleanpav snkrlprtqc
algglsprts lagasmvtrs
cpvtpphpr lltgdgdpp gdtklaldr prgleklgv
hrdavsakl gdtgrgrgq
aprlpwspqa qgcgq

**BLAST search of C-terminal extension finds –
uterocalin-like [Balaenoptera physalus]**

Sequence ID: XP\_082325636.1 – itself
hypothetical protein E2I00\_001780, partial
[Balaenoptera physalus]

Sequence ID: KAB0394076.1 – 99%
hypothetical protein EI555\_013315, partial
[Monodon monoceros]

Sequence ID: TKC51741.1 – 88% extensive
identity

protein FAM193B-like [Balaenoptera
acutorostrata]

Sequence ID: XP\_007194825.2 – 81% identity
with large blocks of identity with gaps

**Number of amino acids:** 162

**Theoretical pI:** 9.37

**Molecular weight:** 18653.37

**Amino acid composition:** CSV format

|  |  |  |
| --- | --- | --- |
| Ala (A) | 11 | 6.8% |
| Arg (R) | 15 | 9.3% |
| Asn (N) | 11 | 6.8% |
| Asp (D) | 6 | 3.7% |
| Cys (C) | 3 | 1.9% |
| Gln (Q) | 7 | 4.3% |
| Glu (E) | 11 | 6.8% |
| Gly (G) | 6 | 3.7% |
| His (H) | 8 | 4.9% |
| Ile (I) | 8 | 4.9% |
| Leu (L) | 19 | 11.7% |
| Lys (K) | 7 | 4.3% |
| Met (M) | 2 | 1.2% |
| Phe (F) | 5 | 3.1% |
| Pro (P) | 9 | 5.6% |
| Ser (S) | 7 | 4.3% |
| Thr (T) | 11 | 6.8% |
| Trp (W) | 2 | 1.2% |
| Tyr (Y) | 3 | 1.9% |
| Val (V) | 11 | 6.8% |
| Pyl (O) | 0 | 0.0% |
| Sec (U) | 0 | 0.0% |

|  |  |  |  |  |  |  |  |  |  |  |  |  |  |  |  |  |  |  |  |  |  |  |  |  |  |  |  |  |  |  |  |  |  |  |  |  |  |  |  |  |  |  |  |  |  |  |  |  |  |  |  |  |  |  |  |  |  |  |  |  |  |  |  |  |  |  |  |
| --- | --- | --- | --- | --- | --- | --- | --- | --- | --- | --- | --- | --- | --- | --- | --- | --- | --- | --- | --- | --- | --- | --- | --- | --- | --- | --- | --- | --- | --- | --- | --- | --- | --- | --- | --- | --- | --- | --- | --- | --- | --- | --- | --- | --- | --- | --- | --- | --- | --- | --- | --- | --- | --- | --- | --- | --- | --- | --- | --- | --- | --- | --- | --- | --- | --- | --- | --- |
| uterocalin-like [Balaenoptera ricei]<br>NCBI Reference Sequence: XP_059782608.1<br>mnllllavgl tllgcpqa lh wgpqdpnfne tlvsgewfla<br>glasnqpkll kededarlli<br>hrvqvtpral qlhlrrkvng acvpitmtan kmkrkfyll<br>enaaqnrvfl gkadpkysyii<br>lcnhreerek evvavslsr tpeaspdall mftdycrnhg<br>ihttniinvrt rtggcpplpq<br>prgaegtart sarlatdrc l ca | Number of amino acids: 162<br>Theoretical pI: 9.18<br>Molecular weight: 18364.19<br>Amino acid composition:<br>CSV format<br>Ala (A)<br>12<br>7.4%<br>Arg (R)<br>12<br>7.4%<br>Asn (N)<br>11<br>6.8%<br>Asp (D)<br>6<br>3.7%<br>Cys (C)<br>4<br>2.5%<br>Gln (Q)<br>7<br>4.3%<br>Glu (E)<br>10<br>6.2%<br>Gly (G)<br>8<br>4.9%<br>His (H)<br>6<br>3.7%<br>Ile (I)<br>7<br>4.3%<br>Leu (L)<br>20<br>12.3%<br>Lys (K)<br>9<br>5.6%<br>Met (M)<br>3<br>1.9%<br>Phe (F)<br>5<br>3.1%<br>Pro (P)<br>11<br>6.8%<br>Ser (S)<br>6<br>3.7%<br>Thr (T)<br>10<br>6.2%<br>Trp (W)<br>2<br>1.2%<br>Tyr (Y)<br>3<br>1.9%<br>Val (V)<br>10<br>6.2%<br>Pyl (O)<br>0<br>0.0%<br>Sec (U)<br>0<br>0.0% | Ala (A) | 12 | 7.4% | Arg (R) | 12 | 7.4% | Asn (N) | 11 | 6.8% | Asp (D) | 6 | 3.7% | Cys (C) | 4 | 2.5% | Gln (Q) | 7 | 4.3% | Glu (E) | 10 | 6.2% | Gly (G) | 8 | 4.9% | His (H) | 6 | 3.7% | Ile (I) | 7 | 4.3% | Leu (L) | 20 | 12.3% | Lys (K) | 9 | 5.6% | Met (M) | 3 | 1.9% | Phe (F) | 5 | 3.1% | Pro (P) | 11 | 6.8% | Ser (S) | 6 | 3.7% | Thr (T) | 10 | 6.2% | Trp (W) | 2 | 1.2% | Tyr (Y) | 3 | 1.9% | Val (V) | 10 | 6.2% | Pyl (O) | 0 | 0.0% | Sec (U) | 0 | 0.0% |
| Ala (A) | 12 | 7.4% |  |  |  |  |  |  |  |  |  |  |  |  |  |  |  |  |  |  |  |  |  |  |  |  |  |  |  |  |  |  |  |  |  |  |  |  |  |  |  |  |  |  |  |  |  |  |  |  |  |  |  |  |  |  |  |  |  |  |  |  |  |  |  |  |  |
| Arg (R) | 12 | 7.4% |  |  |  |  |  |  |  |  |  |  |  |  |  |  |  |  |  |  |  |  |  |  |  |  |  |  |  |  |  |  |  |  |  |  |  |  |  |  |  |  |  |  |  |  |  |  |  |  |  |  |  |  |  |  |  |  |  |  |  |  |  |  |  |  |  |
| Asn (N) | 11 | 6.8% |  |  |  |  |  |  |  |  |  |  |  |  |  |  |  |  |  |  |  |  |  |  |  |  |  |  |  |  |  |  |  |  |  |  |  |  |  |  |  |  |  |  |  |  |  |  |  |  |  |  |  |  |  |  |  |  |  |  |  |  |  |  |  |  |  |
| Asp (D) | 6 | 3.7% |  |  |  |  |  |  |  |  |  |  |  |  |  |  |  |  |  |  |  |  |  |  |  |  |  |  |  |  |  |  |  |  |  |  |  |  |  |  |  |  |  |  |  |  |  |  |  |  |  |  |  |  |  |  |  |  |  |  |  |  |  |  |  |  |  |
| Cys (C) | 4 | 2.5% |  |  |  |  |  |  |  |  |  |  |  |  |  |  |  |  |  |  |  |  |  |  |  |  |  |  |  |  |  |  |  |  |  |  |  |  |  |  |  |  |  |  |  |  |  |  |  |  |  |  |  |  |  |  |  |  |  |  |  |  |  |  |  |  |  |
| Gln (Q) | 7 | 4.3% |  |  |  |  |  |  |  |  |  |  |  |  |  |  |  |  |  |  |  |  |  |  |  |  |  |  |  |  |  |  |  |  |  |  |  |  |  |  |  |  |  |  |  |  |  |  |  |  |  |  |  |  |  |  |  |  |  |  |  |  |  |  |  |  |  |
| Glu (E) | 10 | 6.2% |  |  |  |  |  |  |  |  |  |  |  |  |  |  |  |  |  |  |  |  |  |  |  |  |  |  |  |  |  |  |  |  |  |  |  |  |  |  |  |  |  |  |  |  |  |  |  |  |  |  |  |  |  |  |  |  |  |  |  |  |  |  |  |  |  |
| Gly (G) | 8 | 4.9% |  |  |  |  |  |  |  |  |  |  |  |  |  |  |  |  |  |  |  |  |  |  |  |  |  |  |  |  |  |  |  |  |  |  |  |  |  |  |  |  |  |  |  |  |  |  |  |  |  |  |  |  |  |  |  |  |  |  |  |  |  |  |  |  |  |
| His (H) | 6 | 3.7% |  |  |  |  |  |  |  |  |  |  |  |  |  |  |  |  |  |  |  |  |  |  |  |  |  |  |  |  |  |  |  |  |  |  |  |  |  |  |  |  |  |  |  |  |  |  |  |  |  |  |  |  |  |  |  |  |  |  |  |  |  |  |  |  |  |
| Ile (I) | 7 | 4.3% |  |  |  |  |  |  |  |  |  |  |  |  |  |  |  |  |  |  |  |  |  |  |  |  |  |  |  |  |  |  |  |  |  |  |  |  |  |  |  |  |  |  |  |  |  |  |  |  |  |  |  |  |  |  |  |  |  |  |  |  |  |  |  |  |  |
| Leu (L) | 20 | 12.3% |  |  |  |  |  |  |  |  |  |  |  |  |  |  |  |  |  |  |  |  |  |  |  |  |  |  |  |  |  |  |  |  |  |  |  |  |  |  |  |  |  |  |  |  |  |  |  |  |  |  |  |  |  |  |  |  |  |  |  |  |  |  |  |  |  |
| Lys (K) | 9 | 5.6% |  |  |  |  |  |  |  |  |  |  |  |  |  |  |  |  |  |  |  |  |  |  |  |  |  |  |  |  |  |  |  |  |  |  |  |  |  |  |  |  |  |  |  |  |  |  |  |  |  |  |  |  |  |  |  |  |  |  |  |  |  |  |  |  |  |
| Met (M) | 3 | 1.9% |  |  |  |  |  |  |  |  |  |  |  |  |  |  |  |  |  |  |  |  |  |  |  |  |  |  |  |  |  |  |  |  |  |  |  |  |  |  |  |  |  |  |  |  |  |  |  |  |  |  |  |  |  |  |  |  |  |  |  |  |  |  |  |  |  |
| Phe (F) | 5 | 3.1% |  |  |  |  |  |  |  |  |  |  |  |  |  |  |  |  |  |  |  |  |  |  |  |  |  |  |  |  |  |  |  |  |  |  |  |  |  |  |  |  |  |  |  |  |  |  |  |  |  |  |  |  |  |  |  |  |  |  |  |  |  |  |  |  |  |
| Pro (P) | 11 | 6.8% |  |  |  |  |  |  |  |  |  |  |  |  |  |  |  |  |  |  |  |  |  |  |  |  |  |  |  |  |  |  |  |  |  |  |  |  |  |  |  |  |  |  |  |  |  |  |  |  |  |  |  |  |  |  |  |  |  |  |  |  |  |  |  |  |  |
| Ser (S) | 6 | 3.7% |  |  |  |  |  |  |  |  |  |  |  |  |  |  |  |  |  |  |  |  |  |  |  |  |  |  |  |  |  |  |  |  |  |  |  |  |  |  |  |  |  |  |  |  |  |  |  |  |  |  |  |  |  |  |  |  |  |  |  |  |  |  |  |  |  |
| Thr (T) | 10 | 6.2% |  |  |  |  |  |  |  |  |  |  |  |  |  |  |  |  |  |  |  |  |  |  |  |  |  |  |  |  |  |  |  |  |  |  |  |  |  |  |  |  |  |  |  |  |  |  |  |  |  |  |  |  |  |  |  |  |  |  |  |  |  |  |  |  |  |
| Trp (W) | 2 | 1.2% |  |  |  |  |  |  |  |  |  |  |  |  |  |  |  |  |  |  |  |  |  |  |  |  |  |  |  |  |  |  |  |  |  |  |  |  |  |  |  |  |  |  |  |  |  |  |  |  |  |  |  |  |  |  |  |  |  |  |  |  |  |  |  |  |  |
| Tyr (Y) | 3 | 1.9% |  |  |  |  |  |  |  |  |  |  |  |  |  |  |  |  |  |  |  |  |  |  |  |  |  |  |  |  |  |  |  |  |  |  |  |  |  |  |  |  |  |  |  |  |  |  |  |  |  |  |  |  |  |  |  |  |  |  |  |  |  |  |  |  |  |
| Val (V) | 10 | 6.2% |  |  |  |  |  |  |  |  |  |  |  |  |  |  |  |  |  |  |  |  |  |  |  |  |  |  |  |  |  |  |  |  |  |  |  |  |  |  |  |  |  |  |  |  |  |  |  |  |  |  |  |  |  |  |  |  |  |  |  |  |  |  |  |  |  |
| Pyl (O) | 0 | 0.0% |  |  |  |  |  |  |  |  |  |  |  |  |  |  |  |  |  |  |  |  |  |  |  |  |  |  |  |  |  |  |  |  |  |  |  |  |  |  |  |  |  |  |  |  |  |  |  |  |  |  |  |  |  |  |  |  |  |  |  |  |  |  |  |  |  |
| Sec (U) | 0 | 0.0% |  |  |  |  |  |  |  |  |  |  |  |  |  |  |  |  |  |  |  |  |  |  |  |  |  |  |  |  |  |  |  |  |  |  |  |  |  |  |  |  |  |  |  |  |  |  |  |  |  |  |  |  |  |  |  |  |  |  |  |  |  |  |  |  |  |
| hypothetical protein EI555_013315, partial<br>[Monodon monoceros] GenBank: TKC51741.1<br>Narwhal<br>ll llavgltllg<br>cpqalhwgppq dpnfnetlvv sgewflagma snqpkllked<br>kdarllvhri qvtpralqlh<br>lhrkvngacv pitmmanktk rkfqyllena dqnrlfleev<br>dpksyvilcn hrekrekevv<br>vvnlsttpe aspdallft nycrnghihp tniikvtttg gcpplpq<br>From -<br>glaakkragss qqpgsvviaa skmvhsrgsa rqhgpggrvte<br>pplsppkpg ll llavgltllg<br>cpqalhwgppq dpnfnetlvv sgewflagma snqpkllked<br>kdarllvhri qvtpralqlh<br>lhrkvngacv pitmmanktk rkfqyllena dqnrlfleev<br>dpksyvilcn hrekrekevv<br>vvnlsttpe aspdallft nycrnghihp tniikvtttg<br>gcpplpq prt saqlatdsap<br>plpqicphhp pplsphpll ftgvsprhrr trrsrltqga<br>lqlphplaaa stlleacpac | Number of amino acids: 179<br>Theoretical pI: 9.04<br>Molecular weight: 20173.55<br>Amino acid composition:<br>CSV format<br>Ala (A)<br>11<br>6.1%<br>Arg (R)<br>10<br>5.6%<br>Asn (N)<br>12<br>6.7%<br>Asp (D)<br>6<br>3.4%<br>Cys (C)<br>5<br>2.8%<br>Gln (Q)<br>8<br>4.5%<br>Glu (E)<br>10<br>5.6%<br>Gly (G)<br>9<br>5.0%<br>His (H)<br>7<br>3.9%<br>Ile (I)<br>6<br>3.4%<br>Leu (L)<br>28<br>15.6%<br>Lys (K)<br>11<br>6.1%<br>Met (M)<br>3<br>1.7%<br>Phe (F)<br>5<br>2.8%<br>Pro (P)<br>13<br>7.3%<br>Ser (S)<br>5<br>2.8%<br>Thr (T)<br>11<br>6.1%<br>Trp (W)<br>2<br>1.1%<br>Tyr (Y)<br>3<br>1.7%<br>Val (V)<br>14<br>7.8%<br>Pyl (O)<br>0<br>0.0%<br>Sec (U)<br>0<br>0.0% | Ala (A) | 11 | 6.1% | Arg (R) | 10 | 5.6% | Asn (N) | 12 | 6.7% | Asp (D) | 6 | 3.4% | Cys (C) | 5 | 2.8% | Gln (Q) | 8 | 4.5% | Glu (E) | 10 | 5.6% | Gly (G) | 9 | 5.0% | His (H) | 7 | 3.9% | Ile (I) | 6 | 3.4% | Leu (L) | 28 | 15.6% | Lys (K) | 11 | 6.1% | Met (M) | 3 | 1.7% | Phe (F) | 5 | 2.8% | Pro (P) | 13 | 7.3% | Ser (S) | 5 | 2.8% | Thr (T) | 11 | 6.1% | Trp (W) | 2 | 1.1% | Tyr (Y) | 3 | 1.7% | Val (V) | 14 | 7.8% | Pyl (O) | 0 | 0.0% | Sec (U) | 0 | 0.0% |
| Ala (A) | 11 | 6.1% |  |  |  |  |  |  |  |  |  |  |  |  |  |  |  |  |  |  |  |  |  |  |  |  |  |  |  |  |  |  |  |  |  |  |  |  |  |  |  |  |  |  |  |  |  |  |  |  |  |  |  |  |  |  |  |  |  |  |  |  |  |  |  |  |  |
| Arg (R) | 10 | 5.6% |  |  |  |  |  |  |  |  |  |  |  |  |  |  |  |  |  |  |  |  |  |  |  |  |  |  |  |  |  |  |  |  |  |  |  |  |  |  |  |  |  |  |  |  |  |  |  |  |  |  |  |  |  |  |  |  |  |  |  |  |  |  |  |  |  |
| Asn (N) | 12 | 6.7% |  |  |  |  |  |  |  |  |  |  |  |  |  |  |  |  |  |  |  |  |  |  |  |  |  |  |  |  |  |  |  |  |  |  |  |  |  |  |  |  |  |  |  |  |  |  |  |  |  |  |  |  |  |  |  |  |  |  |  |  |  |  |  |  |  |
| Asp (D) | 6 | 3.4% |  |  |  |  |  |  |  |  |  |  |  |  |  |  |  |  |  |  |  |  |  |  |  |  |  |  |  |  |  |  |  |  |  |  |  |  |  |  |  |  |  |  |  |  |  |  |  |  |  |  |  |  |  |  |  |  |  |  |  |  |  |  |  |  |  |
| Cys (C) | 5 | 2.8% |  |  |  |  |  |  |  |  |  |  |  |  |  |  |  |  |  |  |  |  |  |  |  |  |  |  |  |  |  |  |  |  |  |  |  |  |  |  |  |  |  |  |  |  |  |  |  |  |  |  |  |  |  |  |  |  |  |  |  |  |  |  |  |  |  |
| Gln (Q) | 8 | 4.5% |  |  |  |  |  |  |  |  |  |  |  |  |  |  |  |  |  |  |  |  |  |  |  |  |  |  |  |  |  |  |  |  |  |  |  |  |  |  |  |  |  |  |  |  |  |  |  |  |  |  |  |  |  |  |  |  |  |  |  |  |  |  |  |  |  |
| Glu (E) | 10 | 5.6% |  |  |  |  |  |  |  |  |  |  |  |  |  |  |  |  |  |  |  |  |  |  |  |  |  |  |  |  |  |  |  |  |  |  |  |  |  |  |  |  |  |  |  |  |  |  |  |  |  |  |  |  |  |  |  |  |  |  |  |  |  |  |  |  |  |
| Gly (G) | 9 | 5.0% |  |  |  |  |  |  |  |  |  |  |  |  |  |  |  |  |  |  |  |  |  |  |  |  |  |  |  |  |  |  |  |  |  |  |  |  |  |  |  |  |  |  |  |  |  |  |  |  |  |  |  |  |  |  |  |  |  |  |  |  |  |  |  |  |  |
| His (H) | 7 | 3.9% |  |  |  |  |  |  |  |  |  |  |  |  |  |  |  |  |  |  |  |  |  |  |  |  |  |  |  |  |  |  |  |  |  |  |  |  |  |  |  |  |  |  |  |  |  |  |  |  |  |  |  |  |  |  |  |  |  |  |  |  |  |  |  |  |  |
| Ile (I) | 6 | 3.4% |  |  |  |  |  |  |  |  |  |  |  |  |  |  |  |  |  |  |  |  |  |  |  |  |  |  |  |  |  |  |  |  |  |  |  |  |  |  |  |  |  |  |  |  |  |  |  |  |  |  |  |  |  |  |  |  |  |  |  |  |  |  |  |  |  |
| Leu (L) | 28 | 15.6% |  |  |  |  |  |  |  |  |  |  |  |  |  |  |  |  |  |  |  |  |  |  |  |  |  |  |  |  |  |  |  |  |  |  |  |  |  |  |  |  |  |  |  |  |  |  |  |  |  |  |  |  |  |  |  |  |  |  |  |  |  |  |  |  |  |
| Lys (K) | 11 | 6.1% |  |  |  |  |  |  |  |  |  |  |  |  |  |  |  |  |  |  |  |  |  |  |  |  |  |  |  |  |  |  |  |  |  |  |  |  |  |  |  |  |  |  |  |  |  |  |  |  |  |  |  |  |  |  |  |  |  |  |  |  |  |  |  |  |  |
| Met (M) | 3 | 1.7% |  |  |  |  |  |  |  |  |  |  |  |  |  |  |  |  |  |  |  |  |  |  |  |  |  |  |  |  |  |  |  |  |  |  |  |  |  |  |  |  |  |  |  |  |  |  |  |  |  |  |  |  |  |  |  |  |  |  |  |  |  |  |  |  |  |
| Phe (F) | 5 | 2.8% |  |  |  |  |  |  |  |  |  |  |  |  |  |  |  |  |  |  |  |  |  |  |  |  |  |  |  |  |  |  |  |  |  |  |  |  |  |  |  |  |  |  |  |  |  |  |  |  |  |  |  |  |  |  |  |  |  |  |  |  |  |  |  |  |  |
| Pro (P) | 13 | 7.3% |  |  |  |  |  |  |  |  |  |  |  |  |  |  |  |  |  |  |  |  |  |  |  |  |  |  |  |  |  |  |  |  |  |  |  |  |  |  |  |  |  |  |  |  |  |  |  |  |  |  |  |  |  |  |  |  |  |  |  |  |  |  |  |  |  |
| Ser (S) | 5 | 2.8% |  |  |  |  |  |  |  |  |  |  |  |  |  |  |  |  |  |  |  |  |  |  |  |  |  |  |  |  |  |  |  |  |  |  |  |  |  |  |  |  |  |  |  |  |  |  |  |  |  |  |  |  |  |  |  |  |  |  |  |  |  |  |  |  |  |
| Thr (T) | 11 | 6.1% |  |  |  |  |  |  |  |  |  |  |  |  |  |  |  |  |  |  |  |  |  |  |  |  |  |  |  |  |  |  |  |  |  |  |  |  |  |  |  |  |  |  |  |  |  |  |  |  |  |  |  |  |  |  |  |  |  |  |  |  |  |  |  |  |  |
| Trp (W) | 2 | 1.1% |  |  |  |  |  |  |  |  |  |  |  |  |  |  |  |  |  |  |  |  |  |  |  |  |  |  |  |  |  |  |  |  |  |  |  |  |  |  |  |  |  |  |  |  |  |  |  |  |  |  |  |  |  |  |  |  |  |  |  |  |  |  |  |  |  |
| Tyr (Y) | 3 | 1.7% |  |  |  |  |  |  |  |  |  |  |  |  |  |  |  |  |  |  |  |  |  |  |  |  |  |  |  |  |  |  |  |  |  |  |  |  |  |  |  |  |  |  |  |  |  |  |  |  |  |  |  |  |  |  |  |  |  |  |  |  |  |  |  |  |  |
| Val (V) | 14 | 7.8% |  |  |  |  |  |  |  |  |  |  |  |  |  |  |  |  |  |  |  |  |  |  |  |  |  |  |  |  |  |  |  |  |  |  |  |  |  |  |  |  |  |  |  |  |  |  |  |  |  |  |  |  |  |  |  |  |  |  |  |  |  |  |  |  |  |
| Pyl (O) | 0 | 0.0% |  |  |  |  |  |  |  |  |  |  |  |  |  |  |  |  |  |  |  |  |  |  |  |  |  |  |  |  |  |  |  |  |  |  |  |  |  |  |  |  |  |  |  |  |  |  |  |  |  |  |  |  |  |  |  |  |  |  |  |  |  |  |  |  |  |
| Sec (U) | 0 | 0.0% |  |  |  |  |  |  |  |  |  |  |  |  |  |  |  |  |  |  |  |  |  |  |  |  |  |  |  |  |  |  |  |  |  |  |  |  |  |  |  |  |  |  |  |  |  |  |  |  |  |  |  |  |  |  |  |  |  |  |  |  |  |  |  |  |  |

|  |  |  |  |  |  |  |  |  |  |  |  |  |  |  |  |  |  |  |  |  |  |  |  |  |  |  |  |  |  |  |  |  |  |  |  |  |  |  |  |  |  |  |  |  |  |  |  |  |  |  |  |  |  |  |  |  |  |  |  |  |  |  |  |  |  |  |  |
| --- | --- | --- | --- | --- | --- | --- | --- | --- | --- | --- | --- | --- | --- | --- | --- | --- | --- | --- | --- | --- | --- | --- | --- | --- | --- | --- | --- | --- | --- | --- | --- | --- | --- | --- | --- | --- | --- | --- | --- | --- | --- | --- | --- | --- | --- | --- | --- | --- | --- | --- | --- | --- | --- | --- | --- | --- | --- | --- | --- | --- | --- | --- | --- | --- | --- | --- | --- |
| algdlsnts vasasmvtgs cpltphrpr llgtgdgdp<br>gdtklaldr prgleklqv<br>hqdtvsakl gdgqgalssg qespeavswa rlghvaprlt<br>saqdegelaa gslsrkgpp<br>astagarphp ssqaaltpps lpappgspr tptpearvp<br>thvrcpcllc alilgqhqp<br>vhptdalgps gpghglgeqg afgtvpeaq srlggkdsr<br>ndalggptqr npntgkqthh<br>rlpqgasrpl rsawgqalk lgevspppat phfthsfpr<br>ggqlslhspr qrgrsgprp<br>svpcgrlqqr psrvgiwiga cpvwasaalp vpcgrlqrhp<br>srlgvcsгал sdaaalysws<br>qglwdptqdp gp |  |  |  |  |  |  |  |  |  |  |  |  |  |  |  |  |  |  |  |  |  |  |  |  |  |  |  |  |  |  |  |  |  |  |  |  |  |  |  |  |  |  |  |  |  |  |  |  |  |  |  |  |  |  |  |  |  |  |  |  |  |  |  |  |  |  |  |
| Other lipocalins |  |  |  |  |  |  |  |  |  |  |  |  |  |  |  |  |  |  |  |  |  |  |  |  |  |  |  |  |  |  |  |  |  |  |  |  |  |  |  |  |  |  |  |  |  |  |  |  |  |  |  |  |  |  |  |  |  |  |  |  |  |  |  |  |  |  |  |
| epididymal-specific lipocalin-9 isoform X1<br>[Equus caballus]<br>NCBI Reference Sequence: XP_014591634.3<br>ma~~llll~~slgv slvsa qqldl rtivrrnyi arvsgdwfsv<br>smasddmkri eengdlrvfi<br>qkiksledgg lkfyfqlll gqcvevpmvc ekmekngect<br>isyegenrvl laetdyrvya<br>tfhllnlrng tqtqlalyg ripdlspfl krfekvckky glgpqnivsl<br>idkdpclk | Number of amino acids: 163<br>Theoretical pI: 8.28<br>Molecular weight: 18762.74<br>Amino acid composition:<br>CSV format<br>Ala (A)<br>5<br>3.1%<br>Arg (R)<br>11<br>6.7%<br>Asn (N)<br>8<br>4.9%<br>Asp (D)<br>10<br>6.1%<br>Cys (C)<br>5<br>3.1%<br>Gln (Q)<br>8<br>4.9%<br>Glu (E)<br>11<br>6.7%<br>Gly (G)<br>11<br>6.7%<br>His (H)<br>1<br>0.6%<br>Ile (I)<br>9<br>5.5%<br>Leu (L)<br>21<br>12.9%<br>Lys (K)<br>12<br>7.4%<br>Met (M)<br>4<br>2.5%<br>Phe (F)<br>7<br>4.3%<br>Pro (P)<br>5<br>3.1%<br>Ser (S)<br>9<br>5.5%<br>Thr (T)<br>6<br>3.7%<br>Trp (W)<br>1<br>0.6%<br>Tyr (Y)<br>7<br>4.3%<br>Val (V)<br>12<br>7.4%<br>Pyl (O)<br>0<br>0.0%<br>Sec (U)<br>0<br>0.0% | Ala (A) | 5 | 3.1% | Arg (R) | 11 | 6.7% | Asn (N) | 8 | 4.9% | Asp (D) | 10 | 6.1% | Cys (C) | 5 | 3.1% | Gln (Q) | 8 | 4.9% | Glu (E) | 11 | 6.7% | Gly (G) | 11 | 6.7% | His (H) | 1 | 0.6% | Ile (I) | 9 | 5.5% | Leu (L) | 21 | 12.9% | Lys (K) | 12 | 7.4% | Met (M) | 4 | 2.5% | Phe (F) | 7 | 4.3% | Pro (P) | 5 | 3.1% | Ser (S) | 9 | 5.5% | Thr (T) | 6 | 3.7% | Trp (W) | 1 | 0.6% | Tyr (Y) | 7 | 4.3% | Val (V) | 12 | 7.4% | Pyl (O) | 0 | 0.0% | Sec (U) | 0 | 0.0% |
| Ala (A) | 5 | 3.1% |  |  |  |  |  |  |  |  |  |  |  |  |  |  |  |  |  |  |  |  |  |  |  |  |  |  |  |  |  |  |  |  |  |  |  |  |  |  |  |  |  |  |  |  |  |  |  |  |  |  |  |  |  |  |  |  |  |  |  |  |  |  |  |  |  |
| Arg (R) | 11 | 6.7% |  |  |  |  |  |  |  |  |  |  |  |  |  |  |  |  |  |  |  |  |  |  |  |  |  |  |  |  |  |  |  |  |  |  |  |  |  |  |  |  |  |  |  |  |  |  |  |  |  |  |  |  |  |  |  |  |  |  |  |  |  |  |  |  |  |
| Asn (N) | 8 | 4.9% |  |  |  |  |  |  |  |  |  |  |  |  |  |  |  |  |  |  |  |  |  |  |  |  |  |  |  |  |  |  |  |  |  |  |  |  |  |  |  |  |  |  |  |  |  |  |  |  |  |  |  |  |  |  |  |  |  |  |  |  |  |  |  |  |  |
| Asp (D) | 10 | 6.1% |  |  |  |  |  |  |  |  |  |  |  |  |  |  |  |  |  |  |  |  |  |  |  |  |  |  |  |  |  |  |  |  |  |  |  |  |  |  |  |  |  |  |  |  |  |  |  |  |  |  |  |  |  |  |  |  |  |  |  |  |  |  |  |  |  |
| Cys (C) | 5 | 3.1% |  |  |  |  |  |  |  |  |  |  |  |  |  |  |  |  |  |  |  |  |  |  |  |  |  |  |  |  |  |  |  |  |  |  |  |  |  |  |  |  |  |  |  |  |  |  |  |  |  |  |  |  |  |  |  |  |  |  |  |  |  |  |  |  |  |
| Gln (Q) | 8 | 4.9% |  |  |  |  |  |  |  |  |  |  |  |  |  |  |  |  |  |  |  |  |  |  |  |  |  |  |  |  |  |  |  |  |  |  |  |  |  |  |  |  |  |  |  |  |  |  |  |  |  |  |  |  |  |  |  |  |  |  |  |  |  |  |  |  |  |
| Glu (E) | 11 | 6.7% |  |  |  |  |  |  |  |  |  |  |  |  |  |  |  |  |  |  |  |  |  |  |  |  |  |  |  |  |  |  |  |  |  |  |  |  |  |  |  |  |  |  |  |  |  |  |  |  |  |  |  |  |  |  |  |  |  |  |  |  |  |  |  |  |  |
| Gly (G) | 11 | 6.7% |  |  |  |  |  |  |  |  |  |  |  |  |  |  |  |  |  |  |  |  |  |  |  |  |  |  |  |  |  |  |  |  |  |  |  |  |  |  |  |  |  |  |  |  |  |  |  |  |  |  |  |  |  |  |  |  |  |  |  |  |  |  |  |  |  |
| His (H) | 1 | 0.6% |  |  |  |  |  |  |  |  |  |  |  |  |  |  |  |  |  |  |  |  |  |  |  |  |  |  |  |  |  |  |  |  |  |  |  |  |  |  |  |  |  |  |  |  |  |  |  |  |  |  |  |  |  |  |  |  |  |  |  |  |  |  |  |  |  |
| Ile (I) | 9 | 5.5% |  |  |  |  |  |  |  |  |  |  |  |  |  |  |  |  |  |  |  |  |  |  |  |  |  |  |  |  |  |  |  |  |  |  |  |  |  |  |  |  |  |  |  |  |  |  |  |  |  |  |  |  |  |  |  |  |  |  |  |  |  |  |  |  |  |
| Leu (L) | 21 | 12.9% |  |  |  |  |  |  |  |  |  |  |  |  |  |  |  |  |  |  |  |  |  |  |  |  |  |  |  |  |  |  |  |  |  |  |  |  |  |  |  |  |  |  |  |  |  |  |  |  |  |  |  |  |  |  |  |  |  |  |  |  |  |  |  |  |  |
| Lys (K) | 12 | 7.4% |  |  |  |  |  |  |  |  |  |  |  |  |  |  |  |  |  |  |  |  |  |  |  |  |  |  |  |  |  |  |  |  |  |  |  |  |  |  |  |  |  |  |  |  |  |  |  |  |  |  |  |  |  |  |  |  |  |  |  |  |  |  |  |  |  |
| Met (M) | 4 | 2.5% |  |  |  |  |  |  |  |  |  |  |  |  |  |  |  |  |  |  |  |  |  |  |  |  |  |  |  |  |  |  |  |  |  |  |  |  |  |  |  |  |  |  |  |  |  |  |  |  |  |  |  |  |  |  |  |  |  |  |  |  |  |  |  |  |  |
| Phe (F) | 7 | 4.3% |  |  |  |  |  |  |  |  |  |  |  |  |  |  |  |  |  |  |  |  |  |  |  |  |  |  |  |  |  |  |  |  |  |  |  |  |  |  |  |  |  |  |  |  |  |  |  |  |  |  |  |  |  |  |  |  |  |  |  |  |  |  |  |  |  |
| Pro (P) | 5 | 3.1% |  |  |  |  |  |  |  |  |  |  |  |  |  |  |  |  |  |  |  |  |  |  |  |  |  |  |  |  |  |  |  |  |  |  |  |  |  |  |  |  |  |  |  |  |  |  |  |  |  |  |  |  |  |  |  |  |  |  |  |  |  |  |  |  |  |
| Ser (S) | 9 | 5.5% |  |  |  |  |  |  |  |  |  |  |  |  |  |  |  |  |  |  |  |  |  |  |  |  |  |  |  |  |  |  |  |  |  |  |  |  |  |  |  |  |  |  |  |  |  |  |  |  |  |  |  |  |  |  |  |  |  |  |  |  |  |  |  |  |  |
| Thr (T) | 6 | 3.7% |  |  |  |  |  |  |  |  |  |  |  |  |  |  |  |  |  |  |  |  |  |  |  |  |  |  |  |  |  |  |  |  |  |  |  |  |  |  |  |  |  |  |  |  |  |  |  |  |  |  |  |  |  |  |  |  |  |  |  |  |  |  |  |  |  |
| Trp (W) | 1 | 0.6% |  |  |  |  |  |  |  |  |  |  |  |  |  |  |  |  |  |  |  |  |  |  |  |  |  |  |  |  |  |  |  |  |  |  |  |  |  |  |  |  |  |  |  |  |  |  |  |  |  |  |  |  |  |  |  |  |  |  |  |  |  |  |  |  |  |
| Tyr (Y) | 7 | 4.3% |  |  |  |  |  |  |  |  |  |  |  |  |  |  |  |  |  |  |  |  |  |  |  |  |  |  |  |  |  |  |  |  |  |  |  |  |  |  |  |  |  |  |  |  |  |  |  |  |  |  |  |  |  |  |  |  |  |  |  |  |  |  |  |  |  |
| Val (V) | 12 | 7.4% |  |  |  |  |  |  |  |  |  |  |  |  |  |  |  |  |  |  |  |  |  |  |  |  |  |  |  |  |  |  |  |  |  |  |  |  |  |  |  |  |  |  |  |  |  |  |  |  |  |  |  |  |  |  |  |  |  |  |  |  |  |  |  |  |  |
| Pyl (O) | 0 | 0.0% |  |  |  |  |  |  |  |  |  |  |  |  |  |  |  |  |  |  |  |  |  |  |  |  |  |  |  |  |  |  |  |  |  |  |  |  |  |  |  |  |  |  |  |  |  |  |  |  |  |  |  |  |  |  |  |  |  |  |  |  |  |  |  |  |  |
| Sec (U) | 0 | 0.0% |  |  |  |  |  |  |  |  |  |  |  |  |  |  |  |  |  |  |  |  |  |  |  |  |  |  |  |  |  |  |  |  |  |  |  |  |  |  |  |  |  |  |  |  |  |  |  |  |  |  |  |  |  |  |  |  |  |  |  |  |  |  |  |  |  |

|  |  |  |  |  |  |  |  |  |  |  |  |  |  |  |  |  |  |  |  |  |  |  |  |  |  |  |  |  |  |  |  |  |  |  |  |  |  |  |  |  |  |  |  |  |  |  |  |  |  |  |  |  |  |  |  |  |  |  |  |  |  |  |  |  |  |  |  |
| --- | --- | --- | --- | --- | --- | --- | --- | --- | --- | --- | --- | --- | --- | --- | --- | --- | --- | --- | --- | --- | --- | --- | --- | --- | --- | --- | --- | --- | --- | --- | --- | --- | --- | --- | --- | --- | --- | --- | --- | --- | --- | --- | --- | --- | --- | --- | --- | --- | --- | --- | --- | --- | --- | --- | --- | --- | --- | --- | --- | --- | --- | --- | --- | --- | --- | --- | --- |
| retinol binding protein precursor [Equus caballus]<br>GenBank: AAC48461.1<br>mewvwalvvl aalgsaga er dcrvssfrvk enfdkarfsg twya<br>makkdp eglflqdniv<br>aefsvdeyqg msatakgrvr llnnwdvcad mvgftfdded<br>pakfkmkywg vasflqkgnd<br>dhwiidtdyd tyavqyscr lnlldgtcads ysfvfardpn<br>gfppevqriv rrrqeelcla<br>rqyrlishng ycdgksdrnl l | Number of amino acids: 162<br>Theoretical pI: 4.95<br>Molecular weight: 18704.96<br>Amino acid composition:<br>CSV fo<br>Ala (A)<br>13<br>8.0%<br>Arg (R)<br>12<br>7.4%<br>Asn (N)<br>7<br>4.3%<br>Asp (D)<br>17<br>10.5%<br>Cys (C)<br>5<br>3.1%<br>Gln (Q)<br>6<br>3.7%<br>Glu (E)<br>9<br>5.6%<br>Gly (G)<br>9<br>5.6%<br>His (H)<br>1<br>0.6%<br>Ile (I)<br>4<br>2.5%<br>Leu (L)<br>10<br>6.2%<br>Lys (K)<br>9<br>5.6%<br>Met (M)<br>4<br>2.5%<br>Phe (F)<br>11<br>6.8%<br>Pro (P)<br>5<br>3.1%<br>Ser (S)<br>9<br>5.6%<br>Thr (T)<br>8<br>4.9%<br>Trp (W)<br>4<br>2.5%<br>Tyr (Y)<br>7<br>4.3%<br>Val (V)<br>12<br>7.4%<br>Pyl (O)<br>0<br>0.0%<br>Sec (U)<br>0<br>0.0% | Ala (A) | 13 | 8.0% | Arg (R) | 12 | 7.4% | Asn (N) | 7 | 4.3% | Asp (D) | 17 | 10.5% | Cys (C) | 5 | 3.1% | Gln (Q) | 6 | 3.7% | Glu (E) | 9 | 5.6% | Gly (G) | 9 | 5.6% | His (H) | 1 | 0.6% | Ile (I) | 4 | 2.5% | Leu (L) | 10 | 6.2% | Lys (K) | 9 | 5.6% | Met (M) | 4 | 2.5% | Phe (F) | 11 | 6.8% | Pro (P) | 5 | 3.1% | Ser (S) | 9 | 5.6% | Thr (T) | 8 | 4.9% | Trp (W) | 4 | 2.5% | Tyr (Y) | 7 | 4.3% | Val (V) | 12 | 7.4% | Pyl (O) | 0 | 0.0% | Sec (U) | 0 | 0.0% |
| Ala (A) | 13 | 8.0% |  |  |  |  |  |  |  |  |  |  |  |  |  |  |  |  |  |  |  |  |  |  |  |  |  |  |  |  |  |  |  |  |  |  |  |  |  |  |  |  |  |  |  |  |  |  |  |  |  |  |  |  |  |  |  |  |  |  |  |  |  |  |  |  |  |
| Arg (R) | 12 | 7.4% |  |  |  |  |  |  |  |  |  |  |  |  |  |  |  |  |  |  |  |  |  |  |  |  |  |  |  |  |  |  |  |  |  |  |  |  |  |  |  |  |  |  |  |  |  |  |  |  |  |  |  |  |  |  |  |  |  |  |  |  |  |  |  |  |  |
| Asn (N) | 7 | 4.3% |  |  |  |  |  |  |  |  |  |  |  |  |  |  |  |  |  |  |  |  |  |  |  |  |  |  |  |  |  |  |  |  |  |  |  |  |  |  |  |  |  |  |  |  |  |  |  |  |  |  |  |  |  |  |  |  |  |  |  |  |  |  |  |  |  |
| Asp (D) | 17 | 10.5% |  |  |  |  |  |  |  |  |  |  |  |  |  |  |  |  |  |  |  |  |  |  |  |  |  |  |  |  |  |  |  |  |  |  |  |  |  |  |  |  |  |  |  |  |  |  |  |  |  |  |  |  |  |  |  |  |  |  |  |  |  |  |  |  |  |
| Cys (C) | 5 | 3.1% |  |  |  |  |  |  |  |  |  |  |  |  |  |  |  |  |  |  |  |  |  |  |  |  |  |  |  |  |  |  |  |  |  |  |  |  |  |  |  |  |  |  |  |  |  |  |  |  |  |  |  |  |  |  |  |  |  |  |  |  |  |  |  |  |  |
| Gln (Q) | 6 | 3.7% |  |  |  |  |  |  |  |  |  |  |  |  |  |  |  |  |  |  |  |  |  |  |  |  |  |  |  |  |  |  |  |  |  |  |  |  |  |  |  |  |  |  |  |  |  |  |  |  |  |  |  |  |  |  |  |  |  |  |  |  |  |  |  |  |  |
| Glu (E) | 9 | 5.6% |  |  |  |  |  |  |  |  |  |  |  |  |  |  |  |  |  |  |  |  |  |  |  |  |  |  |  |  |  |  |  |  |  |  |  |  |  |  |  |  |  |  |  |  |  |  |  |  |  |  |  |  |  |  |  |  |  |  |  |  |  |  |  |  |  |
| Gly (G) | 9 | 5.6% |  |  |  |  |  |  |  |  |  |  |  |  |  |  |  |  |  |  |  |  |  |  |  |  |  |  |  |  |  |  |  |  |  |  |  |  |  |  |  |  |  |  |  |  |  |  |  |  |  |  |  |  |  |  |  |  |  |  |  |  |  |  |  |  |  |
| His (H) | 1 | 0.6% |  |  |  |  |  |  |  |  |  |  |  |  |  |  |  |  |  |  |  |  |  |  |  |  |  |  |  |  |  |  |  |  |  |  |  |  |  |  |  |  |  |  |  |  |  |  |  |  |  |  |  |  |  |  |  |  |  |  |  |  |  |  |  |  |  |
| Ile (I) | 4 | 2.5% |  |  |  |  |  |  |  |  |  |  |  |  |  |  |  |  |  |  |  |  |  |  |  |  |  |  |  |  |  |  |  |  |  |  |  |  |  |  |  |  |  |  |  |  |  |  |  |  |  |  |  |  |  |  |  |  |  |  |  |  |  |  |  |  |  |
| Leu (L) | 10 | 6.2% |  |  |  |  |  |  |  |  |  |  |  |  |  |  |  |  |  |  |  |  |  |  |  |  |  |  |  |  |  |  |  |  |  |  |  |  |  |  |  |  |  |  |  |  |  |  |  |  |  |  |  |  |  |  |  |  |  |  |  |  |  |  |  |  |  |
| Lys (K) | 9 | 5.6% |  |  |  |  |  |  |  |  |  |  |  |  |  |  |  |  |  |  |  |  |  |  |  |  |  |  |  |  |  |  |  |  |  |  |  |  |  |  |  |  |  |  |  |  |  |  |  |  |  |  |  |  |  |  |  |  |  |  |  |  |  |  |  |  |  |
| Met (M) | 4 | 2.5% |  |  |  |  |  |  |  |  |  |  |  |  |  |  |  |  |  |  |  |  |  |  |  |  |  |  |  |  |  |  |  |  |  |  |  |  |  |  |  |  |  |  |  |  |  |  |  |  |  |  |  |  |  |  |  |  |  |  |  |  |  |  |  |  |  |
| Phe (F) | 11 | 6.8% |  |  |  |  |  |  |  |  |  |  |  |  |  |  |  |  |  |  |  |  |  |  |  |  |  |  |  |  |  |  |  |  |  |  |  |  |  |  |  |  |  |  |  |  |  |  |  |  |  |  |  |  |  |  |  |  |  |  |  |  |  |  |  |  |  |
| Pro (P) | 5 | 3.1% |  |  |  |  |  |  |  |  |  |  |  |  |  |  |  |  |  |  |  |  |  |  |  |  |  |  |  |  |  |  |  |  |  |  |  |  |  |  |  |  |  |  |  |  |  |  |  |  |  |  |  |  |  |  |  |  |  |  |  |  |  |  |  |  |  |
| Ser (S) | 9 | 5.6% |  |  |  |  |  |  |  |  |  |  |  |  |  |  |  |  |  |  |  |  |  |  |  |  |  |  |  |  |  |  |  |  |  |  |  |  |  |  |  |  |  |  |  |  |  |  |  |  |  |  |  |  |  |  |  |  |  |  |  |  |  |  |  |  |  |
| Thr (T) | 8 | 4.9% |  |  |  |  |  |  |  |  |  |  |  |  |  |  |  |  |  |  |  |  |  |  |  |  |  |  |  |  |  |  |  |  |  |  |  |  |  |  |  |  |  |  |  |  |  |  |  |  |  |  |  |  |  |  |  |  |  |  |  |  |  |  |  |  |  |
| Trp (W) | 4 | 2.5% |  |  |  |  |  |  |  |  |  |  |  |  |  |  |  |  |  |  |  |  |  |  |  |  |  |  |  |  |  |  |  |  |  |  |  |  |  |  |  |  |  |  |  |  |  |  |  |  |  |  |  |  |  |  |  |  |  |  |  |  |  |  |  |  |  |
| Tyr (Y) | 7 | 4.3% |  |  |  |  |  |  |  |  |  |  |  |  |  |  |  |  |  |  |  |  |  |  |  |  |  |  |  |  |  |  |  |  |  |  |  |  |  |  |  |  |  |  |  |  |  |  |  |  |  |  |  |  |  |  |  |  |  |  |  |  |  |  |  |  |  |
| Val (V) | 12 | 7.4% |  |  |  |  |  |  |  |  |  |  |  |  |  |  |  |  |  |  |  |  |  |  |  |  |  |  |  |  |  |  |  |  |  |  |  |  |  |  |  |  |  |  |  |  |  |  |  |  |  |  |  |  |  |  |  |  |  |  |  |  |  |  |  |  |  |
| Pyl (O) | 0 | 0.0% |  |  |  |  |  |  |  |  |  |  |  |  |  |  |  |  |  |  |  |  |  |  |  |  |  |  |  |  |  |  |  |  |  |  |  |  |  |  |  |  |  |  |  |  |  |  |  |  |  |  |  |  |  |  |  |  |  |  |  |  |  |  |  |  |  |
| Sec (U) | 0 | 0.0% |  |  |  |  |  |  |  |  |  |  |  |  |  |  |  |  |  |  |  |  |  |  |  |  |  |  |  |  |  |  |  |  |  |  |  |  |  |  |  |  |  |  |  |  |  |  |  |  |  |  |  |  |  |  |  |  |  |  |  |  |  |  |  |  |  |
| major urinary protein (Mup)-like precursor [Mus musculus]<br>NCBI Reference Sequence: NP_001074754.1<br>mkllllclgl ilvcvha eea ssmgrnfnve kingewytii<br>lasdkrakie ehgimrlfve<br>hihvlenslg fkhftvidee cseiflvadk tekageysvt<br>ydgfkkftvl ktdydnyimf<br>hlinemnget fqlmslygre pdlnsdikek fvkclceehgi<br>ireniidftk tnrclqare | Number of amino acids: 162<br>Theoretical pI: 5.09<br>Molecular weight: 18986.54<br>Amino acid composition:<br>CSV format<br>Ala (A)<br>6<br>3.7%<br>Arg (R)<br>7<br>4.3%<br>Asn (N)<br>10<br>6.2%<br>Asp (D)<br>9<br>5.6%<br>Cys (C)<br>3<br>1.9%<br>Gln (Q)<br>2<br>1.2%<br>Glu (E)<br>21<br>13.0%<br>Gly (G)<br>9<br>5.6%<br>His (H)<br>6<br>3.7%<br>Ile (I)<br>15<br>9.3%<br>Leu (L)<br>12<br>7.4%<br>Lys (K)<br>13<br>8.0%<br>Met (M)<br>5<br>3.1%<br>Phe (F)<br>11<br>6.8%<br>Pro (P)<br>1<br>0.6%<br>Ser (S)<br>8<br>4.9%<br>Thr (T)<br>9<br>5.6%<br>Trp (W)<br>1<br>0.6%<br>Tyr (Y)<br>6<br>3.7%<br>Val (V)<br>8<br>4.9%<br>Pyl (O)<br>0<br>0.0%<br>Sec (U)<br>0<br>0.0% | Ala (A) | 6 | 3.7% | Arg (R) | 7 | 4.3% | Asn (N) | 10 | 6.2% | Asp (D) | 9 | 5.6% | Cys (C) | 3 | 1.9% | Gln (Q) | 2 | 1.2% | Glu (E) | 21 | 13.0% | Gly (G) | 9 | 5.6% | His (H) | 6 | 3.7% | Ile (I) | 15 | 9.3% | Leu (L) | 12 | 7.4% | Lys (K) | 13 | 8.0% | Met (M) | 5 | 3.1% | Phe (F) | 11 | 6.8% | Pro (P) | 1 | 0.6% | Ser (S) | 8 | 4.9% | Thr (T) | 9 | 5.6% | Trp (W) | 1 | 0.6% | Tyr (Y) | 6 | 3.7% | Val (V) | 8 | 4.9% | Pyl (O) | 0 | 0.0% | Sec (U) | 0 | 0.0% |
| Ala (A) | 6 | 3.7% |  |  |  |  |  |  |  |  |  |  |  |  |  |  |  |  |  |  |  |  |  |  |  |  |  |  |  |  |  |  |  |  |  |  |  |  |  |  |  |  |  |  |  |  |  |  |  |  |  |  |  |  |  |  |  |  |  |  |  |  |  |  |  |  |  |
| Arg (R) | 7 | 4.3% |  |  |  |  |  |  |  |  |  |  |  |  |  |  |  |  |  |  |  |  |  |  |  |  |  |  |  |  |  |  |  |  |  |  |  |  |  |  |  |  |  |  |  |  |  |  |  |  |  |  |  |  |  |  |  |  |  |  |  |  |  |  |  |  |  |
| Asn (N) | 10 | 6.2% |  |  |  |  |  |  |  |  |  |  |  |  |  |  |  |  |  |  |  |  |  |  |  |  |  |  |  |  |  |  |  |  |  |  |  |  |  |  |  |  |  |  |  |  |  |  |  |  |  |  |  |  |  |  |  |  |  |  |  |  |  |  |  |  |  |
| Asp (D) | 9 | 5.6% |  |  |  |  |  |  |  |  |  |  |  |  |  |  |  |  |  |  |  |  |  |  |  |  |  |  |  |  |  |  |  |  |  |  |  |  |  |  |  |  |  |  |  |  |  |  |  |  |  |  |  |  |  |  |  |  |  |  |  |  |  |  |  |  |  |
| Cys (C) | 3 | 1.9% |  |  |  |  |  |  |  |  |  |  |  |  |  |  |  |  |  |  |  |  |  |  |  |  |  |  |  |  |  |  |  |  |  |  |  |  |  |  |  |  |  |  |  |  |  |  |  |  |  |  |  |  |  |  |  |  |  |  |  |  |  |  |  |  |  |
| Gln (Q) | 2 | 1.2% |  |  |  |  |  |  |  |  |  |  |  |  |  |  |  |  |  |  |  |  |  |  |  |  |  |  |  |  |  |  |  |  |  |  |  |  |  |  |  |  |  |  |  |  |  |  |  |  |  |  |  |  |  |  |  |  |  |  |  |  |  |  |  |  |  |
| Glu (E) | 21 | 13.0% |  |  |  |  |  |  |  |  |  |  |  |  |  |  |  |  |  |  |  |  |  |  |  |  |  |  |  |  |  |  |  |  |  |  |  |  |  |  |  |  |  |  |  |  |  |  |  |  |  |  |  |  |  |  |  |  |  |  |  |  |  |  |  |  |  |
| Gly (G) | 9 | 5.6% |  |  |  |  |  |  |  |  |  |  |  |  |  |  |  |  |  |  |  |  |  |  |  |  |  |  |  |  |  |  |  |  |  |  |  |  |  |  |  |  |  |  |  |  |  |  |  |  |  |  |  |  |  |  |  |  |  |  |  |  |  |  |  |  |  |
| His (H) | 6 | 3.7% |  |  |  |  |  |  |  |  |  |  |  |  |  |  |  |  |  |  |  |  |  |  |  |  |  |  |  |  |  |  |  |  |  |  |  |  |  |  |  |  |  |  |  |  |  |  |  |  |  |  |  |  |  |  |  |  |  |  |  |  |  |  |  |  |  |
| Ile (I) | 15 | 9.3% |  |  |  |  |  |  |  |  |  |  |  |  |  |  |  |  |  |  |  |  |  |  |  |  |  |  |  |  |  |  |  |  |  |  |  |  |  |  |  |  |  |  |  |  |  |  |  |  |  |  |  |  |  |  |  |  |  |  |  |  |  |  |  |  |  |
| Leu (L) | 12 | 7.4% |  |  |  |  |  |  |  |  |  |  |  |  |  |  |  |  |  |  |  |  |  |  |  |  |  |  |  |  |  |  |  |  |  |  |  |  |  |  |  |  |  |  |  |  |  |  |  |  |  |  |  |  |  |  |  |  |  |  |  |  |  |  |  |  |  |
| Lys (K) | 13 | 8.0% |  |  |  |  |  |  |  |  |  |  |  |  |  |  |  |  |  |  |  |  |  |  |  |  |  |  |  |  |  |  |  |  |  |  |  |  |  |  |  |  |  |  |  |  |  |  |  |  |  |  |  |  |  |  |  |  |  |  |  |  |  |  |  |  |  |
| Met (M) | 5 | 3.1% |  |  |  |  |  |  |  |  |  |  |  |  |  |  |  |  |  |  |  |  |  |  |  |  |  |  |  |  |  |  |  |  |  |  |  |  |  |  |  |  |  |  |  |  |  |  |  |  |  |  |  |  |  |  |  |  |  |  |  |  |  |  |  |  |  |
| Phe (F) | 11 | 6.8% |  |  |  |  |  |  |  |  |  |  |  |  |  |  |  |  |  |  |  |  |  |  |  |  |  |  |  |  |  |  |  |  |  |  |  |  |  |  |  |  |  |  |  |  |  |  |  |  |  |  |  |  |  |  |  |  |  |  |  |  |  |  |  |  |  |
| Pro (P) | 1 | 0.6% |  |  |  |  |  |  |  |  |  |  |  |  |  |  |  |  |  |  |  |  |  |  |  |  |  |  |  |  |  |  |  |  |  |  |  |  |  |  |  |  |  |  |  |  |  |  |  |  |  |  |  |  |  |  |  |  |  |  |  |  |  |  |  |  |  |
| Ser (S) | 8 | 4.9% |  |  |  |  |  |  |  |  |  |  |  |  |  |  |  |  |  |  |  |  |  |  |  |  |  |  |  |  |  |  |  |  |  |  |  |  |  |  |  |  |  |  |  |  |  |  |  |  |  |  |  |  |  |  |  |  |  |  |  |  |  |  |  |  |  |
| Thr (T) | 9 | 5.6% |  |  |  |  |  |  |  |  |  |  |  |  |  |  |  |  |  |  |  |  |  |  |  |  |  |  |  |  |  |  |  |  |  |  |  |  |  |  |  |  |  |  |  |  |  |  |  |  |  |  |  |  |  |  |  |  |  |  |  |  |  |  |  |  |  |
| Trp (W) | 1 | 0.6% |  |  |  |  |  |  |  |  |  |  |  |  |  |  |  |  |  |  |  |  |  |  |  |  |  |  |  |  |  |  |  |  |  |  |  |  |  |  |  |  |  |  |  |  |  |  |  |  |  |  |  |  |  |  |  |  |  |  |  |  |  |  |  |  |  |
| Tyr (Y) | 6 | 3.7% |  |  |  |  |  |  |  |  |  |  |  |  |  |  |  |  |  |  |  |  |  |  |  |  |  |  |  |  |  |  |  |  |  |  |  |  |  |  |  |  |  |  |  |  |  |  |  |  |  |  |  |  |  |  |  |  |  |  |  |  |  |  |  |  |  |
| Val (V) | 8 | 4.9% |  |  |  |  |  |  |  |  |  |  |  |  |  |  |  |  |  |  |  |  |  |  |  |  |  |  |  |  |  |  |  |  |  |  |  |  |  |  |  |  |  |  |  |  |  |  |  |  |  |  |  |  |  |  |  |  |  |  |  |  |  |  |  |  |  |
| Pyl (O) | 0 | 0.0% |  |  |  |  |  |  |  |  |  |  |  |  |  |  |  |  |  |  |  |  |  |  |  |  |  |  |  |  |  |  |  |  |  |  |  |  |  |  |  |  |  |  |  |  |  |  |  |  |  |  |  |  |  |  |  |  |  |  |  |  |  |  |  |  |  |
| Sec (U) | 0 | 0.0% |  |  |  |  |  |  |  |  |  |  |  |  |  |  |  |  |  |  |  |  |  |  |  |  |  |  |  |  |  |  |  |  |  |  |  |  |  |  |  |  |  |  |  |  |  |  |  |  |  |  |  |  |  |  |  |  |  |  |  |  |  |  |  |  |  |

**epididymal-specific lipocalin-9 precursor [Mus musculus]**

NCBI Reference Sequence: NP\_084235.1

mvllllvlglv lsataq fnl htavrrdynl arisgtwyld
siasdnmtri eengdlrlfi
rnikllnngs lqdfhfmlq gecvavtmvc ektknngfs
vayegknkv lletdysmyi
ifymqnikng tktqvlalyg rsilldkthq refenicnly
gldsqniidm tkkdfcfl

Number of amino acids: 161
Theoretical pI: 6.34
Molecular weight: 18766.53

Amino acid composition: CSV format

|  |  |  |
| --- | --- | --- |
| Ala (A) | 6 | 3.7% |
| Arg (R) | 8 | 5.0% |
| Asn (N) | 15 | 9.3% |
| Asp (D) | 10 | 6.2% |
| Cys (C) | 4 | 2.5% |
| Gln (Q) | 6 | 3.7% |
| Glu (E) | 9 | 5.6% |
| Gly (G) | 9 | 5.6% |
| His (H) | 3 | 1.9% |
| Ile (I) | 12 | 7.5% |
| Leu (L) | 19 | 11.8% |
| Lys (K) | 10 | 6.2% |
| Met (M) | 6 | 3.7% |
| Phe (F) | 10 | 6.2% |
| Pro (P) | 0 | 0.0% |
| Ser (S) | 8 | 5.0% |
| Thr (T) | 10 | 6.2% |
| Trp (W) | 1 | 0.6% |
| Tyr (Y) | 8 | 5.0% |
| Val (V) | 7 | 4.3% |
| Pyl (O) | 0 | 0.0% |
| Sec (U) | 0 | 0.0% |

**epididymal-specific lipocalin-9 isoform X1 [Equus caballus]**

NCBI Reference Sequence: XP\_014591635.3

malllslglv slvsa qqldl rtivrrnyni arvsgdwfsv
smasddmkri eengdlrvfi
qkiksledgg lkfyfqllll gqcvevpmvc ekmekngect
isyegenrvl laetdyrvya
tfhllnlrng tqtqvlalyg ripdlspsfl krfekvckky glgpqnivsl
idkdpclk

Number of amino acids: 163
Theoretical pI: 8.28
Molecular weight: 18762.74

Amino acid composition: CSV format

|  |  |  |
| --- | --- | --- |
| Ala (A) | 5 | 3.1% |
| Arg (R) | 11 | 6.7% |
| Asn (N) | 8 | 4.9% |
| Asp (D) | 10 | 6.1% |
| Cys (C) | 5 | 3.1% |
| Gln (Q) | 8 | 4.9% |
| Glu (E) | 11 | 6.7% |
| Gly (G) | 11 | 6.7% |
| His (H) | 1 | 0.6% |
| Ile (I) | 9 | 5.5% |
| Leu (L) | 21 | 12.9% |
| Lys (K) | 12 | 7.4% |
| Met (M) | 4 | 2.5% |
| Phe (F) | 7 | 4.3% |
| Pro (P) | 5 | 3.1% |
| Ser (S) | 9 | 5.5% |
| Thr (T) | 6 | 3.7% |
| Trp (W) | 1 | 0.6% |
| Tyr (Y) | 7 | 4.3% |
| Val (V) | 12 | 7.4% |
| Pyl (O) | 0 | 0.0% |
| Sec (U) | 0 | 0.0% |

**major allergen Equ c 1 precursor [Equus caballus]**

NCBI Reference Sequence: NP\_001075966.1

mkllllclgl ilvca qqeen sdvairnfdi skisgewysi
flasdvkeki eengsmrvfv
dviraldnss lyaeyqtkvn gectefpmvf dkteedgvys
lnydgynvfr isefendehi
ilylvnfdkd rpfqlfefya repdvspeik eefvkivqkr
givkeniidl tkidrcfqlr

gngvaqa

**Number of amino acids:** 165

**Theoretical pI:** 4.51

**Molecular weight:** 19499.85

**Amino acid composition:** CSV format

|  |  |  |
| --- | --- | --- |
| Ala (A) | 5 | 3.0% |
| Arg (R) | 9 | 5.5% |
| Asn (N) | 10 | 6.1% |
| Asp (D) | 14 | 8.5% |
| Cys (C) | 2 | 1.2% |
| Gln (Q) | 6 | 3.6% |
| Glu (E) | 20 | 12.1% |
| Gly (G) | 6 | 3.6% |
| His (H) | 1 | 0.6% |
| Ile (I) | 15 | 9.1% |
| Leu (L) | 9 | 5.5% |
| Lys (K) | 11 | 6.7% |
| Met (M) | 2 | 1.2% |
| Phe (F) | 13 | 7.9% |
| Pro (P) | 4 | 2.4% |
| Ser (S) | 11 | 6.7% |
| Thr (T) | 4 | 2.4% |
| Trp (W) | 1 | 0.6% |
| Tyr (Y) | 8 | 4.8% |
| Val (V) | 14 | 8.5% |
| Pyl (O) | 0 | 0.0% |
| Sec (U) | 0 | 0.0% |

**trichosurin [Trichosurus vulpecula]**

GenBank: AAA84739.1

mkllllsmgl alvcg lqpec srseedlsde kerkweqlsr
hwhtvvlass drslieeegp
frnfiqnitv esgnlngffl trkngqcipl yltafkteea rqfklnyygt
ndvyygsskp
neyakfifyn yhdgkvnvva nlfgrtpnls neikkrfeed
fmnrgfrren ildisevdhc

**Number of amino acids:** 165

**Theoretical pI:** 5.65

**Molecular weight:** 19480.66

**Amino acid composition:** CSV format

|  |  |  |
| --- | --- | --- |
| Ala (A) | 5 | 3.0% |
| Arg (R) | 12 | 7.3% |
| Asn (N) | 15 | 9.1% |
| Asp (D) | 8 | 4.8% |
| Cys (C) | 3 | 1.8% |
| Gln (Q) | 5 | 3.0% |
| Glu (E) | 18 | 10.9% |
| Gly (G) | 9 | 5.5% |
| His (H) | 4 | 2.4% |
| Ile (I) | 8 | 4.8% |
| Leu (L) | 13 | 7.9% |
| Lys (K) | 10 | 6.1% |
| Met (M) | 1 | 0.6% |
| Phe (F) | 12 | 7.3% |
| Pro (P) | 5 | 3.0% |
| Ser (S) | 12 | 7.3% |
| Thr (T) | 7 | 4.2% |
| Trp (W) | 2 | 1.2% |
| Tyr (Y) | 8 | 4.8% |
| Val (V) | 8 | 4.8% |
| Pyl (O) | 0 | 0.0% |
| Sec (U) | 0 | 0.0% |

**beta-lactoglobulin precursor [Bos taurus]**

NCBI Reference Sequence: NP\_776354.2

mkclllalal tcgaq alivt qtmkgldiqk vagtwyslam
aasdisllda qsaplrvyve
elkptpegdl eillqkweng ecaqkkiaa ktkipavfki
dalnenkvlv ldt dykkyll
fcmensaepe qslacqclvr tpevddeale kfdkalkalp
mhirlsfnp t qleeqchi

Number of amino acids: 163

Theoretical pI: 4.83

Molecular weight: 18352.29

**Amino acid composition:** CSV format

|  |  |  |
| --- | --- | --- |
| Ala (A) | 16 | 9.8% |
| Arg (R) | 3 | 1.8% |
| Asn (N) | 5 | 3.1% |
| Asp (D) | 10 | 6.1% |
| Cys (C) | 5 | 3.1% |
| Gln (Q) | 9 | 5.5% |
| Glu (E) | 16 | 9.8% |
| Gly (G) | 4 | 2.5% |
| His (H) | 2 | 1.2% |
| Ile (I) | 10 | 6.1% |
| Leu (L) | 22 | 13.5% |
| Lys (K) | 15 | 9.2% |
| Met (M) | 4 | 2.5% |
| Phe (F) | 4 | 2.5% |
| Pro (P) | 8 | 4.9% |
| Ser (S) | 7 | 4.3% |
| Thr (T) | 8 | 4.9% |
| Trp (W) | 2 | 1.2% |
| Tyr (Y) | 4 | 2.5% |
| Val (V) | 9 | 5.5% |
| Pyl (O) | 0 | 0.0% |
| Sec (U) | 0 | 0.0% |

**Uterocalin [Camelus dromedarius]**

GenBank: KAB1280376.1

This has a deletion of about ten amino acids around position 100 when aligned with Equus uterocalin and others. Therefore not used.

msllllatgl tllgspqt lh qgpqdsnfne tlvs gdwfsa
alsnrprll qegadaqlfi
hsiqvtpral qlhlhrkvqf qppppappst ftdggqkrif
leevdpksyv ifcthhkeqg
ketvvvtlfs rtpkvtqdtl lifknycksh gihktniinv
tkagtlpps q kdqkilvnpr

spsasltl

**Figure S1. Heatmap and Principal Component Analysis of amino acid content to compare perissodactyl uterocalins, uterocalin-like proteins from non-perissodactyls, and a selection of unrelated lipocalins.** Identical analysis to that shown in figure 4 of the main text with additional samples of lipocalins. **Perissodactyl uterocalins** - (horse (*Equus*); rhino (*Diceros* and *Ceratotherium*); tapir (*Tapirus*)). **Uterocalin-like proteins** from other members of the Ferungulata - camel (*Camelus*); pig (*Sus*); cat (*Leopardus*); true seal (*Halichoerus*); sea lion (*Zalophus*); whale (*Megaptera*)), here split into those from Artiodactyla and Carnivora. **Unrelated lipocalins** - epididymis-specific lipocalin 9 (LCN9) from mouse, human and horse, to illustrate the amino acid balance in other proteins associated with reproductive tissues that are sometimes named as uterocalin in the databases; trichosurin, the milk protein from the marsupial *Trichosurus vulpecula* (brush-tailed possum), mouse urinary protein (MUP6); horse plasma retinol (vitamin A) binding protein; horse major skin allergen (Equ c 1); bovine milk beta lactoglobulin (*Bos* milk BLG). Note that the perissodactyl uterocalins group coherently in the bottom left of the heatmap. The 2-D scores plot again emphasises the distinctiveness of the perissodactyl uterocalins. In the VIP (Variable Importance in Projection) analysis variables with high scores are more influential in predicting sample classifications and are often considered key discriminative features – here the essential amino acids leucine (low), lysine (high), and phenylalanine (high) stand out. So also does alanine, a non-essential, and unusually low in perissodactyl uterocalins (see also in the heatmap). Alanine is synthesised endogenously in many tissues by mammals through several metabolic pathways (1) so its depletion in these uterocalins would have no deleterious implications for embryos, and it is frequently substituted with some other amino acids (2).

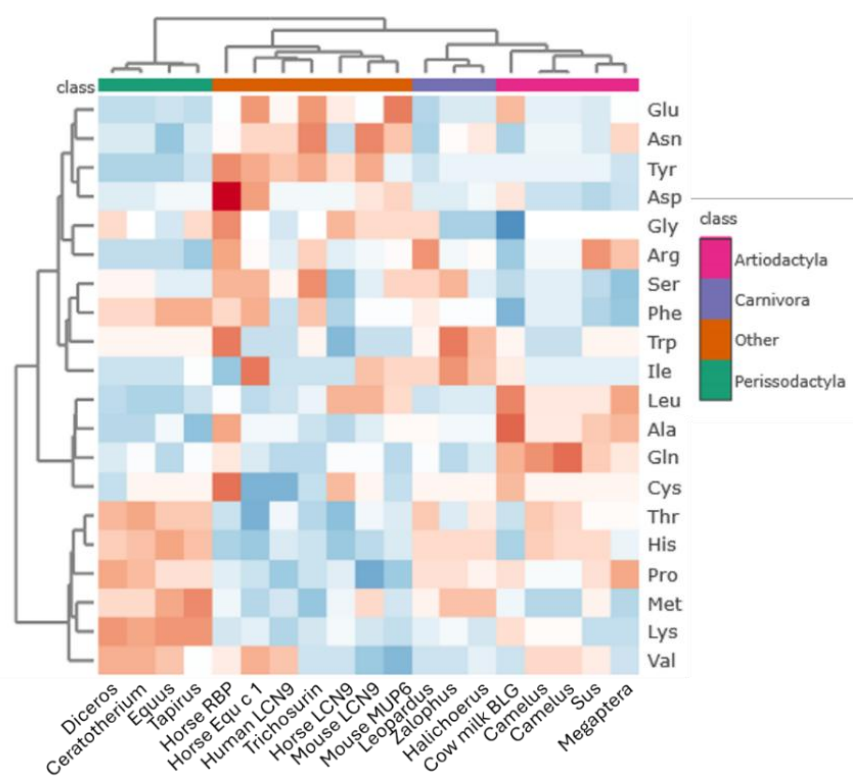

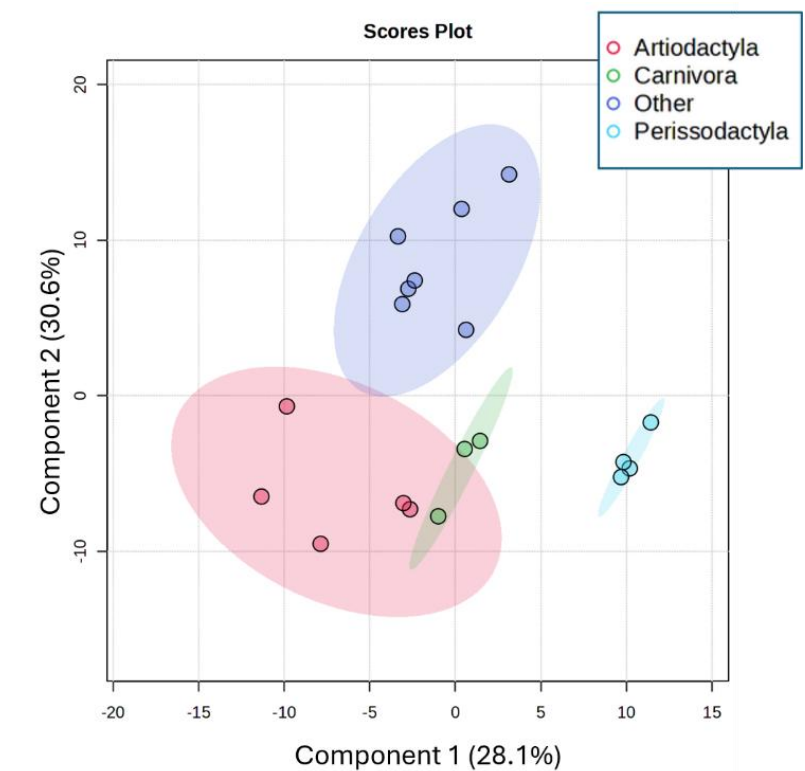

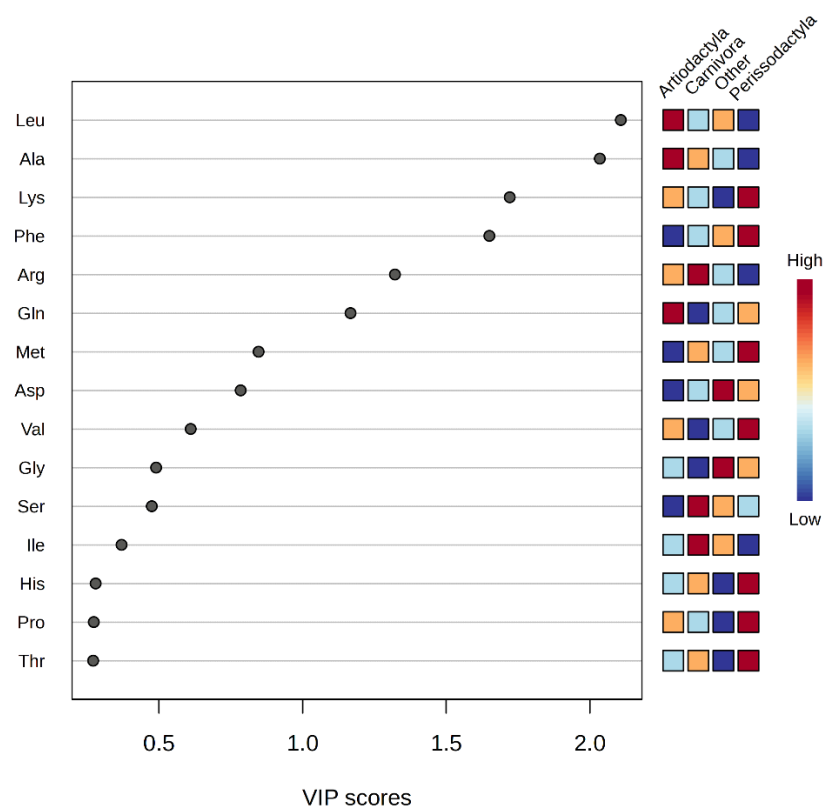

**Table S2. A simple diagnostic for “true” horse, rhinoceros and tapir uterocalins? These have high predicted isoelectric points (pI) combined with high lysine/leucine content ratios.**

The amino acid content and pI values were predicted by ProtParam (Table S1). The lysine/leucine ratios are based on the number of each amino acid in the mature secreted proteins (with secretory signal peptide omitted) and edited if necessary to about 162 amino acids in length (see Table S1).

All the proteins that are similar to the perissodactyl forms all have similarly high pI values, whether from perissodactyls or not. But only the perissodactyl uterocalins have lysine/leucine values above 2, whereas the uterocalin-like proteins from other Ferungulates (here - camels, pig, cat, sea lion, true seals, whale) have values close to unity or below.

Those listed under “other lipocalins” are proteins belonging to the lipocalin family but not related to the uterocalins or uterocalin-like proteins and are included as samples of the wider family.

| Species | Protein<br>(current database<br>naming) | Isoelectric<br>point (pI)<br>(theoretical) | Lysine/leucine ratio<br>(number in protein) |
| --- | --- | --- | --- |
| **Perissodactyl uterocalins** |  |  |  |
| Equus caballus<br>Horse | Uterocalin | 9.72 | 2.5 |
| Equus asinus<br>Donkey | Uterocalin isoform X2 | 9.72 | 2.5 |
| Equus przewalskii<br>Przewalskii's horse or takhi | Uterocalin | 9.72 | 2.5 |
| Equus quagga<br>Quagga (zebra) | Uterocalin isoform X2 | 9.72 | 2.5 |
| Diceros bicornis minor<br>Southern black rhinoceros | Epididymal-specific<br>lipocalin-9 | 9.72 | 2.22 |
| Ceratotherium simum simum<br>Southern white rhinoceros | Epididymal-specific<br>lipocalin-9 | 9.83 | 2.38 |
| Tapirus indicus<br>Indian tapir | Similar to uterocalin | 9.62 | 2.0 |
| **Other uterocalin-like proteins** |  |  |  |
| Camelus ferus<br>Bactrian camel | Uterocalin isoform X2 | 9.48 | 0.87 |
| Camelus dromedarius<br>Dromedary camel | Uterocalin | 9.48 | 0.87 |
| Zalophus californianus<br>California sea lion | Uterocalin-like isoform<br>X1 | 9.39 | 1.0 |
| Sus scrofa<br>Domestic pig | Uterocalin P19<br>precursor | 10.33 | 0.6 |
| Phoca vitulina<br>Harbour seal | Uterocalin-like | 9.28 | 0.92 |
| Halichoerus grypus<br>Grey seal | Uterocalin-like | 9.14 | 0.91 |
| Leopardus geoffroyi<br>Geoffroy's cat | Uterocalin-like isoform X2 | 10.64 | 1.0 |
| Callorhinus ursinus<br>Northern fur seal | Uterocalin-like | 9.17 | 1.2 |
| Megaptera novaeangliae<br>Humpback whale | Uterocalin-like isoform 1-T1 | 9.51 | 0.45 |
| Eschrichtius robustus<br>Grey whale | Hypothetical protein J1605_017252 | 9.62 | 0.45 |
| Balaenoptera musculus<br>Pygmy blue whale | Uterocalin-like | 9.37 | 0.37 |
| Balaenoptera ricei<br>Rice's whale | Uterocalin-like | 9.18 | 0.45 |
| Monodon monoceros<br>Narwhal | Hypothetical protein | 9.04 | 0.39 |
| **Other lipocalins** |  |  |  |
| Equus caballus<br>Horse | Epididymal-specific lipocalin-9 isoform X1 | 8.28 | 0.57 |
| Equus caballus<br>Horse | Retinol binding protein | 4.95 | 0.9 |
| Mus musculus<br>House mouse | Major urinary protein (Mup)-like | 5.09 | 1.08 |
| Mus musculus<br>House mouse | Epididymal-specific lipocalin-9 | 6.34 | 0.53 |
| Equus caballus<br>Horse | Epididymal-specific lipocalin-9 isoform X1 | 8.28 | 0.57 |
| Equus caballus<br>Horse | Major allergen Equ c 1 | 4.51 | 1.22 |
| Trichosurus vulpecula<br>Brushtail possum | Trichosurin<br>Milk protein | 5.65 | 0.77 |
| Bos taurus<br>Cow | Beta-lactoglobulin<br>Milk protein | 4.83 | 0.68 |

### References

1. Kohlmeier M. Amino acids and nitrogen compounds. In: Kohlmeier M, editor. Nutrient Metabolism: Structures, Functions, and Genes. Second ed. Amsterdam, Boston: Academic Press; 2015. p. 266–477.
2. Dayhoff M, Schwartz R, Orcutt B. Atlas of Protein Sequence and Structure. Washington, USA: National Biomedical Research Foundation; 1979.
